# Bipartite invariance in mouse primary visual cortex

**DOI:** 10.1101/2023.03.15.532836

**Authors:** Zhiwei Ding, Dat T. Tran, Kayla Ponder, Zhuokun Ding, Rachel Froebe, Lydia Ntanavara, Paul G. Fahey, Erick Cobos, Luca Baroni, Maria Diamantaki, Eric Y. Wang, Andersen Chang, Stelios Papadopoulos, Jiakun Fu, Taliah Muhammad, Christos Papadopoulos, Santiago A. Cadena, Alexandros Evangelou, Konstantin Willeke, Fabio Anselmi, Sophia Sanborn, Jan Antolik, Emmanouil Froudarakis, Saumil Patel, Edgar Y. Walker, Jacob Reimer, Fabian H. Sinz, Alexander S. Ecker, Katrin Franke, Xaq Pitkow, Andreas S. Tolias

**Affiliations:** Center for Neuroscience and Artificial Intelligence, Baylor College of Medicine, Houston, TX, USA; Department of Neuroscience, Baylor College of Medicine, Houston, TX, USA; Department of Ophthalmology, Byers Eye Institute, Stanford University School of Medicine, Stanford, CA, US; Stanford Bio-X, Stanford University, Stanford, CA, USA; Wu Tsai Neurosciences Institute, Stanford University, Stanford, CA, USA; Charles University, Prague, Czech Republic; Institute of Molecular Biology & Biotechnology, Foundation of Research & Technology - Hellas, Heraklion, Crete, Greece; School of Medicine, University of Crete, Heraklion, Crete, Greece; Institute of Computer Science and Campus Institute Data Science, University of Göttingen, Göttingen, Germany; Institute for Bioinformatics and Medical Informatics, University of Tübingen, Tübingen, Germany; Department of Mathematics, Informatics and Geoscience, University of Trieste, Trieste, Italy; Department of Physiology and Biophysics, University of Washington, Seattle, WA, USA; Computational Neuroscience Center, University of Washington, Seattle, WA, USA; Max Planck Institute for Dynamics and Self-Organization, Göttingen, Germany; Department of Electrical and Computer Engineering, Rice University, Houston, TX, USA; Department of Computer Science, Rice University, Houston, TX, USA; Neuroscience Institute, Carnegie Mellon University, Pittsburgh, PA, USA; Department of Machine Learning, Carnegie Mellon University, Pittsburgh, PA, USA; Department of Electrical Engineering, Stanford University, Stanford, CA, USA

## Abstract

A primary goal of sensory systems is to extract robust and meaningful features that are invariant to variations in the sensory input. Characterizing these invariances at the neuronal level is crucial for understanding how the visual system supports generalization, but the high-dimensional nature of ecological stimuli poses major challenges. Consequently, our understanding of how the brain represents invariances has historically depended on a few examples, such as phase invariance to grating stimuli in V1 complex cells. Here, we leverage the *inception loop* paradigm —iterating between large-scale recordings, deep learning neuronal predictive models, and *in silico* experiments with *in vivo* verification—to characterize neuronal invariances in mouse V1. Using a neuronal predictive model, we synthesized Diverse Exciting Inputs (DEIs) that strongly drive target neurons while differing substantially in image space. These DEIs revealed a novel bipartite invariance: one portion of the receptive field encodes shift-invariant, high-frequency textures, while the other encodes a fixed, low-frequency spatial pattern. This subfield division aligned with object boundaries defined by spatial frequency differences in highly activating stimuli, suggesting bi-partite invariance contributes to segmentation. Our analysis of computational models and anatomical data from the MICrONS dataset revealed a hierarchical organization of excitatory neurons in mouse V1 Layers 2/3: We found that postsynaptic neurons exhibited greater invariance than their presynaptic inputs, while neurons with lower invariance formed more connections. These findings suggest a synaptic-level hierarchy that progressively increases neural invariance within the primary visual cortex. Intriguingly, similar high-low frequency bipartite patterns strongly activate certain units in artificial neural networks, suggesting that universal visual representations govern both biological and artificial systems, potentially aiding in the extraction of visual features from complex backgrounds.

## Introduction

A central challenge of visual perception is to infer stable, behaviorally relevant latent features in the world despite continual fluctuations in the raw sensory inputs. For example, to recognize a familiar face in a crowded environment, the brain must extract relevant features from patterns of light to consistently identify the person, despite variations such as viewing distance, 3D pose, scale, and illumination. While these variations are often considered “nuisance” variables, the brain must represent them because they play crucial roles in other tasks, such as navigating through the crowd to approach the familiar face.

To understand how brains effectively disentangle high-dimensional sensory inputs and robustly extract latent variables (DiCarlo and Cox, 2007; Karklin and Lewicki, 2009; Higgins et al., 2022), it is essential to identify the features to which neurons exhibit selectivity (i.e. features that evoke maximal response) and invariance (i.e. feature variations that preserve high response magnitude). Identifying neuronal invariances is extremely challenging because of the enormous search space of visual stimuli, the non-linear information processing in the brain, and the limited experimental time. As a result, most previous studies have been limited to parametric stimuli (e.g., gratings) or semantic categories (e.g., objects and faces) (Gross et al., 1972; Tsao et al., 2006; Yamins and DiCarlo, 2016; Cadieu et al., 2007; Sharpee et al., 2013; El-Shamayleh and Pasupathy, 2016) that are chosen based on strong assumptions about the structure of invariances in the brain. The classic example of this approach is Hubel & Wiesel’s complex cells in the primary visual cortex (V1) (Hubel and Wiesel, 1962), which are tuned to gratings of a preferred orientation but are invariant to spatial phase (in contrast to simple cells, which are selective to both orientation and spatial phase). So far, however, we know little about other types of invariances in early visual areas beyond these classes of parametric stimuli.

Here, we take a data-driven, systematic approach to study neuronal invariances, leveraging the previously introduced “inception loop” paradigm (Walker et al., 2019). Using large-scale calcium imaging data, we trained a deep neural network model to accurately predict mouse V1 neuronal responses to arbitrary, new natural images. This model enables rapid and high-throughput *in silico* experiments, revealing neuronal response properties with precision and efficiency unattainable through traditional *in vivo* methods.

Using the trained model as a “digital twin” of the visual cortex, we synthesized a set of stimuli for each individual neuron that elicit strong responses while being maximally different from each other—”Diverse Exciting Inputs” (DEIs, Fig. 1a). The variation among a neuron’s DEIs reveal the visual features that define its invariances. To validate these model-generated predictions, we closed the loop by presenting DEIs back to the animal while recording the activity of the same neurons *in vivo*. Our results confirm the model’s predictions, demonstrating that DEIs reliably evoke strong responses in their target neurons.

**Fig. 1.**
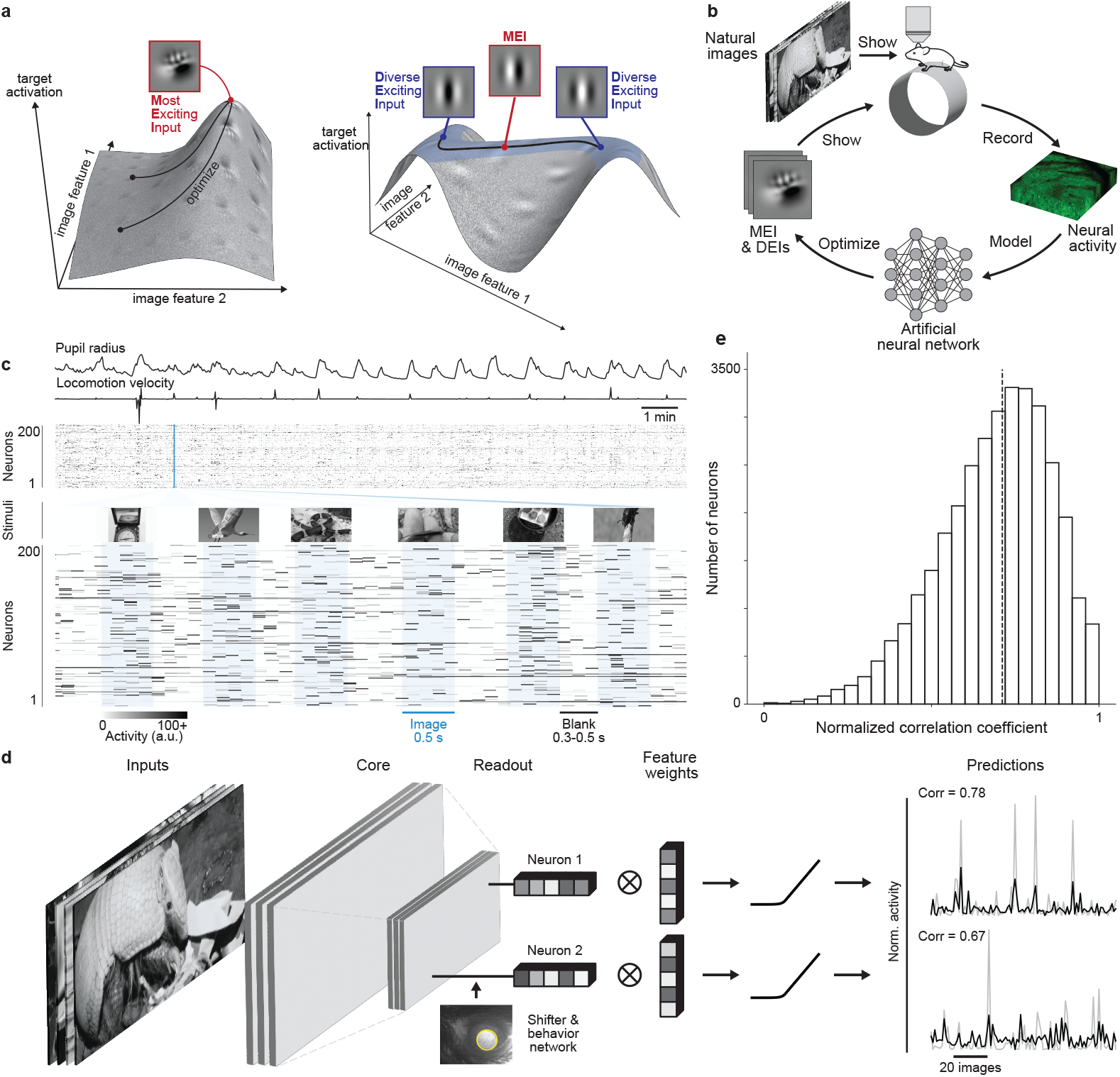
A deep neural network model accurately predicts mouse V1 responses to natural scenes. **a**, Schematic of the optimization of Most Exciting Inputs (MEI) and Diverse Exciting Inputs (DEIs). The vertical axes depict the activation of two model neurons as a function of two example image features. Left, neuron without obvious invariance; right, neuron with phase invariance to its optimal stimulus. Black curves illustrate optimization trajectories for MEI from different initializations (left) and for DEIs as perturbations starting from the MEI along the invariance ridge (right). **b**, Schematic of the inception loop paradigm. On day 1, we presented sequences of natural images and recorded *in vivo* neuronal activity using two-photon calcium imaging. Overnight, we trained an ensemble of CNNs to reproduce the measured neuronal responses and synthesized artificial stimuli for each target neuron *in silico*. On day 2, these stimuli were presented to the same neurons *in vivo* to compare measured and predicted responses. **c**, We presented 5,100 unique natural images to an awake mouse for 500 ms each, interleaved with gray screen gaps of random length between 300 and 500 ms. A subset of 100 images was repeated 10 times to estimate neuronal response reliability. Neuronal activity in V1 L2/3 was recorded at 8 Hz using wide-field two-photon microscopy. Behavioral traces including pupil dilation and locomotion velocity were also recorded. **d**, CNN model architecture schematic. The network is composed of a 3-layer convolutional core with a single-point readout predicting neuronal responses, a shifter network accounting for eye movements, and a behavioral modulator predicting neuron-specific adaptive gain (Sinz et al., 2018; Walker et al., 2019). Right: average responses (gray) to test images for two example neurons with corresponding model predictions (black). **e**, Performance of the model ensemble, measured as the normalized correlation coefficient between predicted and observed responses to the 100 held-out images (*CC*_*norm*_) (Schoppe et al., 2016). Data were pooled over 33,714 neurons from 14 mice (median=0.71, dashed line). Excessively noisy neurons (*CC*_*max*_ *<* 0.1) were excluded (0.2% of all neurons). Neurons with *CC*_*norm*_ outside [0, 1] were clipped (1.2%) for visualization.

The structure of the DEIs reveals a novel functional invariance in V1 neurons, which we refer to as “bipartite invariance”. These DEIs exhibit two distinct, non-overlapping sub-fields. In one subfield, the neuron’s response is selective for a particular texture while remaining robust across different spatial locations, resembling a response to different crops from an underlying texture. In contrast, the other subfield responds strongly only to a specific spatial pattern. Additionally, these neurons show a preference for bipartite stimuli defined by two regions of different spatial frequencies. The spatial division arising from bipartite invariance closely aligns with this bipartite frequency preference, suggesting that these V1 neurons may serve as specialized detectors for object boundaries, particularly those defined by abrupt changes in texture and spatial frequency.

Expanding our investigation, we adapted our methodology to analyze the functional connectomics MICrONs dataset (MICrONS Consortium et al., 2021) using a state-of-the-art foundation model (Wang et al., 2023). This approach reveals a novel hierarchical organization among excitatory neurons in mouse V1 Layer 2/3 that enhances single-neuron functional invariance. Specifically, we find that postsynaptic neurons exhibit greater invariance than their presynaptic partners, and neurons with lower invariance form more connections compared to those with higher invariance. Our findings collectively suggest a novel principle of receptive field organization in the mouse primary visual cortex, offering new insights into how the brain might extract visual features from complex backgrounds and advancing our understanding of circuit-level mechanisms underlying neuronal invariance.

## Results

### Diverse Exciting Inputs (DEIs) identify neuronal invariances

In this study, we employed inception loops (Walker et al., 2019; Bashivan et al., 2019), a closed-loop experimental paradigm, to investigate the invariances of single neurons in mouse V1 (Fig. 1b). An inception loop consists of four key steps:

1. Neuronal recordings: Conduct large-scale recordings with high-entropy natural stimuli to capture diverse neuronal responses (Fig. 1c).
2. Predictive modeling: Train a deep convolutional neural network (CNN) (Sinz et al., 2018; Walker et al., 2019; Franke et al., 2022) to predict the activity of neurons to arbitrary natural images. This model incorporates a non-linear core of convolutional layers shared across all recorded neurons, followed by neuron-specific linear readouts (Fig. 1d).
3. *In silico* experiments: Utilize the deep predictive model of the brain to systematically study the computations of the modeled neurons *in silico* and derive experimentally testable predictions.
4. *In vivo* verification: Validate these predictions through experiments in the actual brain.

We presented 5,100 unique natural images from ImageNet (ILSVRC2012) (Russakovsky et al., 2015) to awake, head-fixed mice while recording the activity of thousands of V1 Layer 2/3 (L2/3) excitatory neurons using two-photon calcium imaging (Fig. 1c). We used the recorded neuronal activity to train CNNs to predict the responses of these neurons to arbitrary natural images. The predictive model included a shifter network to compensate for eye movements and a modulator network for predicting an adaptive gain based on behavioral variables like running velocity and pupil size (Sinz et al., 2018; Walker et al., 2019) (Fig. 1d). We assessed model performance using a held-out set of natural images presented repeatedly during recording. For each neuron, we evaluated the predictive accuracy as the correlation between the recorded and the predicted responses to a novel set of stimuli that were not included in model training (Suppl. Fig. S1b). To account for noise in *in vivo* neuronal response, the correlation was normalized by neuronal self-reliability that served as an estimated upper bound on the model performance (Schoppe et al., 2016) (Suppl. Fig. S1a). The predictive models achieved a median normalized correlation coefficient of 0.71 (Schoppe et al., 2016) (Fig. 1e), comparable to state-of-the-art models of mouse V1 (Franke et al., 2022; Willeke et al., 2022; Lurz et al., 2020).

We adapted and extended recently-developed optimal stimulus synthesis frameworks to map both the selectivity (Walker et al., 2019; Bashivan et al., 2019) and invariance (Cadena et al., 2018) of individual neurons *in silico*. In our study, “selectivity” refers to the specific image features eliciting maximal neuronal responses, while “invariance” denotes image variations preserving high response magnitude. Expanding on our previous work (Walker et al., 2019), which identified a single Most Exciting Input (MEI) for each neuron, we now generate a set of 20 Diverse Exciting Inputs (DEIs; Fig. 1a) to characterize neuronal functional invariance. These DEIs, which we also refer to as “non-parametric DEIs” for ease of comparison in subsequent analyses, are defined as images that are maximally dissimilar in image pixel space but all strongly activate the target neuron. Specifically, during DEI synthesis, each stimulus was optimized to elicit at least 85% of the *in silico* activity evoked by the MEI. This threshold was chosen based on the Hubel & Wiesel model of complex cells (Hubel and Wiesel, 1962)—the most well-understood model of computing invariance—where perfect invariance was not achieved, yet Gabor patches in the preferred orientation and spatial frequency across all phases still highly activated these neurons (i.e. ≥85% of the MEI activation (Cadena et al., 2018)). This threshold represents a compromise where we maintain diversity through a regularization term to create DEIs that are highly diverse while accepting a small drop in activity (see our Methods and also (Cadena et al., 2018)). To generate DEIs, we added different white noises to the neuron’s MEI to create multiple initial images and then optimized these images to strongly drive the target neuron while maximizing their pixel-wise Euclidean distances from each other. This approach allows us to comprehensively map neuronal invariances while maintaining high activation levels, providing a more nuanced characterization of the neuronal response manifold than the single maximal response corresponding to the MEI.

Our DEI synthesis method successfully reproduced the expected functional invariances in simulated Hubel & Wiesel simple and complex cells (Fig. 2a). For simulated simple cells, DEIs resembled Gabor patches with identical orientation, spatial frequency, and phase (Fig. 2a, simulated simple), aligning with linear-nonlinear (LN) model predictions (Bussgang, 1952; Jones and Palmer, 1987a; Heeger, 1992). In contrast, simulated complex cell DEIs included Gabor patches with different phases, reflecting their known phase invariance (Fig. 2a, simulated complex).

**Fig. 2.**
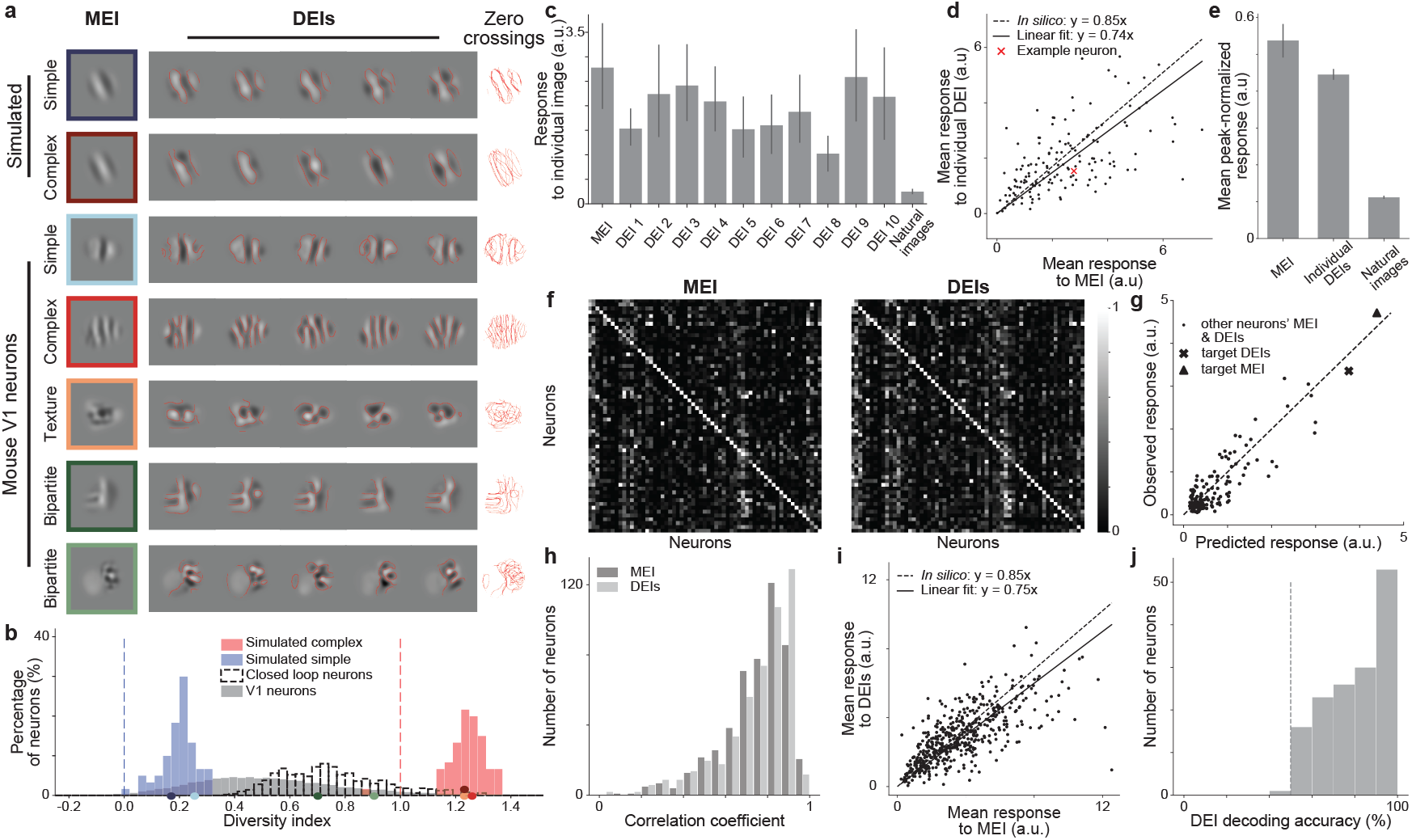
DEIs evoked strong and selective responses in target neurons while exhibiting population-decodable differences. **a**, Examples of MEI and DEIs for simulated simple and complex cells, and mouse V1 neurons. For each neuron, zero-crossing contours from individual DEIs (i.e.locations where the image intensity transitions from positive to negative values or vice versa) were overlaid. While DEIs strongly resembled the MEI, they exhibited complex features differing from the MEI and among themselves. Two types of novel invariances were observed: global shift invariance (texture) and partial shift invariance (bipartite). **b**, Diversity indices for 60 simulated complex cells (red), 60 simulated simple cells (blue), and 10,228 V1 neurons pooled from 14 mice (gray), including 500 tested in closed-loop experiments from 8 mice (unfilled). Diversity index is defined as the normalized average pairwise Euclidean distance in pixel space across the DEIs. Diversity indices for noiseless simple cells (0, blue dashed) and complex cells (1, red dashed) were shown for reference. V1 neuron diversity indices differed from simulated simple and complex cells (*P <* 10^−9^ for both, two-sided Welch’s *t*-test with 72.4 and 69.0 d.f., respectively). For closed-loop experiments, we randomly selected neurons with high diversity indices (see Methods for details). Example neurons from **a** were indicated on the x axis with the corresponding colors. Diversity indices below −0.25 were clipped to −0.25 for visualization (0.09% of all V1 neurons). **c**, Response of an example neuron to its MEI, 10 random DEIs, and 100 random full-field natural images. Only two out of the ten DEIs elicited responses lower than 85% of the MEI response (one after Benjamini/ Hochberg (BH) correction for multiple comparison). **d**, Comparison of mean responses to MEI and one random DEI per neuron. DEIs stimulated *in vivo* responses in target neurons close to the level predicted *in silico* relative to MEI (74 *±* 4% versus 85%) (two-sided Wilcoxon signed-rank test, *W* = 4902, *P* = 0.19) with only 274 out of 1490 DEIs (18.4%) showing responses lower than 85% of the corresponding MEI response (4.8% after BH correction) (*P <* 0.05, one-sided Welch’s *t*-test with 32.6 average d.f.). **e**, Mean peak-normalized *in vivo* responses to MEI and 10 random DEIs per neuron, and 100 random full-field natural images. For each neuron, responses were normalized by the largest mean response across all stimuli. Both MEI and individual DEIs elicited stronger responses in their target neurons compared to full-field natural images (*P <* 10^−9^ for both, two-sided Welch’s *t*-test with 148.7 and 1596.2 d.f., respectively). **c–e**, Responses to each MEI and individual DEI were averaged across 20 repeats; responses to each natural image were averaged across 10 repeats. Data were pooled over 149 neurons from 2 mice. Error bar represented 95% confidence interval. **f–i**, DEI responses were averaged across 20 different DEI with each presented once. **f**, Both MEI and DEIs activated neurons with high specificity. Confusion matrices showed responses of each neuron to MEI (left) and DEIs (right) for 61 neurons in one mouse. Responses of each neuron were normalized, with each row scaled so the maximum response across all images equaled 1. Neurons’ responses to their own MEI and DEIs (along the diagonal) were larger than those to other MEIs and DEIs, respectively (two-sided permutation test, *P <* 10^−4^ for both cases). **g**, Predicted versus observed responses of one example neuron to its own MEI and DEIs and 79 other neurons’ MEI and DEIs. **h**, Our model exhibited high predictive accuracy for both MEI and DEI responses (Pearson correlation coefficient between predicted and observed neuronal responses *r* = 0.74 and 0.75, respectively). **i**, DEIs stimulated *in vivo* responses close to the level predicted *in silico* relative to MEI (75 3% versus 85%) (two-sided Wilcoxon signed-rank test, *W* = 51360, *P* = 4.9 ×10^−4^), with only 9.6% of all neurons showing different responses between DEIs and 85% of MEI (1.2% after BH corrections) (*P <* 0.05, two-sided Welch’s *t*-test with 34.06 average d.f.). **h, i**, Data were pooled over 500 neurons from 8 mice. **j**, *In vivo* population responses in mouse V1 Layer 2/3 discriminated between a randomly selected pair of DEIs for each neuron. DEI identity in individual trials was decoded using a logistic regression classifier (see Method for details), with decoding accuracies across neurons (median = 80%) exceeded chance level (50%, dashed; one-sample *t*-test, *t* = 28.0, *P <* 10^−9^). Data were pooled over 149 neurons from 2 mice.

DEIs from mouse V1 neurons strongly resembled their corresponding MEIs while exhibiting specific variations indicative of different invariance types (Fig. 2a, mouse V1 neurons, see more examples in Suppl. Fig. S2). Some neurons produced nearly identical DEIs, suggesting a lack of invariance akin to simulated simple cells (Fig. 2a, mouse simple). A small subset of V1 neurons exhibited DEIs with varying phases while maintaining consistent orientation and spatial frequency, closely resembling the behavior of simulated complex cells (Fig. 2a, mouse complex).

Among neurons strongly activated by non-Gabor stimuli (Walker et al., 2019; Franke et al., 2022), some appeared to be stimulated strongly by random crops from an underlying texture pattern, demonstrating global shift invariance (Fig. 2a, mouse texture). We termed these “texture cells”, analogous to those observed in hidden layers of deep Artificial Neural Networks (ANNs) trained for object recognition (Zeiler and Fergus, 2014; Olah et al., 2017; Cadena et al., 2018). Intriguingly, many neurons exhibited a novel type of invariance that we denoted as “bipartite receptive field (RF) invariance” or equivalently, “bipartite invariance”, where one portion of their RF preferred a fixed spatial pattern, while the other responded robustly to different spatial translations of a specific texture image (Fig. 2a, mouse bipartite). In other words, the neuron’s response to the variable subfield remained strong when different crops of an underlying texture canvas were presented. We referred to these neurons as “bipartite cells”.

To quantify these phenomena, we computed a diversity index for each neuron using its DEIs. This index was formulated as the normalized average pairwise Euclidean distance in pixel space across the DEIs, with values of 0 and 1 indicating characteristics akin to classical simple and complex cells, respectively. The diversity indices of mouse V1 neurons spanned a continuous spectrum, with those of simulated simple and complex cells at the opposite extremes (Fig. 2b).

To assess whether the invariances captured by DEIs also appear in natural images, we screened over 41 millions crops to identify those that elicited DEI-like activation (Suppl. Fig. S3a). We found that only a small fraction (0.006%) of these images produced responses comparable to DEIs *in silico* (i.e. ≥85% of the MEI activation, Suppl. Fig. S3b) and only 37% of neurons yielded more than 20 such highly activating natural stimuli. Importantly, the highly activating natural crops closely resembled DEIs (Suppl. Fig. S3c, d), albeit with lower diversity (Suppl. Fig. S3e), highlighting the extreme lifetime sparsity of the neural code (Froudarakis et al., 2014). Collectively, these findings suggest that DEIs effectively capture naturally occurring invariances.

To test whether the DEIs synthesized in our deep neural predictive model indeed elicit strong neuronal activities as predicted, we turned to *in vivo* verification. All MEIs and DEIs were first standardized for mean luminance and root mean square (RMS) contrast before presentation. We employed two approaches for DEI verification: (1) randomly selecting 10 DEIs from the set of 20 DEIs synthesized per neuron and presenting each stimulus 20 times, and (2) presenting all 20 DEIs once for each neuron. Additionally, each neuron’s MEI was presented with 20 repeats.

Our results validated that individual DEIs highly activated neurons *in vivo*, eliciting responses comparable to those evoked by MEIs (Fig. 2c–e, Suppl. Fig. S5a). Fig. 2c illustrates this for one example neuron, showing responses to 10 randomly selected DEIs alongside responses to the MEI and 100 random full-field natural images. For this neuron, only two out of ten DEIs elicited responses significantly lower than 85% of the MEI response (one after Benjamini-Hochberg (BH) correction for multiple comparisons). To determine whether our experimental procedure can reliably detect reductions in neuronal responses relative to the MEI, we analyzed the empirical probability of observing significantly lower activation levels given our sample size. For each predefined activation level below 85%, we randomly sampled two sets of 20 MEI trials with replacement for each neuron: one set scaled at 85% (reference) and the other at the selected activation level. We then applied a one-tailed Welch’s *t*-test to assess whether the latter exhibited significantly lower activation than the former. By repeating this process across 149 neurons, we systematically estimated the relationship between the observed fraction of significant tests and response reductions compared to MEI activation *in vivo* (Suppl. Fig. S5b). Empirically, across all neurons tested in closed-loop experiments from two mice, only 274 out of 1,490 DEIs (18.4%) elicited responses significantly lower than 85% of their corresponding MEI (4.7% after BH correction). Thus, we expect individual DEIs to evoke between 64 and 73% of the MEI response *in vivo* (95% CI, Suppl. Fig. S5b). If a substantial proportion of DEIs had activation levels below 64%, we would have observed a significantly greater fraction falling below the 85% threshold. Furthermore, finding natural images that elicit comparable activation is exceedingly rare. When searching 41 million natural image patches, only 0.17% produced responses exceeding 64% of the MEI response, highlighting the extreme sparsity of high-activating stimuli in natural vision. Thus, our results strongly support that DEIs are highly activating by design, and the absence of a larger fraction of weak DEIs is not due to insufficient sample size but rather reflects the inherent effectiveness of DEIs at driving neuronal responses.

We also evaluated whether a set of 20 DEIs, each presented once, elicited the same overall activation as a single randomly selected DEI presented 20 times. Our results demonstrated that this was the case, validating that DEI sets provide a reliable measure of neuronal activation. In subsequent experiments, we utilized DEI sets alongside control stimuli and systematic manipulations of DEIs to investigate their collective properties.

We first demonstrated that, similar to MEIs, DEIs were selective for the neurons they were designed to activate, consistently eliciting higher activity in their target neurons compared to non-target neurons (Fig. 2f, Suppl. Fig. S4). In addition, the digital twin accurately predicted the magnitude of neuronal responses to all synthesized MEIs and DEIs yielding median Pearson correlation coefficients of 0.74 and 0.75, respectively, between predicted and observed responses(Fig. 2g, h), further validating our approach. Importantly, DEIs strongly activated their target neurons *in vivo*, achieving 75 *±* 3% of their corresponding MEI activation (Fig. 2i), close to the model prediction of 85%; this effect was robust after controlling for eye movements (Suppl. Fig. S6).

One potential concern was that differences across DEIs might be indistinguishable to the animal, given the spatial acuity limits of the mouse visual system. To address this, we tested whether the mouse V1 population could detect differences across DEIs by presenting one randomly selected DEI pair for each neuron. We used a logistic regression classifier to decode the DEI identity from the *in vivo* V1 population responses. The analysis revealed a median classification accuracy of 80% across all neurons, substantially higher than the chance level of 50% (Fig. 2j). This result demonstrated that mouse V1, as a whole, can reliably discriminate between DEIs, indicating that the observed invariances in individual V1 neurons reflect relevant image transformations detectable at the V1 population level.

To evaluate whether DEIs represent specific image directions relative to the MEI, we conducted two control experiments comparing DEIs to perturbations of the MEI along random directions and along the natural image manifold. For the first control, we generated 20 synthetic images by perturbing the MEI in random directions while ensuring these images were closer to the MEI than the DEIs in pixel space (Fig. 3a, synthesized controls; Eq. 6). Unlike DEIs, these controls were not optimized for high neuronal activation. The second control involved searching millions of natural image patches to identify 20 images that were closer to the MEI than all DEIs, as measured by Euclidean distance in pixel space (Fig. 3a, natural image controls). During inception loop experiments, both synthetic and natural controls elicited lower responses in their target neurons compared to the corresponding DEIs (Fig. 3b, c). These results demonstrate that DEIs reflect invariances along specific directions away from the MEI in the image manifold, and that mere proximity to the MEI does not guarantee strong neuronal activation.

**Fig. 3.**
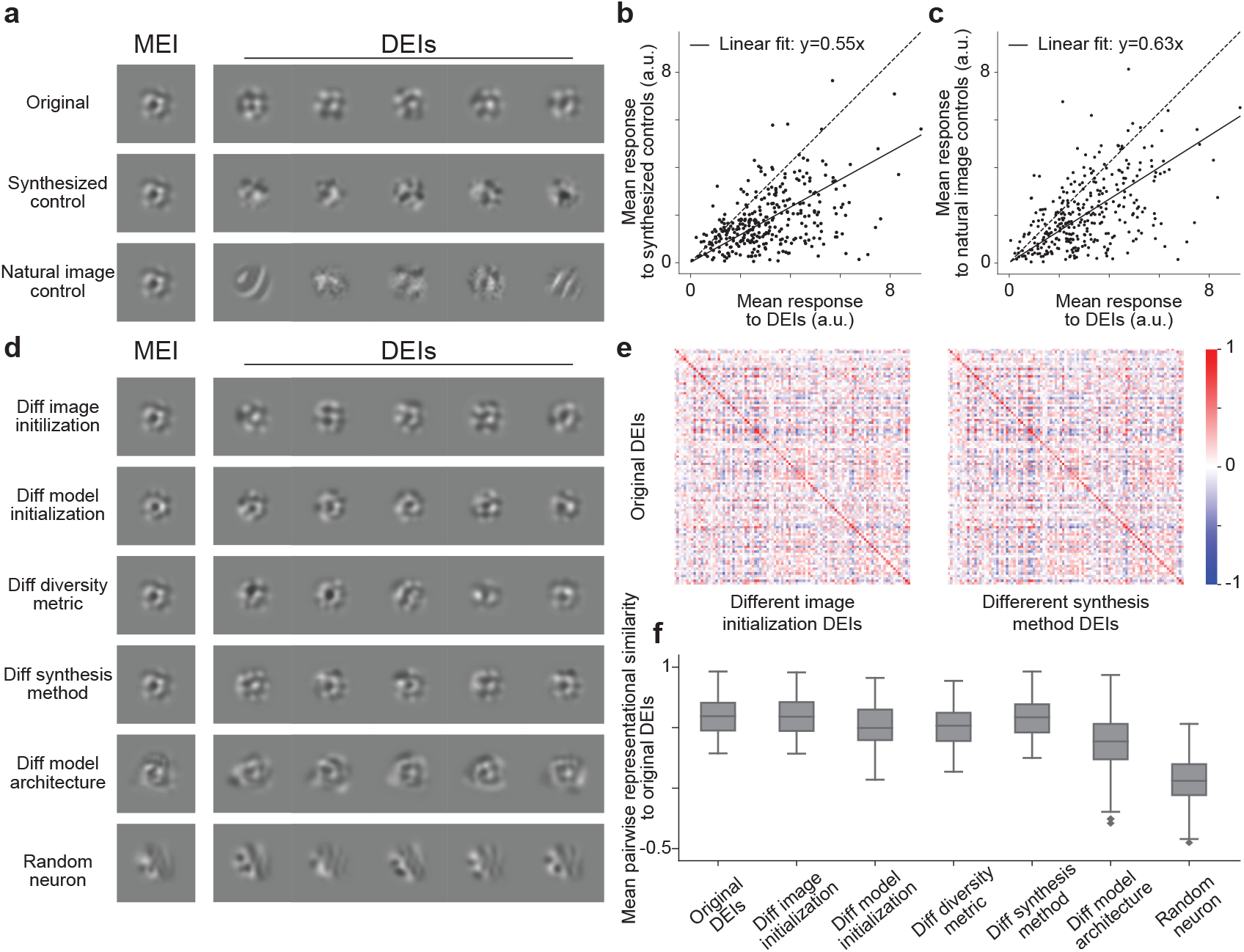
DEIs evoked stronger responses than controls and generalized across different synthesis conditions. **a**, MEI, DEIs (top), synthesized controls (middle), and natural controls (bottom) for one example neuron. Synthesized controls were generated by perturbing MEI in random directions, while natural controls were selected by searching through random natural patches. For each neuron, both controls were restricted to be closer to the MEI than all the DEIs as measured by Euclidean distance in pixel space. **b**, Synthesized controls failed to stimulate their target neurons *in vivo* compared to DEIs (55 *±* 2% of DEI activation, two-sided Wilcoxon signed-rank test, *W* = 3258, *P <* 10^−9^), with 36.8% neurons showing lower responses to synthesized controls compared to DEIs (20.8% after BH corrections; *P <* 0.05, two-sided Welch’s *t*-test with 30.4 average d.f.). **c**, Natural controls failed to stimulate their target neurons *in vivo* compared to DEIs (63*±* 3% of DEI activation, two-sided Wilcoxon signed-rank test, *W* = 6442, *P <* 10^−9^), with 31.5% neurons showing lower responses to synthesized controls compared to DEIs (16.0% after BH corrections; *P <* 0.05, two-sided Welch’s *t*-test with 31.1 average d.f.). **b, c** Response to each stimulus type was averaged over 20 different images with single repeat. Data were pooled from 318 neurons across 5 mice. **d**, MEI and DEIs for the same neuron in **a**, synthesized under various conditions: 1) different image initialization, 2) different model initialization, 3) different diversity metric, 4) different synthesis method (Baroni et al., 2023)), and 5) different model architecture (Willeke et al., 2022). **e**, DEIs synthesized under different conditions maintained high specificity to their target neuron. Confusion matrices showed *In silico* representational similarity between original DEIs and DEIs from different image initialization (left) or DEIs from a different synthesis method (Baroni et al., 2023) (right) (for other conditions, see Suppl. Fig. S8a). Each entry represents the mean pairwise cosine similarity between two sets of DEIs (see Methods for details). Representational similarity between original DEIs and DEIs synthesized from different conditions for the same neurons (diagonal) was larger than cross-neuron similarity (off-diagonal) (two-sided permutation test, *P <* 10^−4^ for all conditions after BH corrections). **f**, DEIs synthesized under different conditions closely resembled the original DEIs. The original DEIs were more similar to DEIs generated from various modifications in **d** than random neurons’ DEIs generated using the original method (two-sided Wilcoxon signed-rank test, *W* = 0, 0, 1, 0, 0, and 426, respectively; *P <* 10^−9^ for all conditions after BH correction). **e**,**f**, Data were pooled from 97 V1 neurons randomly sampled across 8 mice.

To verify the robustness of our DEI synthesis and to ensure that the observed differences were not artifacts of our methodology, we conducted control experiments varying key aspects of our approach (Fig. 3d). We demonstrated robustness of DEIs across various factors, including image initialization, model initialization, diversity evaluation metric, model architecture, and synthesis methodology (Fig. 3d). More specifically, critical controls included:

1. Altering image and model initialization;
2. Using cosine distance in V1 *in silico* population response to evaluate image diversity (“neuronal-space DEIs”);
3. Generating DEIs with an implicit neural representation model (INRM) (Baroni et al., 2023);
4. Using an alternative predictive model architecture (Willeke et al., 2022).

DEIs generated under these conditions consistently showed high specificity for target neurons (Fig. 3e, Suppl. Fig. S8a). To assess the representational similarity between DEI sets produced by different methods and models, we projected the images onto a latent space representing *in silico* population responses from an independent mouse (see Methods for details). DEIs generated using different methods for the same neurons exhibited higher similarity to the original DEIs than those from random neurons (Fig. 3f, Suppl. Fig. S8b). Additionally, DEIs from various methods displayed similar levels of diversity (Suppl. Fig. S8c). *In vivo* validation confirmed that “neuronal-space DEIs” activated target neurons comparably to pixel space DEIs (Suppl. Fig. S7). These results demonstrate that DEIs generalize across various modeling and synthesis conditions, suggesting the observed invariance is an intrinsic neuronal property rather than a methodological artifact.

### Bipartite parameterization of DEIs

Thus far, we have described the bipartite structure of V1 invariances in primarily qualitative terms. To further advance our understanding of this phenomenon, we now introduce concise, quantitative models that characterize these observed invariances with interpretable parameters. We first employed a texture synthesis model to characterize global shift invariance observed in neurons akin to classical complex cells. Neurons exhibiting this type of invariance maintain high activation when different crops of their preferred texture image are presented within the RFs. To model this, we extended the method from Cadena et al. (2018), synthesizing a full-field texture image for each neuron by maximizing the average *in silico* activation of randomly sampled crops (Fig. 4a, b, middle rows). We then sampled random crops from this optimized texture, which we termed “full-texture DEIs” (DEIs_full_). For many neurons, this global shift-invariant model proved in-adequate, and resulted in stimuli that visually deviated from the original non-parametric DEIs (Fig. 4b, c, middle vs top rows). This suggested a more nuanced form of invariance in V1 cells with heterogeneous RFs (Fig. 2a, b). To capture this complexity, we introduced the concept of “partial shift invariance”. This approach parameterized DEIs as the summation of two distinct, non-overlapping subfields within a neuron’s RF (Fig. 4a, b, bottom rows). The first subfield, directly cropped from the MEI, remained fixed across DEIs, and we denoted it as the “fixed subfield”. The second subfield, which we denoted as the “variable subfield”, exhibited shift invariance, maintaining high responses to different crops of a preferred texture image.

**Fig. 4.**
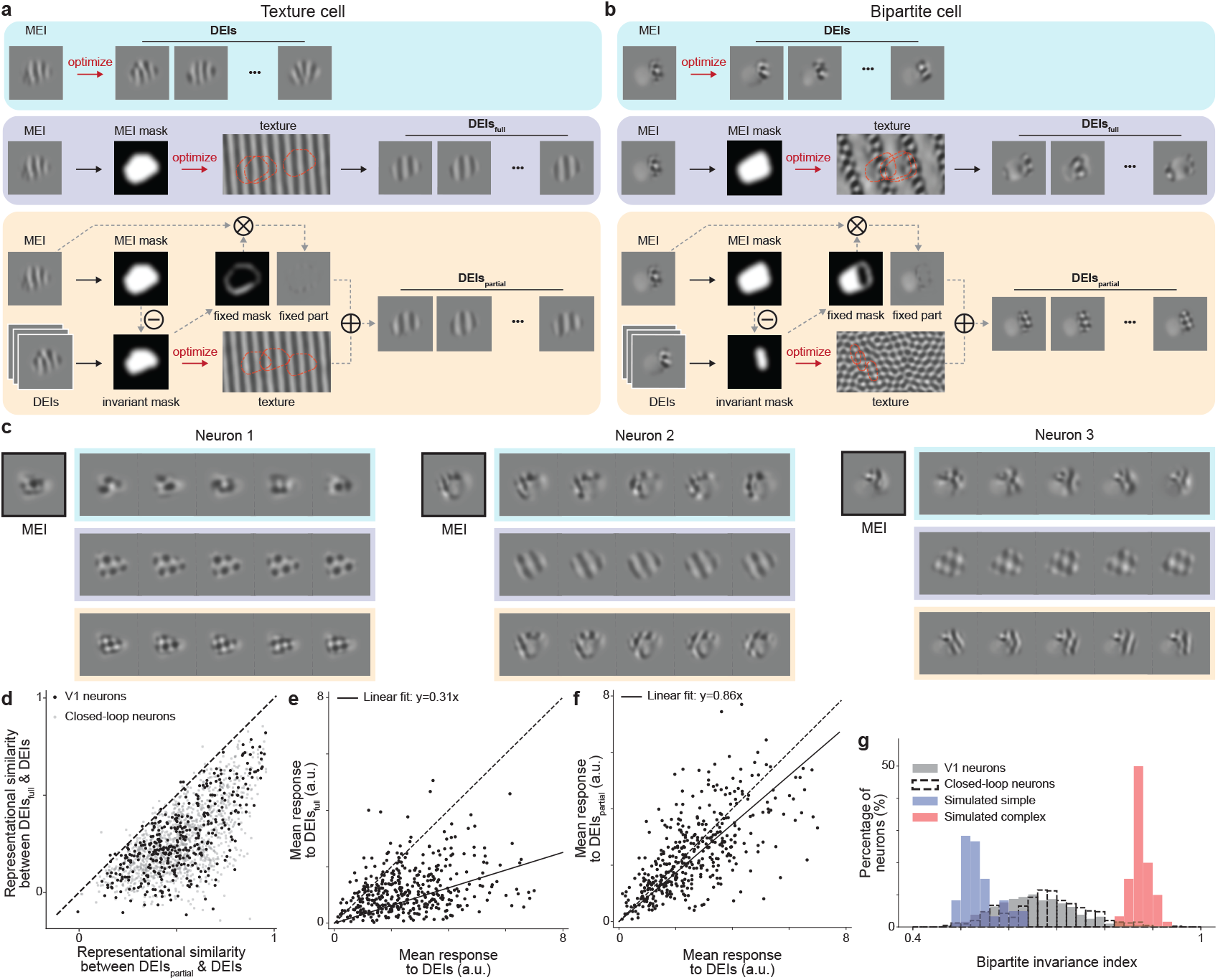
Bipartite parameterization reproduces the visual features and *in vivo* responses of non-parametric DEIs. **a, b**, Schematic of DEI synthesis using the non-parametric approach (DEIs, blue), full-texture parameterization (DEIs_full_, purple), and partial-texture parameterization (DEIs_partial_, orange) for an example V1 texture cell (left) and V1 bipartite cell (right). DEIs_full_ were synthesized by optimizing an underlying texture canvas, from which random crops masked by the MEI mask maximally activated the target neuron. In contrast, DEIs_partial_ comprised two distinct, non-overlapping subfields: a fixed subfield directly masked from the MEI, and a shift-invariant subfield preferring random crops from a texture image synthesized similarly to DEIs_full_, but using only part of the MEI mask for texture optimization. **c**, MEI, DEIs, DEIs_full_, and DEIs_partial_ for three example neurons, with each DEI type indicated by the corresponding color from **a. d**, DEIs_partial_ were more similar to their corresponding non-parametric DEIs than DEIs_full_ for both random V1 neurons and closed-loop neurons (two-sided Wilcoxon signed-rank test, *W* = 2783, *P <* 10^−9^, and *W* = 65, *P <* 10^−9^, respectively). **e**, DEIs_full_ failed to stimulate their target neurons *in vivo* compared to non-parametric DEIs (31 *±* 2 of DEI activation, two-sided Wilcoxon signed-rank test, *W* = 4389, *P <* 10^−9^) with 43.4% of all neurons showing different responses to DEIs_full_ than DEIs (29.4% after BH corrections) (*P <* 0.05, two-sided Welch’s *t*-test with 29.4 average d.f.). **f**, DEIs_partial_ activated their target neurons *in vivo* similarly to non-parametric DEIs (86 *±* 4 of DEI activation, two-sided Wilcoxon signed-rank test, *W* = 32429, *P* = 7.0 ×10^−4^) with only 8.5% of all neurons showing different responses (0.0% after BH corrections) (*P <* 0.05, two-sided Welch’s *t*-test with 33.5 average d.f.). **e, f**, *In vivo* responses to DEIs, DEIs_full_, and DEIs_partial_ were averaged across 20 different images with single repeat. **g**, Bipartite invariance indices of V1 neurons were larger than those of simulated simple cells (60 cells, blue) and lower than those of simulated complex cells (60 cells, red) (*P <* 10^−9^ for both, two-sided Welch’s *t*-test with 95.5 and 213.8 d.f., respectively; see *Bipartite invariance index* in Methods for details). Data were pooled from 6 mice, displaying a total of 1200 neurons for random V1 neurons; closed-loop neurons comprised 401 neurons pooled from 8 mice.

To identify these subfields, we employed the following procedure:

1. The variable subfield was defined by selecting the region with the highest pixel-wise variance across non-parametric DEIs, progressively expanding it by adjusting the threshold of pixel-wise variance.
2. The fixed subfield was defined as the complement of the variable subfield.
3. For each candidate variable subfield, we optimized a full-field texture image and generated texture-based DEIs by combining random crops from this texture with the corresponding fixed subfield from the MEI.

To quantify each neuron’s partial shift invariance, we examined the relationship between the *in silico* response and variable subfield size (Suppl. Fig. S9a). We summarized this with a “bipartite invariance index”, calculated as the Area Under the Curve (AUC) of a quadratic spline fit to the activation-subfield size trade-off curve (Suppl. Fig. S9b, c). This index revealed differences between neuron types: simulated simple cells exhibited the lowest values (median=0.53), simulated complex cells had the highest values (median=0.87), while V1 neurons fell in between (median=0.65) (Fig. 4g). The bipartition index showed high consistency across model ensembles (Pearson *r* = 0.66, Suppl. Fig. S9d). For each neuron, we selected the variable subfield by maximizing the harmonic mean of the texture-based DEIs’ *in silico* response and their diversity. We termed these optimized stimuli “partial-texture DEIs” (DEIs_partial_). Remarkably, DEIs_partial_ visually resembled the non-parametric DEIs more closely than DEIs_full_, as quantified by representational similarity (Fig. 4d). During closed-loop experiments, DEIs_partial_ activated neurons at a level comparable to non-parametric DEIs (86% of the DEI response), while DEIs_full_ elicited much weaker *in vivo* responses (31% of the DEI response) (Fig. 4e, f). Notably, we still observed strong *in vivo* responses to both DEIs_partial_ and non-parametric DEIs even after excluding neurons whose DEIs_partial_ were dominated by fixed subfields resembling the MEI (Suppl. Fig. S10).

We next tested the necessity and specificity of the two subfields in DEIs_partial_ by isolating or swapping the content within each subfield (Suppl. Fig. S11a). We found that both subfields were necessary for high activation—masking out the fixed or variable subfield content from the MEI reduced *in vivo* responses in target neurons to 74% and 33%, respectively (Suppl. Fig. S11b, c). Similarly, the contents within both subfields were highly specific. Replacing the fixed subfield content with random natural image patches, or swapping the optimized texture image for the variable subfield with textures from other neurons in DEIs_partial_ decreased activity to 55% and 74%, respectively (Suppl. Fig. S11d, e). While our closed-loop validation primarily focused on neurons exhibiting high levels of invariance, we also randomly selected neurons from all reliable and well-predicted V1 neurons (corresponding to 79.0% *±* 0.5% of all neurons imaged per scan) for closed-loop verification. This confirmed that our findings generalized to the broader population (Suppl. Fig. S12).

We also conducted a series of additional control experiments to validate our findings and to rule out alternative explanations for the observed bipartite RF structure:

1. Imaging contamination: To mitigate concerns about potential contamination in the calcium imaging data, we replicated our findings using Neuropixels (Jun et al., 2017) recordings from mouse V1 (Suppl. Fig. S13, S14). The electrophysiology results revealed similar distributions of diversity and bipartite invariance indices between spike-sorting-identified “single units” and two-photon imaging units while “multi-unit activity” (MUA) exhibited higher values for these indices (Suppl. Fig. S14c, d; see Methods for details). Importantly, across single units, we found no correlation between either metric and inter-spike-interval (ISI) violations (Hill et al., 2011), a measurement of unit contamination (Suppl. Fig. S14g, h). These findings indicate that the observed bipartite structure reflects genuine properties of V1 L2/3 neurons rather than an artifact of single-unit contamination.
2. Shift invariance within fixed subfield and model complexity: To address the possibility that the fixed subfield also exhibits shift invariance and to ensure that the observed bipartite structure is not merely a consequence of increased model complexity, we developed a “two-variable-subfield” model. This model optimized two distinct textures independently for both the variable and the fixed subfields (Suppl. Fig. S15a bottom, b, c). While this approach produced slightly more diverse images by design, these images were less similar to the original DEIs and less effective at activating neurons *in silico* (eliciting 69% of the DEI_partial_ activation, Suppl. Fig. S15d–f). These findings underscore the functional distinction between the two subfields and demonstrate that the efficacy of the bipartite parameterization is not simply due to increased model complexity.
3. Necessity of spatial division: To determine whether the spatial division of the receptive field is essential for the observed bipartite invariance, we developed a “no-spatial-division” parameterization. In this model, DEIs were parameterized as the summation of two fully superimposed subfields, unlike the non-overlapping subfields in the bipartite parameterization (Suppl. Fig. S16a–c). This alternative model performed worse in capturing both *in silico* activation and diversity of DEIs (88% of the DEI_partial_ activation and 75% of the DEI_partial_ diversity, Suppl. Fig. S16d-h), confirming the importance of spatial division in our model.
4. Center-surround interaction: To determine whether the observed bipartite structure might merely be a manifestation of center-surround organization arising from extra-classical surround modulation, we performed additional experiments where we presented sparse noise stimuli and natural images in the same scans. We found that the fixed and variable subfields did not maintain a consistent spatial relationship with the classical “minimum response field” (MRF) (Suppl. Fig. S17c, d). A companion paper also found that the extra-classical excitatory contextual modulation fields in mice are much larger than the size of the MEI (approximately 20 degrees larger; see Fig. 2e in Fu et al. (2024)). These findings collectively indicate that the bipartite structure represents a novel organization beyond the classical center-surround framework.
5. Eye movements: To address the potential influence of trial-to-trial eye movements on the observed bipartite RFs, we trained separate models on subsets of trials using inclusion criteria based on different thresholds of eye movement size. We found that the resulting DEIs were highly similar across all models (Suppl. Fig. S18a, b), and the bipartite invariance index showed no significant variation among these models (Suppl. Fig. S18c). These results showed that the observed bipartite RF structure is an intrinsic property of V1 neuronal responses, rather than an artifact of eye movements.

Collectively, these findings demonstrate that V1 neuron receptive fields are best characterized by a bipartite structure, featuring one subfield that prefers a fixed spatial pattern and another that optimally responds to random crops of an underlying texture image.

### Bipartite structure aligns with natural object boundaries defined by spatial frequency differences

Previous studies have demonstrated that Most Exciting Inputs (MEIs) capture complex spatial features prevalent in natural scenes (Walker et al., 2019). Building on this insight and the bipartite RF structure we identified, we hypothesized that these neurons may play a role in visual segmentation by detecting object boundaries through variations of texture (Zhan and Baker Jr, 2006). Given our partial-texture model’s partitioning of the RF into two often distinct subfields, we further postulate that V1 neurons might preferentially respond to object boundaries defined by texture discontinuities.

To test this hypothesis, we utilized a natural image dataset with manual segmentation labels, Caltech-UCSD Birds-200-2011 (CUB) (Wah et al., 2011). The CUB dataset is a comprehensive collection of 11,788 images spanning 200 bird species, each annotated with pixel-resolution segmentation masks for object and background. We screened over a million crops from the CUB dataset *in silico*, matching mean and RMS contrast to the MEI and DEIs, to identify highly activating crops for each V1 neuron (Fig. 5a). Across the population, highly activating crops were more likely to contain object boundaries than random crops (Suppl. Fig. S19b). To further quantify the alignment between the bipartite RF structure and the object boundaries in highly activating CUB crops, we computed a matching score between the segmentation label and the “bipartite mask” defined from DEIs_partial_ (Fig. 5a, see Methods for details). Highly activating image crops exhibited better alignment between bipartite subfield divisions and object boundaries compared to random crops, indicating a preferential response to object-background divisions (Suppl. Fig. S19a, Fig. 5b).

**Fig. 5.**
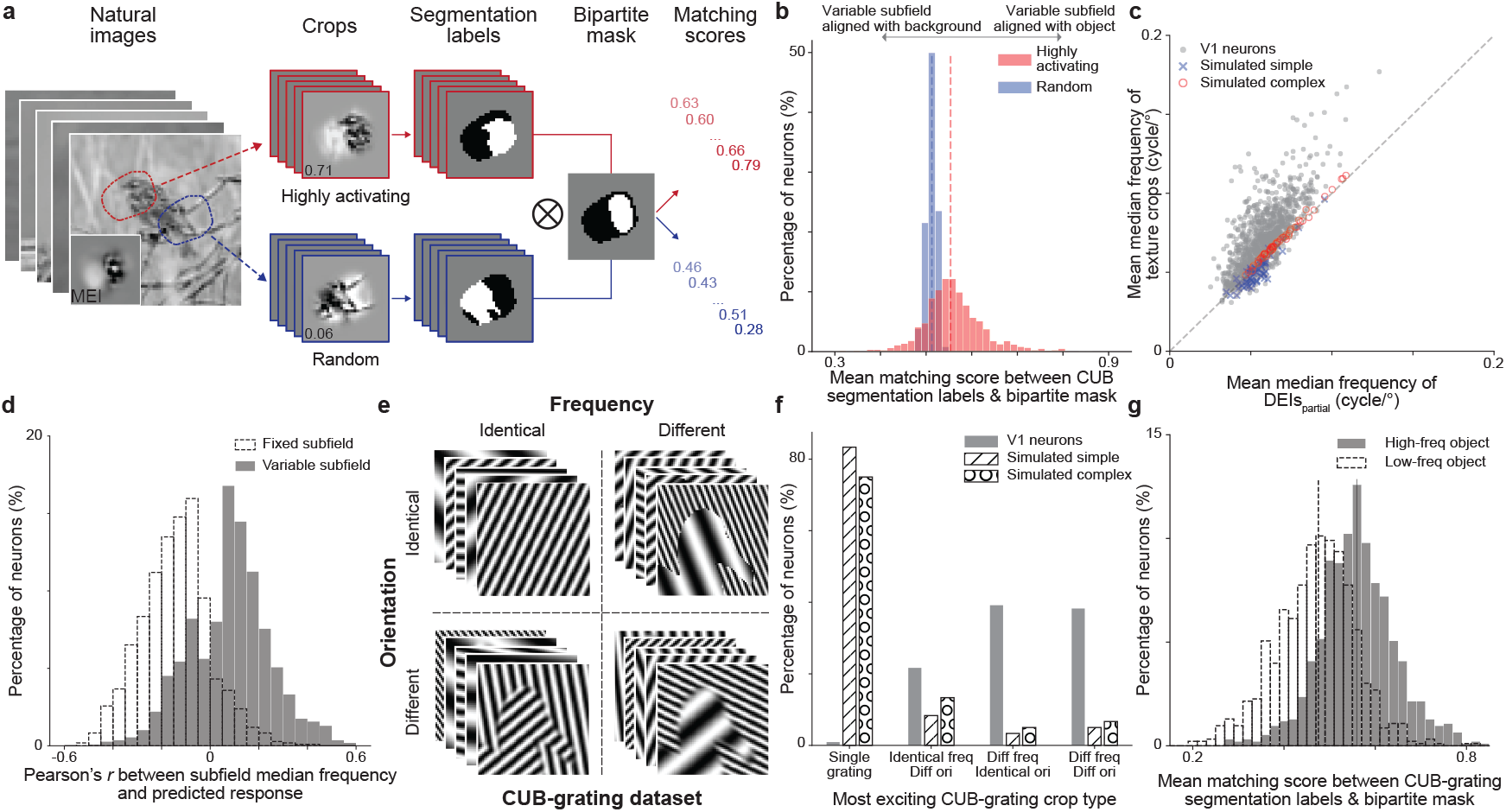
Bipartite structure aligns with natural object boundaries formed by spatial frequency differences. **a**, We screened over 1 million crops from the Caltech-UCSD Birds-200-2011 (CUB) dataset using our predictive model to find 100 most highly activating (red) and 100 random (blue) crops for each neuron, along with their corresponding *in silico* responses and manual segmentation labels. We computed a matching score for each crop based on its segmentation label (white denotes object, black denotes background) and the neuron’s “bipartite mask” defined by its DEIs_partial_ (white denotes variable subfield, black denotes fixed subfield). A score of 1 indicated perfect matching between the variable subfield and object (white to white) within the RF while a score of 0 indicated the opposite. **b**, Highly activating natural crops with object boundaries yielded higher matching scores than random natural crops with object boundaries (two-sided Wilcoxon signed-rank test, *W* = 82849, *P <* 10^−9^), with 51.4% of all neurons showing greater matching scores for highly activating crops than random natural crops (46.4% after BH correction) while only 4.9% showing lower matching scores to highly activating crops (4.1% after BH correction) (*P <* 0.05, two-sided Welch’s *t*-test, with 76.2 average d.f. respectively). One neuron (0.08%) was excluded from this analysis as it strictly preferred crops without object boundaries. **c**, Most V1 neurons preferred higher spatial frequency content in the variable subfield compared to the fixed subfield. The mean median frequency of texture crops was higher than that of DEIs_partial_ (two-sided Wilcoxon signed-rank test, *W* = 34563, *P* = 3.0 × 10^−6^) with 76.5% of all neurons preferring content with a higher median spatial frequency in the variable subfield than in the fixed subfield (75.5% after BH correction) while only 5.3% of neurons preferring the opposite (5.0% after BH correction) (*P <* 0.05, two-sided Welch’s *t*-test with 33.3 average d.f. respectively). Texture crops were synthesized by replacing content in DEIs_partial_ fixed subfield with random crops from the texture optimized for the variable subfield. Median frequencies were averaged across 20 different texture crops and DEIs_partial_, respectively. In contrast, simulated simple cells (blue cross) showed no preference for median frequency between the two subfields (two-sided Wilcoxon signed-rank test, *W* = 284, *P* = 3.0 × 10^−6^). Simulated complex cells (red circle) preferred higher median frequency in the variable subfield (two-sided Wilcoxon signed-rank test, *W* = 22, *P <* 10^−9^), albeit with marginal effect. **d**, V1 neuronal responses correlate with spatial frequency within the variable and fixed subfield. We applied the fixed subfield mask to the CUB natural image dataset to extract 10,000 crops and computed their median frequency. We then combined these crops with the original variable subfield from the MEI and passed the resulting images through the predictive model to obtain the predicted responses. For the majority of neurons (79.08%), we observed a negative correlation between the fixed subfield’s median frequency and the predicted response (median =−0.14, one-tailed one sample *t*-test against mean of 0, *t* = −33.38, *p <* 10^−9^, *d*.*f*. = 1089). In contrast, for most neurons (64.75%), we observed a positive correlation between the variable subfield’s median frequency and the predicted response (median = 0.09, one-tailed one sample *t*-test against mean of 0, *t* = 16.23, *p<* 10^−9^, *d*.*f*. = 1083). All p-values were corrected for multiple comparison using BH procedure. Four neurons were excluded in the fixed subfield analysis due to excessively small fixed subfield size. **e**, Parametric “CUB-grating” dataset with the CUB segmentation labels. The object and background content were replaced with grating patterns of four different types: 1) homogeneous pattern without using segmentation labels (“single grating”); 2) same spatial frequency but different orientations; 3) same orientation but different spatial frequencies; and 4) different frequencies and orientations (see Methods for more details). **f**, Using CUB-grating, we identified the most activating crop for each neuron (similar as in **a**). Simulated simple and complex cells predominantly preferred single grating images (83.3% and 75%, respectively). In contrast, V1 neurons exhibited a different pattern of preference (one-way chi-squared test, *χ*^2^ = 8510, *P <* 10^−9^, and *χ*^2^ = 5538, *P <* 10^−9^ for comparison against simulated simple and complex cells, respectively). While most simulated simple (83.3%) and complex (75%) cells preferred single grating images, V1 neurons almost exclusively preferred images with object boundaries (99.1%). V1 neurons showed preferences for boundaries defined by differences in spatial frequency alone (39.2%), orientation alone (21.6%), or a combination of both (38.3%). The marginal difference in preference was greater for spatial frequency than for orientation (*p<* 0.05, two-sided marginal difference bootstrapping). **g**, We divided CUB-grating images with different object-background spatial frequencies into two datasets: 1) “high-frequency object” and 2) “low-frequency object”. Following the procedure in **a**, we identified 100 most activating crops for each dataset and calculated the mean matching score for neurons preferring images containing different frequencies (from **f**). The matching scores for “high-frequency object” dataset were higher than those for “low-frequency object” (two-sided Wilcoxon signed-rank test, *W* = 340648, *P <* 10^−9^) with 66.4% of all neurons showed greater matching scores for “high-frequency object” crops (same after BH correction), while 23.4% showed smaller matching scores (23.3% after correction) (*P <* 0.05, two-sided Welch’s *t*-test, 170.2 average d.f.). **b, c, f, g**, These results generalized across different inclusion criteria used to identify patches containing object boundaries (Suppl. Fig. S20). **a–g**, Data were pooled from 6 mice, including 1200 randomly selected neurons. Simulated simple and complex cells included 60 neurons each.

Next, we investigated which low-level visual statistics contribute to these alignment results. Analysis of DEIs_partial_ revealed that most V1 neurons (76.5%) preferred spatial pattern with higher median frequency in the variable subfield compared to the fixed subfield (Fig. 5c; see Methods for details). Notably, this preference was absent in simulated simple and complex cells subjected to the same optimization procedure (Fig. 5c). To test whether this bias extends to natural images, we analyzed *in silico* responses to natural image patches with varying frequency biases. We found that for most neurons, natural patches with higher frequency content in the variable subfield elicited stronger activation (64.8%), whereas the fixed subfield exhibited the opposite trend, with lower frequency content inducing stronger activation (79.1%) (Fig. 5d). These findings led us to hypothesize that V1 neurons are particularly sensitive to object boundaries defined by differences in spatial frequency.

To explicitly test this hypothesis, we created a modified CUB dataset (“CUB-grating”) where natural image content was replaced with grating stimuli of varying spatial frequencies and orientations, while preserving naturalistic boundaries, and presented crops from this dataset *in silico* (Fig. 5e). Our analysis revealed striking differences between simulated cells and V1 neurons. While most simulated simple (83.3%) and complex (75%) cells preferred single grating images, V1 neurons almost exclusively preferred images with object boundaries (99.1%) (Fig. 5f). Specifically, V1 neurons showed preferences for boundaries defined by differences in spatial frequency alone (39.2%), orientation alone (21.6%), or a combination of both (38.3%). Notably, the difference in preference was greater for frequency than for orientation (Fig. 5f). Similar to highly activating natural crops (Fig. 5c), segmentation labels in highly activating CUB-grating images also aligned with the bipartite mask (Fig. 5g). However, in highly activating CUB-grating patches, the variable and fixed portions of the bipartite mask corresponded to high and low spatial frequency regions, respectively, rather than the object or background areas. Our analysis revealed that mouse V1 neurons preferentially responded to object boundaries defined by frequency discontinuities, with the variable subfield favoring higher spatial frequency than the fixed subfield.

### The MICrONS dataset reveals synaptic connectivity reflecting a functional invariance hierarchy in V1 Layer 2/3

To gain insights into the synaptic-level cortical architecture and its relationship to neuronal response invariances, we sought to connect functional properties of neurons with their inter-neuronal connectivity patterns. This investigation was made possible by recent advances in large-scale functional connectomics, which allow for simultaneous measurement of neuronal activity and synaptic connectivity in the same tissue. We utilized the MICrONS functional connectomics dataset, which includes responses from over 75,000 excitatory neurons to natural movies and reconstructed synaptic-level connectivity derived from electron microscopy data at nanometer resolution (MICrONS Consortium et al., 2021). To quantify functional invariances, we employed a dynamic digital twin model of the MICrONS mouse that uses the foundation core model from Wang et al. (2023) (see Methods for details), which accurately predicted responses to various stimulus domains including natural movies, static images, and artificial parametric stimuli.

We first validated our pipeline’s accuracy in computing DEIs from the dynamic digital twins by conducting closed-loop experiments on three new mice (Fig. 6a). These dynamic digital twins were trained following the same procedure as the MICrONS mouse digital twin. For each mouse, we recorded neuronal responses to both our static natural image sets and the same dynamic movie stimuli (including natural and parametric movie clips) used in the MICrONS dataset. We then trained two deep CNN models for each mouse using the same architecture as in our main experiments:

**Fig. 6.**
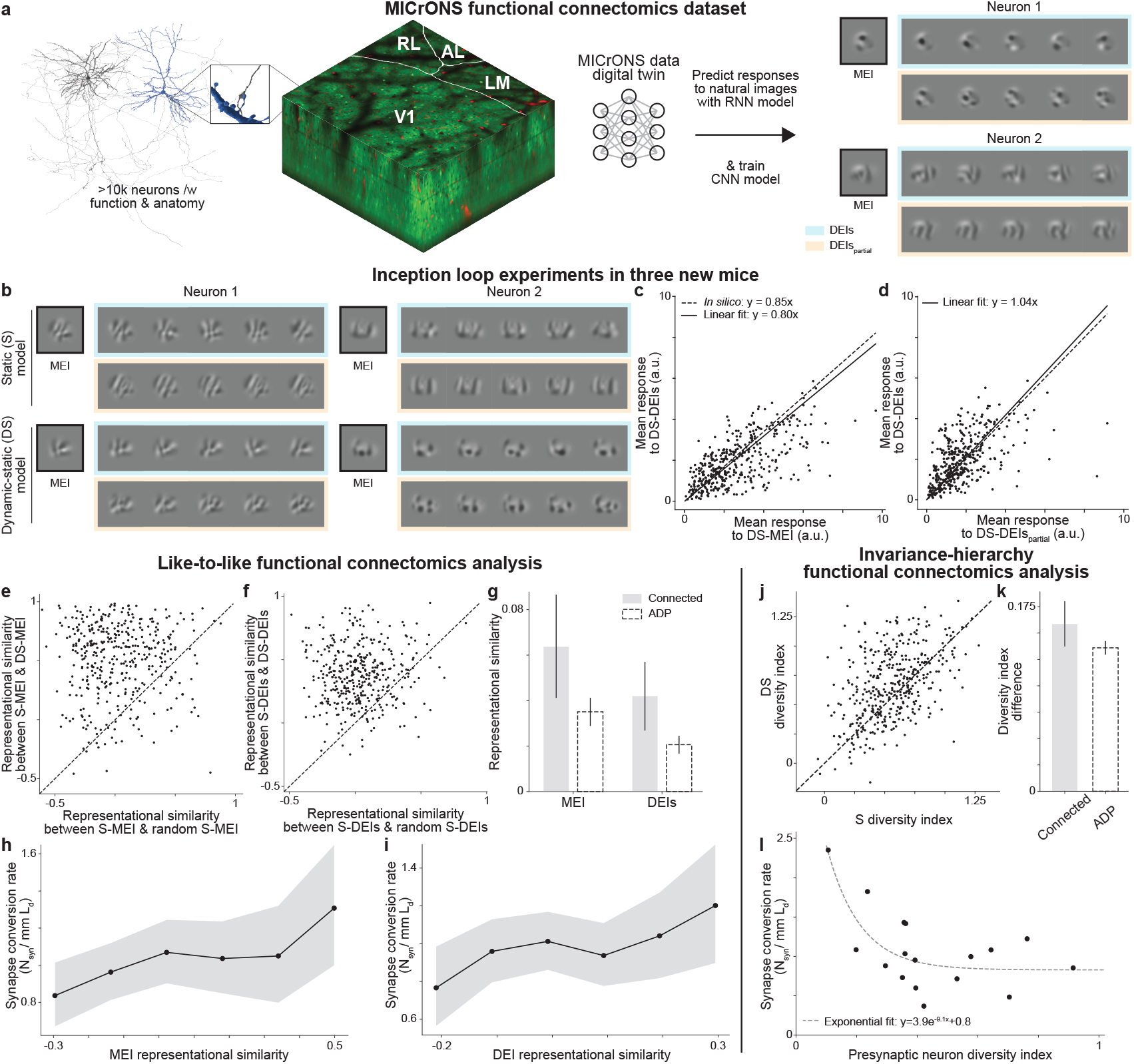
MICrONS functional connectomics analysis. **a**, Schematic of the MICrONS functional connectomics dataset (MICrONS Consortium et al., 2021), comprising responses of >75k neurons to dynamic stimuli and their reconstructed sub-cellular connectivity from electron microscopy data. We employed the MICrONS “digital twin” (Wang et al., 2023), trained on dynamic stimuli (denoted as “dynamic” model) to predict responses to natural images used in our experiments. A new model was trained on these *in silico* predictions (“dynamic-static” or “DS” model) and used to synthesize MEIs, DEIs, and DEIs_partial_. **b–d**, *In vivo* verification of stimuli synthesized from the DS model. **b**, MEIs and DEIs optimized using our standard model (“static” or “S”) and the DS model for two example neurons. **c**, DS-DEIs stimulated neurons *in vivo* at 80 *±* 3% of DS-MEI activation, close to the *in silico* prediction of 85% (two-sided Wilcoxon signed-rank test, *W* = 31534, *P* = 2.8 ×10^−4^), with only 10.3% of all neurons showing different responses between DEIs and 85% of MEI (0.25% after BH correction) (*P <* 0.05, two-sided Welch’s *t*-test with 32.0 average d.f.). **d**, DS-DEIs_partial_ activated target neurons similarly to DS-DEIs (two-sided Wilcoxon signed-rank test, *W* = 29878, *P* = 1.4× 10^−5^) with only 9.5% of all neurons showing different responses (0.0% after BH correction) (*P <* 0.05, two-sided Welch’s *t*-test with 32.0 average d.f.). **c, d**, DS-MEI responses were averaged across 20 repeats of the same image while DS-DEIs and DS-DEIs_partial_ responses were averaged across 20 different images with single repeat. **e–i**, Like-to-like functional connectomics analysis in V1 Layer 2/3 using the MICrONS dataset. **e**, DS-MEIs were more similar to S-MEIs of the same neuron than S-MEIs of other random neurons (two-sided Wilcoxon signed-rank test, *W* = 4537, *P <* 10^−9^). **f**, Similarly, DS-DEIs were more similar to S-DEIs of the same neuron than S-DEIs of other random neurons (two-sided Wilcoxon signed-rank test, *W* = 3969, *P <* 10^−9^). **g** MEIs and DEIs of connected pairs (0.06 *±* 0.02 and 0.04 *±* 0.02) are more similar than those of the Axonal-Dendritic Proximity (ADP) control pairs (Ding et al., 2023) (0.03 *±* 0.01 and 0.021 *±* 0.004)(*P <* 10^−4^, two-sided bootstrapped mean difference after BH correction). **h, i**, Synapse conversion rate increased linearly with the MEI (**h**) and DEI (**i**) representational similarity for neuron pairs (*P* = 0.014 and 0.0034, respectively, two-sided *t*-test for linear coefficient against 0 using Poisson generalized linear mixed model with random intercepts). Neuron pairs were binned by their MEI and DEI similarity, respectively. **j–l**, Invariance-hierarchy analysis in V1 Layer 2/3 using the MICrONS dataset. **j**, Diversity indices from the DS model highly correlated with those from the S model (Pearson *r* = 0.46, *P <* 10^−9^, two-sided *t*-test). **k**, Connected pairs showed larger diversity index increase than Axonal-Dendritic Proximity (ADP) controls (0.16 *±* 0.02 and 0.14 *±* 0.01, respectively; *P <* 10^−4^, two-sided bootstrapped mean difference against 0 after BH correction). Diversity index difference is defined as the change in diversity index between the postsynaptic (or the ADP) and the presynaptic neuron for a connected pair and a ADP control pair, respectively. **l**, Presynaptic neurons with lower diversity indices showed higher synapse conversion rate (Spearman’s rank correlation coefficient *ρ* = 0.52, *P* = 0.02, two-sided *t*-test). This relationship was well-modeled by an exponential decay (*R*^2^ = 0.56). **c–j**, Data for *in vivo* verification of the dynamic-static model were pooled over 399 neurons from 3 mice. **g–l**, Data for MICrONS functional connectomics analysis were pooled over 19 presynaptic neurons forming 706 connected pairs and 18,162 ADP controls. Error bars and shaded areas represented 95% confidence intervals from 10,000 bootstraps.

1. A “static” (*S*) model trained directly on *in vivo* responses to static natural images, replicating our standard approach.
2. A “dynamic-static” (*DS*) model trained on *in silico* responses to static natural images generated by its dynamic digital twin counterpart.

Both *S* and *DS* models accurately predicted responses to a held-out test set of static natural images (Suppl. Fig. S21a). We found that MEIs, DEIs, and DEIs_partial_ synthesized from the *DS* model were visually similar to those from the *S* model (Fig. 6b), with high representational similarity (Fig. 6e, f, Suppl. Fig. S21e). The diversity indices and bipartite invariance indices calculated from the *DS* model also closely matched those of the *S* model, yielding Pearson correlations of 0.46 and 0.66, respectively (Fig. 6j, Suppl. Fig. S21f). Moreover, when presented back to the animals during inception loop experiments, the MEIs, DEIs, and DEIs_partial_ synthesized from the *DS* model elicited high activities in their target neurons (Suppl. Fig. S22, S23, S24). DEIs achieved 80 *±* 3% of their corresponding MEI activation *in vivo* (Fig. 6c), closely matching the model’s prediction of 85%. DEIs_partial_ evoked equally strong responses as DEIs (Fig. 6d). This agreement between *in silico* predictions and *in vivo* responses demonstrated the validity of using dynamic digital twins to study neuronal invariances. It confirms that our analysis pipeline, when applied to the MICrONS dataset, can reliably capture and replicate the functional properties of real neurons, including their response invariances.

In our analysis of the MICrONS dataset, we focused on V1 Layer 2/3 excitatory neurons that had anatomically matched counterparts in the EM volume. We then restricted our analysis to neurons with high response reliability (*CC*_*max*_ *>* 0.4), accurate digital twin predictions (*CC*_*abs*_ *>* 0.2), comprising 77% of all functional recorded neurons. When combining with the manually proofread connectivity graph of the MICrONS dataset (MICrONS Consortium et al., 2021), these resulted in 19 presynaptic neurons and 570 postsynaptic partners, forming 706 connected pairs in V1 Layer 2/3.

A well-established principle in the functional connectomics domain is the like-to-like connectivity rule—excitatory neurons with similar response properties are more likely to form connections (Ko et al., 2011; Wertz et al., 2015; Lee et al., 2016; Rossi et al., 2020; Ding et al., 2023). We reexamined the like-to-like connectivity rule through the lens of MEI and DEI similarities, leveraging the synaptic-level resolution of the MICrONS dataset. We employed the Axonal-Dendritic Proximity (ADP) control—introduced in the companion paper by Ding et al. (2023) and computed via the NEURD connectomics analysis package (Celii et al., 2023)—to identify neuron pairs that have the physical opportunity to connect but do not, accounting for the inhomogeneous distribution of axons and dendrites in the cortex. Our analysis confirmed that synaptically connected neuron pairs have more similar MEIs and DEIs than ADP controls in V1 Layer 2/3 (Fig. 6g). These results provide strong evidence that the like-to-like rule (Ko et al., 2011; Wertz et al., 2015; Lee et al., 2016; Rossi et al., 2020; Ding et al., 2023) operates with synaptic-level precision rather than being simply a byproduct of broader spatial patterns of neuronal organization. We further analyzed the relationship between functional similarity and the synapse conversion rate (Lee et al., 2016; Ding et al., 2023)—defined as the number of synapses formed per unit length of axon-dendrite overlap between neuron pairs. We found that the synapse conversion rate increased linearly with the representational similarity of MEIs and DEIs for neuron pairs (Fig. 6h, i). These findings further corroborate the results reported by Ding et al. (2023), who demonstrated that the like-to-like connectivity rule in the feature domain operates at the synaptic level across different types of connections, both within and across cortical layers and areas.

We next investigated the relationship between neuronal invariance and circuit structure. Hierarchical models of the cortex have long speculated that complex functional invariance could arise from the convergence of excitatory pre-synaptic inputs with simpler invariances (Hubel and Wiesel, 1962; Riesenhuber and Poggio, 1999; Serre and Riesenhuber, 2004). The seminal example is Hubel & Wiesel’s hypothesis that complex cells achieve phase invariance by combining inputs from spatially aligned simple cells with similar orientation but different phase preferences (Hubel and Wiesel, 1962). However, evidence for this decades-old model has primarily relied on correlational analyses (Alonso and Martinez, 1998), with direct evidence remaining elusive due to the challenge of simultaneously measuring both physiology and wiring of the same neurons. We hypothesized that synaptically connected pairs of neurons would exhibit a greater increase in functional invariance compared to ADP controls. We first validated that the diversity indices calculated from our *DS* model closely matched those from the *S* model (Fig. 6j). Our analysis revealed that synaptically connected neuron pairs indeed showed greater increases in diversity index than ADP controls (Fig. 6k), suggesting that the increase in functional invariance occurs at the synaptic level. Intriguingly, we found no difference between the mean diversity indices of postsynaptic partners and ADP controls (Suppl. Fig. S25). Furthermore, we found that the synapse conversion rate decreased exponentially as the presynaptic neuron’s diversity index increases (Fig. 6l). This relationship implies that excitatory neurons with lower functional invariance are more likely to form intralaminar connections in V1 Layer 2/3. Collectively, these findings provide evidence for a hierarchical organization among excitatory neurons in mouse V1 Layer 2/3 that enhances single-neuron functional invariance.

Our study demonstrates the utility of employing the dynamic digital twin of the MICrONS mouse for novel downstream analyses and experiments beyond the scope of the original experimental design. This approach is particularly valuable given the scarcity and complexity of obtaining large-scale data that integrate both dense synaptic-level structure and function within the same brain tissue.

## Discussion

Invariant object recognition is a challenge central to visual perception. One perspective on how this is achieved by the visual system is the *object manifold disentanglement hypothesis* (DiCarlo and Cox, 2007). Within this framework, an object manifold can be thought of as the continuous set formed by applying all natural transformations to a given object—e.g. rotations, scalings, translations, lighting changes, etc. In the pixel space, the manifolds corresponding to different objects are entangled and difficult to distinguish. The idea core to this hypothesis is that the visual system gradually disentangles these manifolds through stages of hierarchical processing along the ventral visual pathway, ultimately enabling the linear decoding of object identity from the population representation at the top of the hierarchy. Experimental evidence suggests that single neurons in higher visual areas extract and integrate information from simpler feature detectors in lower areas to represent more complex features and construct invariances to feature transformations such as translation, rotation or change of texture (Hubel and Wiesel, 1962; Tanaka, 1996; Poggio and Bizzi, 2004; Yamins et al., 2014). This idea inspired the original development of the convolutional neural network architecture (Fukushima, 1980), and is reflected in the single neuron response properties that emerge in networks trained on image recognition tasks (Olah et al., 2017, 2020).

Characterizing the structure of neuronal invariances is crucial for understanding how the brain accomplishes the challenging task of object recognition. However, thus far, systematic characterization of single-cell invariance properties has remained limited, with only a few classic examples discovered using parametric or semantically meaningful stimuli that are heavily biased (Hubel and Wiesel, 1962; El-Shamayleh and Pasupathy, 2016; Quiroga et al., 2005). Recent advances in building digital twins of the brain and using non-parametric deep learning-based image synthesis have opened new avenues for finding the preferred stimuli of visual neurons (Walker et al., 2019; Bashivan et al., 2019; Ponce et al., 2019). Yet, most efforts in this direction have focused primarily on characterizing neuronal *selectivity* rather than invariance, and have emphasized feature visualization without further interpreting the invariant response properties of neurons.

In this study, we extended previous work combining large-scale neuronal recording and deep neural networks for the study of neuronal selectivity (Walker et al., 2019) to the invariance problem. Modifying a diverse feature visualization approach previously developed in ANNs (Cadena et al., 2018), we synthesized Diverse Exciting Inputs (DEIs) for individual neurons in mouse V1 Layer 2/3. These highly activating images reveal novel natural-occurring invariance that extend beyond the classical phase invariance described by Hubel and Wiesel (Hubel and Wiesel, 1962).

Particularly, we found a novel bipartite invariance in mouse V1 neurons: one RF subfield preferred a fixed spatial pattern, while the other preferred random crops from a texture image. While previous studies suggested a bimodal distribution of phase invariance corresponding to simple and complex cells (Niell and Stryker, 2008), our findings reveal that bipartite invariance in mouse V1 L2/3 cannot be explained as a continuum between these classical models or as a combination of overlapping simple and complex cells. A null model that parameterizes DEIs as a learned weighted summation of two fully overlapping subfields fail to produce DEIs as diverse and highly activating as bipartite DEIs, demonstrating that bipartite structure is necessary. Moreover, shift invariance is primarily localized to the variable subfield, as introducing it in the fixed subfield reduces responses.

Additionally, we show that bipartite structure cannot be explained by classical center-surround interactions, consistent with findings from (Fu et al., 2024), which demonstrated that MEIs correspond well to classical RF measurements, while extra-classical surround modulation extends far beyond the MEI. In particular, we observed no consistent spatial relationship between the minimum response field (MRF) and either the fixed or variable subfields, further ruling out center-surround mechanisms as an explanation for bipartite invariance.

While we have focused primarily on shift invariance, it is unlikely to be the only type of invariance existing in mouse visual system. As an initial effort to parameterize novel empirical invariances, it is also worth acknowledging that our partial-texture model proposes a simple hypothesis of a binary division of the presence and absence of shift invariance in the RF without considering more complicated scenarios such as nonlinear cross-subfield interactions. We also acknowledge that parameterizing complex invariances (e.g., 3D pose) for higher visual area remains challenging. Future studies using photo-realistic rendering engines with explicitly defined latent variables and image transformation will allow for a more generalized parameterization of invariances in a well-defined latent space, including 3D pose and other complex transformations. Nonetheless, we believe the novel bipartite invariance can be of great use as a computational principle for future designs of biologically-plausible or brain-inspired computer vision systems (Dapello et al., 2020). On the other hand, it can also serve as an empirical test for theoretically driven (Maruyama et al., 1992; Poggio and Girosi, 1990; Anselmi et al., 2015) or data-driven models (Schrimpf et al., 2018; Cadena et al., 2019; Schrimpf et al., 2020; Wang and Ponce, 2022; Willeke et al., 2022) that aim to explain and predict neuronal responses in the visual system.

In our study, the two RF subfields of the bipartite structure exhibit distinct characteristics, differing in both level of invariance and preferred spatial frequency. This property bears a striking resemblance to “high-low frequency detectors” observed in artificial neural networks, which detect low-frequency patterns on one side of their receptive field and high-frequency patterns on the other (Schubert et al., 2021). This parallel suggests that bipartite invariance with frequency bias may be a common feature shared between biological and artificial visual systems for boundary detection. Schubert et al. (2021) proposed these units encode natural boundaries defined by spatial frequency variation, aligning with our finding that V1 Layer 2/3 neurons prefer natural image patches containing object boundaries. While both classical simple/complex cells and V1 neurons are strongly activated by object boundaries in natural images (Biederman, 1987; von der Heydt and Peterhans, 1989; Gilbert and Wiesel, 1990; Kapadia et al., 1995), they prefer distinct boundary-constructing features. Classical cells prefer boundaries formed by rapid luminance changes (Hubel and Wiesel, 1962; Zhan and Baker Jr, 2006), whereas V1 neurons prefer boundaries defined by spatial frequency and orientation heterogeneity.

Our findings further complement behavioral studies showing that mice are able to use texture-based cues for segmentation (Kirchberger et al., 2021; Schnabel et al., 2018). While previous research emphasized boundaries constructed by orientation or phase differences (Schnabel et al., 2018), our results indicate that spatial frequency variation could serve as an additional second-order visual cue for boundary detection in the mouse visual system. Notably, humans also use spatial frequency as a visual cue for object/background assignment, often perceiving higher frequency regions as objects (Klymenko and Weisstein, 1986; Klymenko et al., 1989). This preference mirrors that of V1 neurons, suggesting potential common strategies for object/background segmentation between mice and primates.

The brain’s ability to generalize has long been hypothesized to rely on a cortical hierarchy where neurons tuned to simpler features combine to build complex functional invariance (Riesenhuber and Poggio, 1999; Serre and Riesenhuber, 2004; DiCarlo and Cox, 2007; DiCarlo et al., 2012; Poggio and Bizzi, 2004). This concept originates from Hubel & Wiesel’s model of complex cells achieving phase invariance by integrating inputs from simple cells (Hubel and Wiesel, 1962). Despite its longevity, empirical validation has been challenging due to difficulties in simultaneously studying physiology and wiring at the single-cell level (Lichtman and Denk, 2011; Briggman and Bock, 2012), and accurately modeling and measuring functional invariance (DiCarlo et al., 2012; Rust and DiCarlo, 2010).

Our study overcomes these challenges by utilizing the MICrONS dataset, the largest functionally-imaged EM dataset to date (MICrONS Consortium et al., 2021), and a digital twin model from a state-of-the-art foundation model for mouse visual cortex (Wang et al., 2023). This approach enabled us to identify concrete evidence supporting this decades-old hypothesis at the individual neuron level. We uncovered two key findings supporting hierarchical organization within V1 Layer 2/3:

1. Postsynaptic neurons exhibit higher level of functional invariance than their presynaptic counterparts.
2. Lower invariance presynaptic neurons form exponentially more synapses per unit of axon-dendrite co-traveling distance.

These findings provide the first evidence of a functional invariance hierarchy at the individual neuron level within the same cortical area and layer, mediated by horizontal connections. This contrasts with previous models like HMAX (Riesenhuber and Poggio, 1999; Serre and Riesenhuber, 2004; Poggio and Bizzi, 2004), which focused on hierarchies between cortical areas. Our results indicate that this hierarchical organization is regulated at both individual synapse and the presynaptic neuron level, revealing previously unrecognized computational flexibility. These findings align with studies demonstrating the importance of lateral connections for invariant object representation (Keck and Lücke, 2010; Crutcher, 2024), and could guide the design of more biologically plausible AI systems. Although the MEIs and DEIs synthesized from the dynamic-static digital twin elicited somewhat weaker responses compared to those from our standard static model, this imperfect replication did not impede our ability to extract meaningful insights and reveal significant relationships between neuronal structure and function.

As connectomics proofreading for the MICrONS dataset continues (MICrONS Consortium et al., 2021; Celii et al., 2023), we anticipate to gain a more comprehensive understanding of invariant object recognition mechanisms. Future access to multiple presynaptic neurons per postsynaptic neuron will allow detailed examination of how presynaptic inputs shape the bipartite properties of postsynaptic neurons. We also aim to extend our analysis to higher cortical areas to explore functional invariance build-up across the visual processing hierarchy. Future studies using more sophisticated models or direct *in vivo* measurements could further validate and refine these findings, potentially uncovering additional insights in cortical processing organization. Moreover, it would be important to compare the extent to which our current findings generalize to other species, such as non-human primates, where there are some similarities but also important differences in the functional organization of V1.

Overall, our work represents a significant advancement in understanding cortical processing and neuronal tuning. By developing a novel framework that combines large-scale neuronal recordings with advanced deep neural network techniques, we have enabled the systematic characterization of single-neuron invariances. The discovery of bipartite invariance in mouse V1 challenges long-held assumptions about receptive field homogeneity and offers new insights into natural image segmentation. Furthermore, leveraging the MICrONS dataset has allowed us to provide the first empirical evidence for a functional invariance hierarchy within V1 Layer 2/3, validating and extending theoretical models of cortical organization. The flexibility of our paradigm opens up possibilities for exploring neuronal invariances across various cortical regions, sensory modalities, and species, promising to illuminate the complex nature of neuronal coding more broadly, and potentially informing the development of more sophisticated, biologically-plausible artificial intelligence systems.

## ACKNOWLEDGEMENTS

The authors thank David Markowitz, the IARPA MICrONS Program Manager, who coordinated this work during all three phases of the MICrONS program. We thank IARPA program managers Jacob Vogelstein and David Markowitz for co-developing the MICrONS program. We thank Jennifer Wang, IARPA SETA for her assistance. The work was supported by the Intelligence Advanced Research Projects Activity (IARPA) via Department of Interior/ Interior Business Center (DoI/IBC) contract numbers D16PC00003, D16PC00004, and D16PC0005. The U.S. Government is authorized to reproduce and distribute reprints for Governmental purposes notwithstanding any copyright annotation thereon. XP acknowledges support from NSF CAREER grant IOS-1552868 and from the funds provided by the National Science Foundation and by DoD OUSD (R&E) under Cooperative Agreement PHY-2229929 (The NSF AI Institute for Artificial and Natural Intelligence, ARNI). ZhuD, SP, XP, and AST acknowledge support from NSF NeuroNex grant 1707400. AST, XP, KJ, and JR are supported by RF1 MH130416. AST also acknowledges support from the National Institute of Mental Health and National Institute of Neurological Disorders And Stroke under Award Number U19MH114830 and National Eye Institute award numbers R01 EY026927 and Core Grant for Vision Research T32-EY-002520-37. MD is supported by the European Union’s Horizon 2020 research and innovation program under the Marie Skłodowska-Curie grant agreement No. 101025482. EF is supported by a European Research Council (ERC) grant (ERC-2022-STG, NEURACT, Grant agreement No: 101076710) and by the Hellenic Foundation for Research and Innovation (HFRI) under the 2nd Call for HFRI Research Projects to Support Faculty Members and Researchers with Grant agreement No. 4049. LB, JA are supported by EU Horizon 2020 Maria SklodowskaCurie grant agreement No 861423 and ERDF-Project Brain dynamics, No. CZ.02.01.01\00\22_008\0004643. SS is supported by funding from the Amaranth Foundation. Disclaimer: The views and conclusions contained herein are those of the authors and should not be interpreted as necessarily representing the official policies or endorsements, either expressed or implied, of IARPA, DoI/IBC, or the U.S. Government. This work was also supported by the European Research Council (ERC) under the European Union’s Horizon Europe research and innovation programme (Grant agreement No. 101041669) as well as the Deutsche Forschungsgemeinschaft (DFG, German Research Foundation), project ID 432680300 (SFB 1456, project B05).

## AUTHOR CONTRIBUTIONS

We adopted the following contribution categories from CRediT (Contributor Roles Taxonomy). Authors within each category are sorted in the same order as in the author list.

**Conceptualization**: ZhiD, DTT, ZhuD, PF, EC, AC, StP, JF, SAC, FA, EF, SaP, EdYW, JR, FHS, ASE, KF, XP, AST

**Investigation**: ZhiD, DTT, KP, ZhuD, RF, LN, LB, MD, TM, AST

**Methodology**: ZhiD, DTT, ZhuD, EC, AC, LB, ErYW, JF, TM, AE, KW, EF, AST

**Data curation**: ZhiD, DTT, ZhuD, PF, MD, ErYW, StP, CP

**Formal Analysis**: ZhiD, DTT

**Project administration**: ZhiD, DTT, AST

**Supervision**: JR, FHS, ASE, KF, XP, AST

**Funding acquisition**: SS, KF, JR, XP, AST

**Resources**: KP, RF, LN, MD, TM, JR, KF, XP, AST

**Software**: ZhiD, DTT, ZhuD, EC, LB, ErYW, CP, AE, KW, SaP

**Validation**: ZhiD, DTT, KP, ZhuD, RF, LN, PF, ErYW, StP, TM

**Visualization**: ZhiD, DTT, PF, StP

**Writing – original draft**: ZhiD, DTT, AST

**Writing – review & editing**: ZhiD, DTT, ZhuD, PF, LB, MD, StP, FA, SS, SaP, JR, FHS, ASE, KF, XP, AST

## Methods

### Neurophysiological experiments

#### Two-photon calcium imaging

The following procedures were approved by the Institutional Animal Care and Use Committee of Baylor College of Medicine. 17 mice (*Mus musculus*: 9 male, 8 female) aged from 6 to 17 weeks, expressing GCaMP6s in excitatory neurons via Slc17a7-Cre and Ai162 transgenic lines (stock nos. 023527 and 031562, respectively; The Jackson Laboratory) were selected for experiments. The mice were anesthetized and a 4-mm craniotomy was made over the visual cortex of the right hemisphere as described previously (Reimer et al., 2014; Froudarakis et al., 2014). For functional imaging, mice were head-mounted above a cylindrical treadmill and calcium imaging was performed using a Chameleon Ti-Sapphire laser (Coherent) tuned to 920 nm and a large field-of-view meso-scope equipped with a custom objective (0.6 numerical aperture, 21 mm focal length) (Sofroniew et al., 2016). Laser power at the cortical surface was kept between 13.18 mW and 21.96 mW and maximum laser output of 61 mW was used at 245 *µm* from the surface.

We also recorded the rostro-caudal treadmill movement as well as the pupil dilation and movement. The treadmill movement was measured via a rotary optical encoder with a resolution of 8,000 pulses per revolution and was recorded at approximately 100 Hz in order to extract locomotion velocity. Light diffusing from the laser during scanning through the pupil was used to capture pupil diameter and eye movements. The images of the left eye were reflected through a hot mirror and captured with a GigE CMOS camera (Genie Nano C1920M; Teledyne Dalsa) at 20 fps with a resolution of 246-384 pixels × 299-488 pixels. A DeepLabCut model (Mathis et al., 2018) was trained on 17 manually labeled samples from 11 animals to label each frame of the compressed eye video with 8 eyelid points and 8 pupil points at cardinal and inter-cardinal positions. Pupil points with high likelihood were fit with the smallest enclosing circle, and the radius and center of this circle was extracted.

We delineated visual areas by manually annotating the retinotopic map generated by pixel-wise response to a drifting bar stimulus across a 4,000 × 3,600*µm*^2^ region of interest (0.2px*µm*^−1^) at 200 *µ*m depth from the cortical surface. The imaging site in V1 was chosen to minimize blood vessel occlusion and maximize stability. Imaging was performed using a remote objective to sequentially collect ten 630 × 630*µm*^2^ fields per frame at 0.4 px*µm*^−1^ xy resolution at approximately 8 Hz for all scans. We allowed only 5*µ*m spacing across depths to achieve dense imaging coverage of a 630 × 630 × 45*µm*^3^ *xyz* volume. The most superficial plane positioned in L2/3 was around 200*µ*m from the surface of the cortex. Thanks to our dense sampling, cells in the imaged volume were heavily over-sampled, often appearing in at least 2 or more imaging planes. This allowed matching across days with 2.5 *±* 2.6*µm* vertical distance between masks (see details below). We performed raster and motion correction on the imaging data and then deployed CNMF algorithm (Pnevmatikakis et al., 2016) implemented by the CaImAn pipeline (Giovannucci et al., 2019) to segment and deconvolve the raw fluorescence traces. Additionally, cells were selected by a classifier (Giovannucci et al., 2019) trained to detect somata based on the segmented cell masks to result in 7,049–8,238 soma masks per scan. The full two-photon imaging processing pipeline is available at (https://github.com/cajal/pipeline).

We did not employ any statistical methods to predetermine sample sizes but our sample sizes are similar to those reported in previous publications. Data collection and analysis were not performed blind to the conditions of the experiments but no animal or collected data point was excluded for any analysis performed.

#### Electrophysiological recording

Six mice (*Mus musculus*: 2 male, 4 female) aged from 14 to 27 weeks were selected for experiments, with 2 females expressing GCaMP6s in excitatory neurons via Slc17a7-Cre and Ai162 transgenic lines (stock nos. 023527 and 031562, respectively; The Jackson Laboratory) and the rest being C57BL/6J wildtype (stock no. 000664; The Jackson Laboratory). We performed acute recordings using Neuropixels probes 1.0 in awake, head-fixed mice according to (Jun et al., 2017). In brief, animals were implanted with a headpost and habituated to the experimental setup (head fixation on a treadmill) after recovery. On the recording day, the animals were briefly anaesthetized with isoflourane and a 1-mm craniotomy was made above visual cortex (approximately 2.9 mm lateral to the midline sagittal suture and anterior to the lambda suture) (Froudarakis et al., 2014). The animals were then transferred to the experimental setup and allowed to recover from anaesthesia. Location of probe insertion was chosen according to stereotaxic coordinates for targeting V1 using Pinpoint (Birman et al., 2023), with all penetrations ranging from 600*µm* to 1100*µm* on the anteroposterior axis, 2900*µm* to 3500*µm* on the mediolateral axis, and at an angle of 55° or 60° with respect to the ventrodorsal axis. One probe was smoothly lowered through the craniotomy to the final depth according to the trajectory planning with Pinpoint (Birman et al., 2023) to cover the whole cortex (covering 1800 − 2000*µm* of the probe) and allowed to settle for approximately 20 minutes before any recording. Visual area segmentation were performed by mapping the reversals of the retinotopy based on the RF progression along the probe as described previously (Tafazoli et al., 2017). Neuronal activity recordings were made with custom-written software in LabView and then automatically spike sorted with the Kilosort3 spike sorting software (Pachitariu et al., 2023). An external infrared light was used as the light source for capturing pupil diameter and eye movements. A DeepLabCut model (Mathis et al., 2018) was trained on 13 manually labeled samples from 4 animals to label each frame of the compressed eye video with 8 eyelid points and 8 pupil points at cardinal and intercardinal positions. Pupil location and radius was extracted following the identical procedure described in *Two-photon calcium imaging*. From a total of 9 recording sessions, 3283 neurons were detected by the spike-sorting algorithm (136–547 per session), with 364 neurons from V1 Layer 2/3 (12–95 per session). All V1 Layer 2/3 neurons were compiled together for predictive model training, and then neurons classified as “single units” or “multi-unit activity” (MUA) were used separately for downstream analysis. We evaluated the level of unit contamination using inter-spike-interval (ISI) violations, following the approach introduced by Hill et al. (2011). This metric represents the relative firing rate of hypothetical contaminating sources that produce these violations, with higher ISI violations indicating greater level of contamination.

#### Visual stimuli presentation

Visual stimuli were presented 15 cm away from the left eye with a 25” LCD monitor (31.8 x 56.5 cm, ASUS PB258Q) at a resolution 1080 ×1920 pixels and refresh rate of 60 Hz. We positioned the monitor so that it was centered on and perpendicular to the surface of the eye at the closest point, corresponding to a visual angle of 2.2° /cm on the monitor. In order to estimate the luminance level of the stimuli presented on the monitor, we taped a photodiote at the top left corner of the monitor and recorded its voltage during stimulus presentation, which is approximated linearly correlated with the monitor luminance. The conversion between photodiode voltage and luminance was estimated from luminance measurements from a luminance meter (LS-100 Konica Minolta) for 16 equidistant pixel values ranging from 0 to 255 while simultaneously recording photodiode voltage. Since the relationship between photodiode voltage and luminance is usually stable, we only perform such measurement every a few months. In the beginning of every experimental session, we computed the gamma between pixel intensity and photodiode voltage by measuring photodiode voltage at 52 equidistant pixel values ranging from 0 to 255; then we further interpolated the corresponding luminance at each pixel intensity. For closed-loop experiments, the pixel-luminance interpolation computed on day 1 was used throughout the loop. All stimuli used in the current study were presented at gamma value ranging from 1.59 to 1.77 and monitor luminance ranging from 0.07 *±* 0.16*cd/m*^2^ to 9.58 *±* 0.65*cd/m*^2^.

### Presentation of natural stimuli

To fit neurons’ responses, 5,100 natural images from ImageNet (ILSVRC2012) were cropped to fit a 16:9 monitor aspect ratio and converted to gray scale. To collect data for training a predictive model of the brain, we showed 5,000 unique images as well as 100 additional images repeated 10 times each. This set of 100 images were shown in every scan for evaluating cell response reliability within and between scans. Each image was presented on the monitor for 500 ms followed by a blank screen lasting between 300 and 500 ms, sampled uniformly. Identical natural stimuli were used for two-photon imaging and electrophysiological experiments. To maintain the animal’s alertness throughout each scan, we interspersed an additional set of six brief movie clips at regular intervals.

### Neuronal data processing and predictive modeling

#### Preprocessing of neuronal responses and behavioral data

Neuronal responses were deconvolved using constrained nonnegative calcium deconvolution and then accumulated between 50 and 550ms after stimulus onset of each trial using a Hamming window. All the segmented neuronal masks from each individual scan were used for model trainig, including duplicates resulting from dense imaging. The corresponding pupil movement and treadmill velocity for each trial were also extracted and integrated using the same Hamming window. Each dataset consists of 4500 and 500 unique images for training and validation, respectively; an additional set of 100 images presented with 10 repeats was used for model evaluation. The original stimuli presented to the animals were isotropically downsampled to 64×36 pixels for model training. For day 1 model training scans, input images, neuronal responses, and behavioral traces were normalized (z-scored for input images and divided by standard deviation for the rest) across the training set during model training and evaluation. Trials with invalid behavioral data (0.8 *±* 1.2%) were excluded from model training. For closed-loop verification scans, neuronal responses and behavioral traces were normalized across all trials.

#### Predictive model architecture and model training

We followed the same network architecture and training procedure as described previously (Sinz et al., 2018; Walker et al., 2019). Each model comprises: a shared non-linear core (157, 920 parameters), neuron-specific linear readouts at six different spatial scales (579 parameters per neuron), a behavioral modulator (150 shared parameters across all neurons and 84, 007 parameters per neuron), and a pupil position shifter network shared across all neurons (57 parameters). The common core is a 3-layer CNN with full skip connections. Each layer contains a convolutional layer with no bias, followed by batch normalization, and an exponential linear unit (ELU) nonlinearity. The readout models the neuronal response as an affine function of the core outputs followed by ELU nonlinearity and an offset of 1 to guarantee positiveness. Additionally, we model the location of a neuron’s RF with a spatial transformer layer reading from a single grid point that extracts the feature vector from the same location at different scales of the downsampled feature outputs. The modulator computes a gain factor for each neuron that simply scales the output of the readout layer using a two-layer fully connected multi-layer perceptron (MLP) with rectified linear unit (RELU) nonlinearity and a shifted exponential non-linearity to ensure positive outputs. Finally, because training mice to fixate their gaze is impractical, we estimated the trial-by-trial RF displacement shared across all neurons using a shifter network composed of a three-layer MLP with a tanh nonlinearity. For all model training, we adhered to the methodology outlined in Walker et al. (2019), training four instances of the same network with different initializations by minimizing the Poisson los 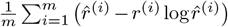 where *m* denotes the number of neurons, 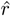 the predicted neuronal response, and *r* the observed response.

### Evaluation of model performance and neuronal reliability

Predictions from all four models are averaged for model benchmarking and image generation. We computed the model performance *CC*_*abs*_ for each neuron on the same held-out data as the correlation between the model predicted response 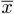 and the recorded responses 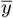 averaged across 10 repetitions:

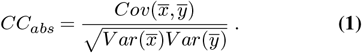

. To assess reliability of neuronal responses, we computed *CC*_*max*_ (Schoppe et al., 2016) as

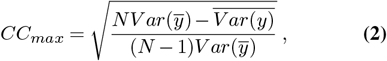

where *y* is the *in vivo* responses, and *N* is the number of trials. This metric captures the consistency of neuronal responses to identical visual stimuli in held-out data, serving as an upper bound for our model’s potential performance. We then estimated the normalized correlation coefficient (*CC*_*norm*_) (Schoppe et al., 2016) as the fraction of variation in neuronal responses to identical stimuli accounted for by the model prediction:

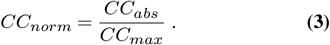

### Non-parametric synthesis of optimal stimuli and controls

#### Neuron selection

This section describes neuron section for stimulus synthesis for 14 out of the 17 mice used for all experiments except for dynamic-static model validation. We first excluded neuronal masks within 10*µ*m from the edge of the imaging volume, and then ranked the remaining masks based on descending model predictive accuracy. To avoid duplicated neurons, we started from the lowest-ranked neuron and iteratively added neurons such that they are at least 25*µ*m apart and have functional correlation *<* 0.4 with all neurons selected. This filtering left us with 2,081–2,676 unique neurons for each scan. We restricted all analyses to neurons that exhibit reasonable levels of response reliability as well as model predictive accuracy. We evaluated neuronal reliability using “oracle score” (Walker et al., 2019) (a metric highly correlated with *CC*_*max*_ (Schoppe et al., 2016), Pearson correlation *r* = 0.9) for each neuron by correlating its leave-one-out mean response with that of the remaining trial across 100 images in the held-out test set. For synthetic stimulus generation, we applied hard thresholds on oracle score and model test correlation to include 19.9% of the population for mouse 1 and 79.0 *±* 0.5% of the population for mice 2–14.

#### Generation of Most Exciting Input (MEI)

For each individual neuron, we adapted the activation maximization procedure described earlier (Walker et al., 2019) to find the stimulus that optimally drive each individual neuron. Starting with Gaussian white noise, we iteratively refined the image by adding the gradient of the target neuron’s predicted response using a SGD optimizer with learning rate of 1.0 for 1000 iterations. To mitigate high-frequency artifacts in image synthesis, we applied a Gaussian filter (*‡* = 1.0) to smoothen the gradient at every optimization step. To determine the appropriate root mean square (RMS) contrast value for our synthetic stimuli, we conducted a pilot analysis in which we aggregated MEI masks from thousands of neurons into an average mask and measured mean contrast within this average mask across all the training set natural images presented. To prevent saturation and ensure that the synthetic stimuli remain within the well-trained contrast domain of the natural images used during model training, we standardized the image to a fixed mean of 0 and RMS contrast of 0.25—the value obtained from the pilot analysis—following each gradient ascent step.

We computed a weighted mask for each MEI to capture the region containing the vast majority of the variance in the MEI image. We computed a pixel-wise z-score on the MEI and thresholded at z-score above 1.5 to identify the highly contributing pixels. Then we closed small holes/gaps using binary closing, searched for the largest connected region to create a binary mask *M* where *M* = 1 if the pixel is in the largest region identified. Then, a convex hull was calculated using the identified pixels. Lastly, to avoid edge artifacts, we smoothed out the mask using a Gaussian filter with *σ* = 1.5 to avoid potential edge effects.

#### Generation of Diverse Exciting Inputs (DEIs)

We modified procedures described previously (Cadena et al., 2018) to optimize DEIs. For each individual neuron, we synthesized a set of images initiating from MEI that preserve high activation while differ as much as possible from each other. To optimize this set, we initiated from 20 instances of the target neuron’s MEI with different additive Gaussian white noises *MEI* + *σ*_*i*_ = *I*_*i*_ where 1 ≤ i ≤ 20 and iteratively minimize the loss:

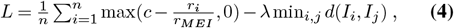

where *r*_*i*_ and *r*_*MEI*_ are the model predicted response to DEI_*i*_ and MEI, *c* is the minimum activation relative to *r*_*MEI*_ that we target each DEIs for, *d*(*I*_*i*_, *I*_*j*_) is the Euclidean distance in pixel space between DEI_*i*_ and DEI_*j*_ measured within the MEI mask (i.e. the neuron’s receptive field). The first term encourages all DEIs to achieve high activation, while the second term maximizes the minimum pair-wise distance among DEIs. Specifically, we required each DEI to evoke at least 85% of their corresponding MEI (*c* = 0.85). This threshold was selected based on the previous finding that additional decrease in target response leads to marginal gain in minimum pair-wise distance among DEIs for simulated complex cells (Cadena et al., 2018). Note that the minimum, instead of average distance, is used in the second term to avoid solutions that form the set of DEIs into clusters by pushing apart the most similar pair of DEIs at every iteration. We employed the same gradient blurring and post-gradient image standardization as in MEI optimization. We optimized the DEI set for 3000 iterations with a learning rate of 1000 for the first 2000 iterations and decayed to 100 for the last 1000 iterations. This learning rate decay helped to further mitigate the occurrence of high-frequency artifacts. We performed the optimization for every target neuron with a series of diversity regularization hyper-parameter */*, densely sampled from 1×10^−4^ to 5×10^−2^. For each neuron, the set optimized using the largest *λ* that preserved minimal response greater than 85% of the MEI response was selected as the DEIs and used for downstream analyses and experiments.

### Diversity index

To quantify the diversity level of each set of DEIs we derived a diversity index based on the average pair-wise Euclidean distance of the DEIs. To position this metric on a meaningful spectrum with interpretable reference points, we estimated diversity levels of idealized simple and complex cells (see details in *Simulation of simple and complex cells*). Particularly, we estimated the lower/upper bounds (*d*_lower_ and *d*_upper_) as the median average pairwise Euclidean distance of DEIs from a population of noiseless simple/complex cells, respectively. We performed an exhaustive search through the Gabor parameter space to identify their DEIs. When standardized with a fixed mean and RMS contrast, DEIs from idealized simple cells have the same average pairwise Euclidean distance regardless of the underlying Gabor parameters. Similarly, idealized complex cells with different Gabor parameters have identical yet higher average pairwise Euclidean distance. For each real neuron, a diversity index (*D*) is calculated for each mouse V1 neuron *i* based on the average pairwise Euclidean distance of its DEIs *d*^(*i*)^ as

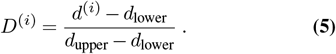

### Natural image and synthesized controls for the invariance manifold

To evaluate the specificity of the invariance manifold represented by the DEIs, we designed two types of control stimuli: natural image controls and synthesized controls. Both controls were strictly closer to the MEI than all the DEIs, as quantified by the corresponding metric used in DEI generation. For each neuron, we first computed the minimum distance from the DEIs to the MEI within the MEI mask, denoted as *d*_*target*_, which served as the distance budget for control image selection or synthesis. For natural image controls, we searched through more than 40 millions of natural image patches to identify those with distances from the MEI between 80% and 100% of *d*_*target*_. The synthesized controls were generated using a modified version of our DEI synthesis objective (Eqn. 4), where the first term aimed to match the distance from control images to the MEI to *d*_*target*_ rather than encouraging high activation:

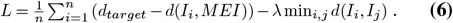

For both control types, we created 20 different images per neuron and presented each once *in vivo* during closed-loop experiments.

#### Generalization of DEIs

To test the generalizability of our DEI synthesis methodology, we modified key components of the synthesis pipeline and compared the resultant DEIs:

1. Image initialization: DEIs were initiated with full-field random white noise instead of a combination of the Most Exciting Inputs (MEI) and random white noises.
2. Model initialization: The in silico model ensemble was trained from scratch using a different random initialization seed.
3. Individual model synthesis: DEIs were generated using the response of a single model from the ensemble rather than the average response of four models.
4. Diversity metric: DEIs were synthesized with diversity measured in neuronal representational space instead of pixel space, as detailed in *Generation of DEIs in neuronal representational space*.
5. Synthesis methodology: DEIs were generated using an alternative approach described in *Generation of DEIs with implicit neural representation model and contrastive regularization*.
6. Model architecture: DEIs were produced using the distinct model architecture outlined in Willeke et al. (2022).

We computed representational similarity (as detailed in *Representational similarity*) and average pairwise Euclidean distance of DEIs generated from various conditions to assess the robustness of DEIs.

#### Selection of natural DEIs

For each neuron, we searched through 41 million ImageNet image patches *in silico* to identify natural crops that elicited activations equal to or greater than 85% of the MEI response (i.e. DEI-like activation). To mitigate the effect of contrast difference at the edges of masked natural crops and MEIs, we refined MEI masks by shrinking them until the activation of masked MEI dropped below 95% of the original MEI activation, following the approach of Walker et al. (2019). Each crop was then masked using the refined MEI mask, and its mean and RMS contrast were adjusted to match those of the MEI. For neurons with at least 20 highly activating crops, we selected 20 natural DEIs—matching the number of synthesized DEIs per neuron—by greedily maximizing their minimum pairwise distance, mirroring the DEI synthesis procedure. These selected images are denoted as “natural DEIs”.

#### Generation of DEIs in neuronal representational space

We utilized the same loss function as in 4 but quantified pairwise DEI diversity *d*(*DEI*_*i*_,*DEI*_*j*_) as the negative Pearson correlation between model-predicted neuronal population response vectors *r*_*i*_ and *r*_*j*_ to *DEI*_*i*_ and *DEI*_*j*_:

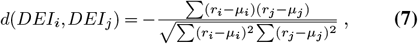

where *µ*_*i*_, *µ*_*i*_ represent the mean neuronal population responses. To compute these population responses, we aligned all neurons’ receptive field (RF) centers with that of the target neuron for which the DEIs were being optimized. We refer to thee DEIs generated through this method as “neuronal-space DEIs”.

#### Generation of DEIs from an implicit neural representation model with contrastive regularization

Following the approach detailed in Baroni et al. (2023), we used an implicit neural representation model (INRM) to map from a low-dimensional periodic latent space (1D or 2D) to a manifold in image space representing the invariant transformations of a given neuron. The INRM we used was a fully connected feed-forward neural network mapping from pixel coordinates and latent inputs to pixel values. Our model consists of 4 layers of 50 hidden nodes, followed by an hyperbolic tangent nonlinearity and a sigmoid function as final non-linearity. We used positional encoding on both the latent space and the co-ordinate space. Each latent input could be mapped to one image and changing the latent input corresponded to moving along one invariant dimension of the neuron. The images were standardized to a fixed mean and RMS contrast and clipped between values corresponding to the black and white pixels on the monitor before being passed to the digital twin to get the predicted response.

During training of an INRM, a jittering grid of uniformly distanced points was sampled from the latent space and mapped into a set of images. The training objective was composed of one activation term that maximizes the activation of the generated images and one contrastive term that encourages diversity across images and ensures smooth transitions in image space when navigating the latent space. Specifically, the contrastive regularization term achieved this by encouraging images corresponding to nearby points in latent space to have high cosine similarity and those corresponding to distant points in latent space to have low cosine similarity. The contrastive regularization temperature (Baroni et al., 2023) was set to 0.3. The latent space grid size was 20 points in 1D and 7 points in 2D per dimension. The neighboring radius, which determined close-by points in the latent space, was set to 10% of the grid in the 1D case and 20% of the grid in the 2D. We used an Adam optimizer with learning rate of 0.001 to optimized the INRM weights. After a minimum of 500 weight updates, the regularization strength was decreased by a factor of 0.8 every time the activity stopped increasing (initial strength of 2.0, one check every 50 steps with patience of 5). Training was stopped when the resultant images achieved an average response larger than 85% of the MEI response and a minimum response larger than 75% of the MEI response. To avoid image artifacts, gradients were Gaussian blurred (*σ* = 1.0) and contrastive regularization was applied only on pixels within the MEI mask.

This method learns a continuous manifold of stimuli. In the 1D case, we sampled 20 DEIs corresponding to uniformly distant points in latent space. In the 2D case, since different latent dimensions could learn transformations associated with different image diversity, we obtained 20 DEIs by starting from an initial set of images corresponding to randomly sampled points in latent space and then optimizing them to maximize the minimum pairwise distance.

### Bipartite parameterization of DEIs

#### Bipartite model

We proposed a texture generative model to produce texture-based DEIs composed of two complementary subfields as follows:

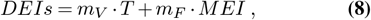

where the first term is the variable subfield randomly cropped from an optimized texture canvas *T* using a mask *m*_*V*_. The second term is a fixed subfield masked directly from the original MEI. This model could be reduced to a full-texture model to describe global shift invariance if the entire receptive field (i.e. the MEI mask *m*_*MEI*_) was used as *m*_*V*_. We generated the texture *T* following Cadena et al. (2018) by maximizing the average activation of randomly sampled crops from *T* using *m*_*V*_. We followed the same loss as in non-parametric DEIs generation 4 to jointly maximize the activation and diversity of DEIs with the same regularizations (i.e. Gaussian blurring on the gradient and learning rate decay) but in this case, DEIs were parameterized as in 8. The same post-gradient image standardization was applied on these parametric DEIs.

To ensure that *m*_*V*_ captures the region of the original non-parametric DEIs from which we observed the most diversity, we pre-computed a series of *m*_*V*_ s by varying the threshold on the pixel-wise variance across the DEIs. Specifically, starting from the pixels with the largest variance across DEIs, we kept expanding the *m*_*V*_ by requesting increasing fraction of the total variance from 0.2 to 0.6 within the variable subfield. The complement to *m*_*V*_ within *m*_*MEI*_ was used as the fixed subfield mask (*m*_*F*_). In general, the average predicted activation of the texture-based DEIs decreased as the size of *m*_*V*_ increased. For each set of texture-based DEIs resultant from each pair of subfield masks, we computed the harmonic mean between normalized activation and diversity index.

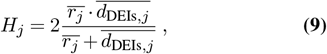

where 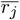 is the average activation and 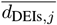 is the average pair-wise Euclidean distance, normalized by the maximal corresponding value across all different sets of DEIs using the series of *M*_*V*_ s, respectively. We denoted the set of texture-based DEIs with the maximum harmonic mean as the “partial-texture DEIs” (DEIs_partial_). The set of texture-based DEIs resultant from the full-texture model were denoted as the “full-texture DEIs” (DEIs_full_).

#### Bipartite invariance index

The “bipartite invariance index” was devised to summarize the extent of partial shift invariance exhibited by a neuron. Using the series of subfields masks and their corresponding texture-based DEIs as described above, we fitted a quadratic-smoothing spline to model the relationship between the *in silico* neuronal activation and the variable subfield size. To capture the full range of this relationship, we uniformly sampled the variable subfield size between 0 and 1 and evaluated the predicted response at each point using the fitted spline. Finally, we calculated the Area Under the Curve (AUC) of these predicted responses across the range of subfield sizes. This AUC value serves as our bipartite invariance index, encapsulating the neuron’s response profile across various subfield sizes and thus providing a comprehensive measure of its degree of partial shift invariance.

#### Preferred spatial frequency of bipartite RF subfields

Due to challenges of direct frequency analysis on small image windows (i.e. subfield masks), we employed an indirect comparative approach using two sets of images: 1) the full-field DEIs_partial_ 2) modified the full-field DEIs_partial_ where the content in the fixed subfield was substituted by random crops masked from the same texture optimized for variable subfield using the fixed subfield mask and standardized to have the same mean and RMS contrast as the original fixed subfield content. Both sets of images maintain the identical bipartite structure, differing only in the spatial content within the fixed subfield mask, thus providing an indirect but equitable way to compare frequency preferences of content from the two subfields. For each set of images, we first computed the radial power spectrum using 10 equally spaced bins. The resulting power spectra were then averaged to obtain the mean radial power spectrum, from which we estimated the median frequency using linear interpolation.

#### Necessity and specificity of two subfields in the bipartite RF

We masked out or swapped the content of either subfield to evaluate its necessity and specificity in eliciting higher neuronal responses, respectively. When isolating a subfield, we prioritized maintaining the integrity of its pixels. This was achieved by computing a binary mask for the subfield and smoothing the edge only strictly outside this mask. As a result, some parts of the complementary subfield remained visible in the stimuli, potentially leading to an underestimation of the complementary subfield’s necessity. Consequently, parts of the complementary subfield were preserved in the visual stimuli, making the measured necessity of the complementary subfield an underestimate of the actual effect. For the specificity assessment, we either replaced the fixed subfields with different random natural image crops or the variable subfields by random crops masked from different random neurons’ optimized textures.

### Controls for bipartite RF structure

#### Control parameterization: “two-variable-subfield DEIs”

To investigate whether the fixed subfield exhibits shift invariance and if DEIs can be better explained by a more complex model, we modified the bipartite model such that both subfields are treated as shift invariant, described by:

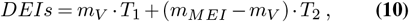

Here, the first term mirrors the bipartite parameterization, while the second term represents a second variable subfield, randomly cropped from a second optimized texture canvas *T*_2_. We followed the same procedure as in *Bipartite model* to sample a series of *m*_*V*_ and optimized *T*_1_ for each *m*_*V*_. Then we used the complementary subfield mask within the MEI mask *m*_*MEI*_ *−m*_*V*_ to optimize for a second texture canvas *T*_2_. We then combined crops masked from each subfield’s preferred texture to get sets of texture-based DEIs and selected the set with the highest harmonic mean of diversity and *in silico* activation as the “two-invatiant-field DEIs”.

#### Control parameterization: “no-spatial-division DEIs”

To assess the necessity of spatial division between the two subfields in the bipartite model, we developed an alternative parameterization that represents DEIs as a weighted summation of two fully overlapping subfields spanning the entire RF (estimated as the MEI mask *m*_*MEI*_): a fixed component directly from the MEI and a variable component cropped from a synthesized full-field texture. This model was implemented as

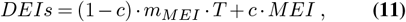

where the hyper-parameter *c* regulates the ratio between the variable and fixed subfield contribution. We uniformly sampled *c* between 0 and 1, where 0 signifies an ideal complex cell and 1 an ideal simple cell. For each *c*, we optimized the texture *T* following the same procedure as described in *Bipartite model*. We then combined the two overlapping subfields to get sets of texture-based DEIs and selected the set with the highest harmonic mean of diversity and *in silico* activation as “no-spatial-division DEIs”. We also fit the *in silico* activation as a quadratic-smoothing spline of the average pairwise Euclidean distance for each neuron (i.e. diversity). The spline fit was utilized to interpolate the diversity of these texture-based DEIs when their mean *in silico* activation was matched to that of the non-parametric DEIs. Similarly, we interpolated the mean *in silico* activation of these texture-based DEIs when their diversity was matched to that of the non-parametric DEIs. The same fitting and interpolation was also done for the biparitite model, allowing a direct comparison of how well these two parameterizations captured the diversity and *in silico* activation of the non-parametric DEIs.

#### Replication of bipartite structure using electrophysiological data

For Neuropixels electrophysiological data, we employed two strategies: training models from scratch, or utilizing a core pretrained on two-photon imaging data and then training the remaining components (including neuron-specific readouts, shifter, and modulator components) using Neuropixels data. The latter approach, particularly beneficial due to the limited number of neurons available from each Neuropixels recording session, improved the median normalized correlation coefficient (*CC*_*norm*_) from 0.64 to 0.73. We then generated MEIs, DEIs, texture-based DEIs following the same protocol as applied on the two-photon imaging models. For comparison of diversity and bipartite invariance indices between neurons from imaging and electrophysiological data, we applied identical functional thresholding (oracle score *>* 0.22 and model test correlation *>* 0.42, respectively, calculated as the median threshold from 14 mice used for two-photon closed-loop experiments) on both neuron populations to ensure fair comparison.

#### Comparison of bipartite RF structure and the Minimum Response Field (MRF)

To investigate the relationship between classical receptive fields estimated with the Minimum Response Field (MRF) Fu et al. (2024) and the bipartite RF structure, we presented sparse noise stimuli (Jones and Palmer, 1987b) prior to and after the natural image stimuli (detailed in *Presentation of natural stimuli*) in the same two-photon imaging scan. The stimuli comprised circular bright (pixel value=255) and dark (pixel value=0) dots, each spanning 7 degrees in visual angle, presented against a gray background (pixel value=128) on a 9 ×9 grid covering 40% of the monitor’s central area. Each dot was displayed for 250 ms per location with 16 repetitions (8 prior to and 8 after the natural stimuli). For both bright and dark dots, we aggregated neuronal responses from 50 to 300 ms post stimulus onset for each trial, creating separate ON and OFF maps. We then applied one-way ANOVA to these maps to identify neurons exhibiting significant spatial variation in their responses. The MRF was determined by aggregating ON and OFF maps, maximizing the averaged response per location, and fitting a 2D Gaussian. For quality control, we excluded neurons with extreme MRF sizes (bottom 5% and top 95%) and those with low Signal-to-Noise Ratio (SNR), calculated as 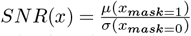, where the mask was obtained by thresholding the fitted Gaussian at 0.3. To evaluate the spatial relationship between the MRF and the bipartite structure, we calculated (1) the average pairwise distance between the MRF and each subfield across all pixels; (2) the overlap between the MRF and each subfield. To estimate the diameter of MEI, variable subfield, and MRF, we first binarized their masks (threshold = 0.3) and defined the diameter as the longest distance between points on the border of the mask.

#### Evaluation of pupil position uncertainty

To evaluate whether the bipartite structure is related to uncertainty in the trial-by-trial pupil shift predicted by the shifter network, we trained three additional versions of the model by sub-sampling trials based on the corresponding pupil movement: 1) minimal-movement model, where we removed trials with pupil distance from the mean position larger than 3 units in the eye camera coordinate system, which corresponded to approximately 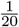 of the median MEI diameter (2.86∘ visual angle); the median percentage of trials included was 33.1%, 2) small-movement model, where trials with the bottom 50 percentile of pupil movements were included, and 3) large-movement model, where trials with the top 50 percentile of pupil movements were included. For each model we generated MEI, DEIs, and partial texture DEIs for each neuron.

### *In vivo* closed-loop verification of synthesized stimuli

#### Neuron selection

This section describes neuron section for stimulus synthesis for 14 out of the 17 mice used for all experiments except for dynamic-static model validation. For 9 out of 14 mice, we randomly selected neurons with relatively high level of invariance (detailed below); for the remaining 5 mice, we randomly selected neurons from all candidates that survived our oracle score and model performance criteria (see above). To remove the confounding effect of RF size on neurons’ invariance level, we fit a least squares regression from the MEI mask size to the diversity index computed from DEIs (see above) using 1000 random neurons compiled across 8 pilot datasets. For each neuron, the residual between the actual average DEIs pair-wise Euclidean distance and the predicted distance from the MEI mask size was calculated as its diversity residual. This diversity residual served as an size-independent evaluation of neuron’s invariance level. For each of the 9 mice, we randomly selected neurons from the top 50 percentile among all neurons with positive diversity residuals.

#### Presentation of synthetic stimuli

For all MEIs, DEIs, texture-based DEIs, and control stimuli, we masked the stimuli with the MEI mask and standardized all masked stimuli such that they have fixed value of mean (3.09*cd/m*^2^) and RMS contrast (0.25*cd/m*^2^) in the luminance space with only small amount of deviation due to clipping within the 8-bit range. The fixed mean and contrast valued were chosen to be close to the corresponding values of the training set while minimizing the amount of clipping when converting synthetic stimuli from the z-score space to the 8-bit image space. All pixels outside of the MEI mask are at 128, the same intensity as the blank screen in between consecutive trials. For each neuron, its MEI was presented for 20 repeats; for 2 mice, 10 random DEIs for each neuron werxe selected and each presented for 20 repeats, and the remaining mice, each of 20 DEIs or control stimuli for each neuron were presented once.

#### Monitor positioning across days

In order to optimize the monitor position for centered visual cortex stimulation, we mapped the aggregate receptive field of the scan field region of interest (ROI) using sparse noise stimuli comprised of bright (pixel value=255) and dark (pixel value=0) dots. We tiled the center of the screen in a 10 × 10 grid with single dots in random locations, with 10 repetitions of 200 ms presentation at each location. The RF was then estimated by averaging the calcium trace of an approximately 150 × 150*µm*^2^ window in the ROI from 0.5–1.5 s after stimulus onset across all repetitions of the stimulus for each location. The resulted two-dimensional map was fitted with an elliptic two-dimensional Gaussian to find a center. To keep a consistent monitor placement across all imaging sessions, we positioned the monitor such that the aggregate RF of ROI in the first session was placed at the center of the monitor and then fixed the monitor position across the subsequent sessions within a closed-loop experiment. An L-bracket on a six-dimensional arm was fitted to the corner of the monitor at its location in the first session and locked in position such that the monitor could be returned to the same position between scans and across imaging sessions.

#### Cell matching across days

In order to return to the same image site, the craniotomy window was leveled with regard to the objective with six d.f., five of which were locked between days. A structural 3D stack encompassing the volume was imaged at 0.8 × 0.8 × 1 *px*^3^*µm*^−3^ *xyz* resolution with 100 repeats. The stack contained two volumes each with 150 fields spanning from 50 µm above the most superficial scanning field to 50 *µm* below the deepest scanning field; each field was 500 × 800*µm*^2^, together tiling a 800 × 800*µm*^2^ field of view (300*µm* overlapped). This was used to register the scan average image into a shared *xyz* frame of reference between scans across days. To match cells across imaging scans, each two-dimensional scanning plane was registered to the three-dimensional stack through an affine transformation (with nine d.f.) to maximize the correlation between the average recorded plane and the extracted plan from the stack. Based on its estimated coordinates in the stack, each cell was matched to its closest cell across scans. To further evaluate the functional stability of neurons across scans, in each scan we included an identical set of 100 natural images with each repeated 10 times (referred as oracle images). For each pair of matched neurons from two different scans, we compute the correlation between their average-trial responses to the oracle images. In order to be included for downstream analyses, the matched cell pair need to (1) have an inter-cellular distance smaller than 10*µm*; (2) achieve a functional correlation equal to or greater than the top 1 percentile of correlation distribution between all unmatched cell pairs (estimated as 0.42); (3) survive manual curation of the matched pair’s physical appearance in the processed average frame. Among all closed-loop scans, 56 *±* 16% of closed-loop neurons per scan survived all three criteria.

### Analysis of *in vivo* neuronal responses

#### In vivo *response comparisons and statistical analysis*

Recorded responses were normalized across all presented images within each scan. The responses of the matched neurons were then averaged across either 20 repetitions of a single image for MEI or single repetitions of 20 different images for DEIs, texture-based DEIs, and different types of control stimuli. For individual neurons, the statistical significance of response differences across stimulus types was assessed using two-sided Welch’s *t*-tests. The overall response difference across stimulus types compiled across all neurons was assessed using two-sided Wilcoxon signed-rank tests.

Neuronal responses were normalized across all presented images within each scan. For matched neurons, we averaged responses across either 20 repetitions of a single image (for MEI) or single repetitions of 20 different images (for DEIs, texture-based DEIs, and various control stimuli). To assess the statistical significance of response differences, we employed two-sided Welch’s *t*-tests for individual neurons. For evaluating the overall difference in average responses across all neurons, we utilized two-sided Wilcoxon signed-rank tests.

#### Decoding DEIs from population responses

To assess whether differences across DEIs can be represented by V1 population responses, we identified the most dissimilar pair of DEIs for each neuron based on their corresponding diversity metric. We then presented each DEI of this pair (*DEI*_1_ and *DEI*_2_) 20 times. To quantify the neuronal discriminability between these DEIs, we implemented a 5-fold cross-validated logistic classification with L2 regularization on the V1 population responses. This classifier was trained to distinguish whether each single-trial population response originated from *DEI*_1_ or *DEI*_2_. The optimal regularization strength was determined empirically by fitting the logistic regression model on an independent pilot dataset.

#### Individual DEI response analysis

To compare the *in vivo* responses of individual DEIs to their corresponding MEI, we randomly selected 10 DEIs for each neuron and presented each DEI 20 times. Using a two-sided Welch’s test, we assessed whether responses to individual DEIs differed from 85% of their corresponding MEI response. To investigate whether the relative strength of DEI responses to MEI is influenced by whether single or multiple DEIs were presented, we implemented two bootstrapping strategies on the same data set: averaging across 20 repeats of the same DEI, and averaging across 20 single trials from different DEIs. For each bootstrap iteration, we estimated a robust linear coefficient between DEIs and MEI using the RANSAC algorithm (Fischler and Bolles, 1981). We then examined whether the difference in linear coefficients estimated from the two bootstrapping strategies differed from zero.

### *In silico* quantification and analysis

#### In silico *stimuli presentation*

To ensure the most reliable predictions from our model, we standardized all images to match the training set statistics before presenting them *in silico*. The training set images on average had approximately a mean of 0 and RMS contrast of 0.8 (after z-scoring) within the MEI mask. Through synthesizing MEIs with a series of full-field statistics constraints, we determined that full-field images with a mean of 0 and RMS contrast of 0.25 best replicated these statistics. Therefore, we adopted a uniform preprocessing procedure for all images: first, we applied the corresponding MEI mask to each image, then normalized the entire image to mean of 0 and RMS contrast of 0.25.

#### Simulation of simple and complex cells

The response of an idealized simple cell was modeled as convolution with a 2D Gabor filter followed by a rectified linear activation function (ReLU) (Jones and Palmer, 1987a). An idealized complex cell was formulated by the classical energy model (Adelson and Bergen, 1985), where the response was modeled as the square root of the summation of squared outputs to a quadrature pair of 2D Gabor filters. A Gabor image was generated as

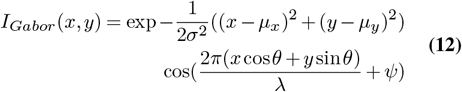

, where *µ*_*x*_ and *µ*_*y*_ control the center of the Gabor, *σ* is the standard deviation of the Gaussian envelope, and *θ, λ* and *ψ* control the orientation, spatial frequency and phase of the grating, respectively. For all simulated cells, *µ*_*x*_ and *µ*_*y*_ were set to 0; *θ* and *psi* were randomly sampled from the range of [0, *π*] and [0,2*π*], respectively. We then selected *σ* and *λ* values that closely match neuronal properties in our dataset. *σ* values were selected from the range of [4.4°, 10.9°] visual angle, as inferred from MEI mask sizes from 1000 random neurons in 8 pilot datasets. For */*, we first searched for Gabor images with the highest predicted activation for real neurons using a range of [0.02° /cycle, 0.12° /cycle] (Niell and Stryker, 2008), and then randomly selected *λ* values from those corresponding to the optimal Gabor images. We then randomly combined these parameters to simulate the ground-truth Gabor stimuli for 60 simple and 60 complex cells. To ensure sufficient frequency representation within the Gaussian window, we constrained *λ* to be no more than twice the value of *σ*.

For each simulated cell, we collected idealized responses to 5000 random ImageNet images, using each response as the mean of a Poisson distribution from which we sampled a noisy response. This noisy input-response dataset was then used to train a predictive model with an architecture identical to that used for real neurons. Finally, we applied the same image optimization procedure described above to generate MEI and DEIs using the simple and complex cell predictive models. This procedure aims to simulate the noise inherent in biological systems and create predictive models for simulated cells that more closely resemble those of real neurons.

### Representational similarity

To quantify similarity of visual stimuli in a space that is relevant to mouse V1 population functionality, we first obtained a low-dimensional latent representation for each stimulus and then assessed the similarity between stimulus pairs using this latent representation. We used a model ensemble trained on a held-out dataset to predict population responses to a random set of MEIs from 14 different animals (500 per animal) after these MEIs were centered and standardized. We then performed Principal Component Analysis (PCA) to identify the top 53 principal components (PCs) capable of explaining 95% of the response variance across all neurons. For any given stimulus, we centered and standardized it, passed it through the designated model ensemble, and projected the resultant population response onto the 53-dimensional space to derive its latent neuronal representation. We then computed representational similarity of each pair of stimuli using cosine similarity in this latent space. To compute similarity between sets of stimuli (e.g. sets of DEIs generated from various conditions), we calculated the average pairwise representational similarity.

#### The CUB and CUB-grating datasets

To study the relationship between invariance and natural stimuli, we sampled over 1 million crops from the Caltech-UCSD Birds-200-2011 (CUB) image dataset (Wah et al., 2011). This dataset contains 11,788 natural images across 200 bird categories, each featuring a single bird in its natural habitat. We resized original images to 64×64 pixels and sampled them using a 36×36 pixel window with a stride of 2. Each image also includes a manual semantic segmentation label identifying the bird region as a probability map, which we binarized using a threshold of 0.5. To test whether object boundaries defined by spatial frequency differences strongly activate V1 neurons, we created a modified dataset, “CUB-grating”, by replacing object and background content with grating patterns. We generated four equal-sized image types (2 million each):

1. Homogeneous grating pattern without using segmentation labels (“single grating”).
2. Same spatial frequency but varying orientations
3. Same orientation but varying spatial frequencies
4. Both varying frequencies and orientations

To determine the frequency range for high and low frequency patterns, we sampled 1000 random neurons across 8 pilot mice and fitted optimal Gabor-filter stimuli for each neuron using their corresponding predictive model. We defined high frequency as 5.83°/cycle and 15.55°/cycle (5th to 50th percentile) and low frequency as 15.55°/cycle to 58.3°/cycle (50th to 95th percentile). We independently uniformly sampled frequency, orientations, and phases for both the object and background gratings, then normalized them to have identical mean and RMS contrast. We masked the grating images with their corresponding object and background masks from the binarized segmentation label. To minimize edge artifacts, we applied a Gaussian filter (*σ* = 1.5) to blur object-background boundaries.

#### Analyses on highly activating crops in the CUB and CUB– grating datasets

To assess the alignment of the spatial structure between neurons’ bipartite RF and the object-background division in the CUB dataset, we screened over 1 million CUB image crops *in silico* for each target neuron to identify the 100 most highly activating ones. Each crop was masked from a full-field image with the target neuron’s MEI mask and standardized to match the MEI’s mean and RMS contrast within the mask. We classified a crop as containing object boundary if it comprised at least 20% of both object and background within the target neuron’s receptive field (RF). We also reproduced our findings with 10% and 30% thresholds. Crops without object boundaries were excluded from downstream analyses. To obtain a bipartite mask for each neuron, we binarized its MEI mask 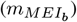 and variable subfield mask 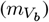 by thresholding at 0.3, assigning a value of 1 for each pixel within 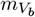 and −1 for each remaining pixel within 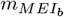. Similarly, using the manual segmentation label for each image crop, we assigned a value of 1 if the pixel is within the object and −1 if the pixel belongs to the background. We quantified the alignment between a crop’s segmentation label and the neuron’s bipartite mask using a matching score defined as 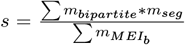 where a score of 1 indicates perfect alignment of the variable subfield with the object and fixed subfield with the background, and 0 indicates the reverse. The same procedure was applied on 100 random images to serve as a baseline to account for inherent bias of the dataset. This protocol was also used to evaluate matching in the CUB-grating dataset.

#### Spatial frequency tuning analysis in bipartite subfields

To systematically study how neuronal responses vary with spatial frequency in each subfield, we performed additional analyses using natural images. For each target neuron, we applied the fixed subfield mask to the CUB natural image dataset to extract 10,000 random crops and computed their median spatial frequency. These crops were then combined with the original variable subfield (masked directly from the MEI), and the resulting images were fed into our predictive model to obtain predicted responses. For each neuron, we calculated the Pearson correlation coefficient between the median frequency of the crops and the predicted responses. We repeated this process for the variable subfield as well. Finally, we corrected for multiple comparisons using the Benjamini–Hochberg procedure and reported only those neurons with a significant correlation (*p<* 0.05) after correction.

### Functional connectomics analyses on the MICrONS dataset

#### Replication of DEIs in MICrONS and closed-loop validation

Recently, a large-scale functional connectomics dataset of mouse visual cortex (“MICrONS”), including responses of >75k neurons to full-field natural movies and the reconstructed sub-cellular connectivity of the same cells from electron microscopy data (MICrONS Consortium et al., 2021). A dynamic digital twin model (Wang et al., 2023) of mouse visual cortex exhibits not only a high predictive performance for natural movies, but also accurate out-of-domain performance on other stimulus classes such as drifting Gabor filters, directional pink noise, and random dot kinematograms. Here, we took advantage of the model’s ability to generalize to other visual stimulus domains and extracted specific functional properties from this digital twin model in order to relate them to the neurons’ connectivity and anatomical properties. Specifically, we first trained a dynamic digital twin of the CvT-Lstm architecture (Wang et al., 2023) in the MICrONS neuronal population using all movie clips from the MICrONS stimulus set. We then presented a movie of 5100 unique natural images as described in *Presentation of natural stimuli* (except that every image was shown once since the model prediction is deterministic) to the dynamic digital twin and used these *in silico* responses to train a static model (“dynamic-to-static” model). MEIs, DEIs, and the bipartite parameterization were subsequently extracted from the “dynamic-to-static” model.

To validate the images generated from this “dynamic-to-static” model, we recorded the visual activity of the same neuronal population to static natural images as well as to the identical natural movies that were used in the MICrONS dataset in 3 new mice. Neurons across scans were matched anatomically as described in *Cell matching across days*. Based on neuronal responses we trained two versions of static models: one directly on static image responses as described in previous sections, and one “dynamic-to-static” model. We then presented MEIs, DEIs, and partial texture DEIs extracted from both static models back to the mice in closed-loop experiments. Since the static and dynamic stimuli were presented in two separate scans on day 1, only neurons that had unique one-to-one matching between the two scans (54 *±* 3%) and had matching distance smaller than 5*µm* (76 *±* 5%) were considered for image synthesis. We further excluded neurons with bottom 1 percentile of *CC*_*max*_ and *CC*_*abs*_ and then randomly selected neurons from the remaining population for closed-loop validation. For scans with synthesized images, only neurons that have good matching (criteria as described in) to both of the day 1 static and dynamics scans were included for data analysis. On average, 33 *±* 2% of closed-loop neurons per scan were kept for data analysis.

#### Neuron selection for functional connectomics analysis

We focused our analysis exclusively on V1 Layer 2/3 excitatory neurons, using area membership labels per neuron provided by the MICrONS release (MICrONS Consortium et al., 2021; Ding et al., 2023). Neurons with reliable visual responses (*CC*_*max*_ *>* 0.4) that are well predicted by the digital twin model (*CC*_*abs*_ *>* 0.2) were included in the downstream analysis, following the methodology described in Schoppe et al. (2016) and Ding et al. (2023). These criteria resulted in 19 presynaptic neurons and 706 connected pairs for downstream analysis.

To control for neuronal connectivity at a finer synaptic level, we followed the procedure outlined in Ding et al. (2023) to identify Axonal-Dendritic Proximity (ADP) controls. These neurons had a dendritic skeleton passing within 5 µm of the presynaptic neuron’s axonal skeleton (3D Euclidean distance) but were not observed to form a synapse with the presynaptic neuron. This process produced 2,486 ADP neurons and 18,162 pairs of ADP controls.

#### Functional analysis on the MICrONS dataset

To elucidate functional differences between connected pairs and Axonal-Dendritic Proximity (ADP) controls, we aggregated data across all presynaptic neurons. However, naive aggregation is problematic due to varying functional and connectivity metrics of different presynaptic neurons.

To address this, we performed the following corrections:

##### Correction on functional metric

We implemented a two-step standardization process for each pairwise metric, i.e. MEI and DEI pairwise similarity, and diversity index difference. First, we adjusted the pairwise value by subtracting the presynaptic neuron’s mean value, calculated as the average across all its connected pairs and ADP control pairs. We then added back a regional baseline level, computed as the mean value across all connected pairs and ADP controls within V1 Layer 2/3. This correction was applied to all pairwise metrics for both connected pairs and ADP controls.

##### Correction on connectivity metric

When aggregating connected pairs, we weighted each pair by the number of synapses observed between them and then adjusted for presynaptic neuron synapse conversion rates. We calculated the synapse conversion rate for each presynaptic neuron as the ratio between the total number of synapses formed from its axon and the total co-traveling distance with dendrites from its postsynaptic targets and ADP controls within V1 Layer 2/3. The expected number of synapses between a pair was then calculated as the product of this rate and the co-traveling distance. We adjusted the observed number of synapses by this expected value and added back a regional expected number of synapses based on the pair’s co-traveling distance and the regional synapse conversion rate.

We then performed weighted bootstrapping on connected and ADP pairs independently, using the adjusted number of synapses as weight for connected pairs and co-traveling distance for ADP controls. To quantify synapse conversion rate as a function of functional similarity, we adapted the procedure from Ding et al. (2023). We binned all neuron pairs (both connected and ADP control) according to their pairwise value. For each bin, synapse conversion rate was defined as the ratio of the number of observed synapses to the total co-traveling distance between presynaptic neurons’ axon arbors and their targets’ dendritic skeletons within the bin.

We included only bins containing more than 10 connected neuron pairs and representing at least than 2.5% of all connected neuron pairs. To estimate the standard deviation of the synapse conversion rate, we resampled the connected and ADP pairs with replacement, binned the resampled distributions, and calculated the standard deviation within each bin.

## Statistics

All statistical tests were reported directly in figure captions with corresponding sample sizes, test statistics, and p-values. Permutation tests and bootstrapping procedures were conducted using 10,000 permutations or resamplings with replacement. P-values for permutation tests and bootstrapping less than 10^−4^ were reported as *P <* 10^−4^; otherwise, exact p-values were provided. The linear coefficient was computed as the average of values obtained from 1,000 independent robust regressions using the RANSAC algorithm (Fischler and Bolles, 1981). For Welch’s *t*-tests and one-sample *t*-tests, normality was assumed but not explicitly tested. In Wilcoxon signed-rank tests, p-values less than 10^−9^ were reported as *P <* 10^−9^; otherwise, exact p-values were provided. For multiple comparisons, we applied the Benjamini-Hochberg (BH) correction and reported both the fraction of significant comparison before and after correction, along with corrected p-values.

## Software

Experiments and analysis are carried out with custom built data pipelines. The data pipeline is developed in Matlab and Python with the following tools: Psychtoolbox, ScanImage, DeepLabCut, CAIMAN, and Labview were used for data collection. DataJoint, MySQL, and CAVE were used for storing and managing data. Numpy, pandas, SciPy, statsmodels, scikit-learn, and PyTorch were used for model training and statistical analysis. Matplotlib, seaborn were used for graphical visualization. Jupyter, Docker, and Kubernetes were used for code development and deployment.

## Data and code availability

All data and the analysis code will be made publicly available in an online repository latest upon journal publication. Please contact us if you would like to get access before that time.

## References

E. H. Adelson and J. R. Bergen. Spatiotemporal energy models for the perception of motion. Josa a, 2(2):284–299, 1985.

J.-M. Alonso and L. M. Martinez. Functional connectivity between simple cells and complex cells in cat striate cortex. Nature neuroscience, 1(5):395–403, 1998.

Anselmi, L. Rosasco, C. Tan, and T. Poggio. Deep convolutional networks are hierarchical kernel machines. arXiv preprint arXiv:1508.01084, 2015.

Baroni, M. Bashiri, K. F. Willeke, J. Antolík, and F. H. Sinz. Learning invariance manifolds of visual sensory neurons. In NeurIPS Workshop on Symmetry and Geometry in Neural Representations, pages 301–326. PMLR, 2023.

P. Bashivan, K. Kar, and J. J. DiCarlo. Neural population control via deep image synthesis. Science, 364(6439):eaav9436, 2019.

I. Biederman. Recognition-by-components: a theory of human image understanding. Psychological review, 94(2):115, 1987.

D. Birman, K. J. Yang, S. J. West, B. Karsh, Y. Browning, I. B. Laboratory, J. H. Siegle, and N. A. Steinmetz. Pinpoint: trajectory planning for multi-probe electrophysiology and injections in an interactive web-based 3d environment. bioRxiv, 2023.

K. L. Briggman and D. D. Bock. Volume electron microscopy for neuronal circuit reconstruction. Current opinion in neurobiology, 22(1):154–161, 2012.

J. J. Bussgang. Crosscorrelation functions of amplitude-distorted gaussian signals. Technical Report 216, 1952.

S. A. Cadena, M. A. Weis, L. A. Gatys, M. Bethge, and A. S. Ecker. Diverse feature visualizations reveal invariances in early layers of deep neural networks. In Proceedings of the European Conference on Computer Vision (ECCV), pages 217–232, 2018.

S. A. Cadena, G. H. Denfield, E. Y. Walker, L. A. Gatys, A. S. Tolias, M. Bethge, and A. S. Ecker. Deep convolutional models improve predictions of macaque v1 responses to natural images. PLoS computational biology, 15(4):e1006897, 2019.

C. Cadieu, M. Kouh, A. Pasupathy, C. E. Connor, M. Riesenhuber, and T. Poggio. A model of v4 shape selectivity and invariance. Journal of neurophysiology, 98(3):1733–1750, 2007.

Ding, Tran et al. | Bipartite invariance in mouse primary visual cortex B. Celii, S. Papadopoulos, Z. Ding, P. G. Fahey, E. Wang, C. Papadopoulos, A. Kunin, S. Patel, J. Alexander Bae, A. L. Bodor, D. Brittain, J. Buchanan, D. J. Bumbarger, M. A. Castro, E. Cobos, S. Dorkenwald, L. Elabbady, A. Halageri, Z. Jia, C. Jordan, D. Kapner, N. Kemnitz, S. Kinn, K. Lee, K. Li, R. Lu, T. Macrina, G. Mahalingam, E. Mitchell, S. S. Mondal, S. Mu, B. Nehoran, S. Popovych, C. M. Schneider-Mizell, W. Silversmith, M. Takeno, R. Torres, N. L. Turner, W. Wong, J. Wu, S.-C. Yu, W. Yin, D. Xenes, L. M. Kitchell, P. K. Rivlin, V. A. Rose, C. A. Bishop, B. Wester, E. Froudarakis, E. Y. Walker, F. H. Sinz, H. Sebastian Seung, F. Collman, N. M. da Costa, R. Clay Reid, X. Pitkow, A. S. Tolias, and J. Reimer. NEURD: A mesh decomposition framework for automated proofreading and morphological analysis of neuronal EM reconstructions. bioRxiv, page 2023.03.14.532674, Mar. 2023.

G. Crutcher. Lateral connections improve generalizability of learning in a simple neural network. Neural Computation, 36(4):705–717, 2024.

J. Dapello, T. Marques, M. Schrimpf, F. Geiger, D. Cox, and J. J. DiCarlo. Simulating a primary visual cortex at the front of cnns improves robustness to image perturbations. Advances in Neural Information Processing Systems, 33:13073–13087, 2020.

J. J. DiCarlo and D. D. Cox. Untangling invariant object recognition. Trends in cognitive sciences, 11(8):333–341, 2007.

J. J. DiCarlo, D. Zoccolan, and N. C. Rust. How does the brain solve visual object recognition? Neuron, 73(3):415–434, 2012.

Z. Ding, P. G. Fahey, S. Papadopoulos, E. Y. Wang, B. Celii, C. Papadopoulos, A. B. Kunin, A. Chang, J. Fu, Z. Ding, et al. Functional connectomics reveals general wiring rule in mouse visual cortex. bioRxiv, 2023.

Y. El-Shamayleh and A. Pasupathy. Contour curvature as an invariant code for objects in visual area v4. Journal of Neuroscience, 36(20):5532–5543, 2016.

M. A. Fischler and R. C. Bolles. Random sample consensus: a paradigm for model fitting with applications to image analysis and automated cartography. Communications of the ACM, 24 (6):381–395, 1981.

K. Franke, K. F. Willeke, K. Ponder, M. Galdamez, N. Zhou, T. Muhammad, S. Patel, E. Froudarakis, J. Reimer, F. H. Sinz, et al. State-dependent pupil dilation rapidly shifts visual feature selectivity. Nature, 610(7930):128–134, 2022.

E. Froudarakis, P. Berens, A. S. Ecker, R. J. Cotton, F. H. Sinz, D. Yatsenko, P. Saggau, M. Bethge, and A. S. Tolias. Population code in mouse v1 facilitates readout of natural scenes through increased sparseness. Nature neuroscience, 17(6):851–857, 2014.

J. Fu, S. Shrinivasan, K. Ponder, T. Muhammad, Z. Ding, E. Wang, Z. Ding, D. Tran, P. G. Fahey, S. Papadopoulos, S. Patel, J. Reimer, A. S. Ecker, X. Pitkow, R. M. Haefner, F. H. Sinz, K. Franke, and A. S. Tolias. Pattern completion and disruption characterize contextual modulation in mouse visual cortex. bioRxiv, 2024.

K. Fukushima. Neocognitron: A self-organizing neural network model for a mechanism of pattern recognition unaffected by shift in position. Biological cybernetics, 36(4):193–202, 1980.

C. D. Gilbert and T. N. Wiesel. The influence of contextual stimuli on the orientation selectivity of cells in primary visual cortex of the cat. Vision research, 30(11):1689–1701, 1990.

A. Giovannucci, J. Friedrich, P. Gunn, J. Kalfon, B. L. Brown, S. A. Koay, J. Taxidis, F. Najafi, J. L. Gauthier, P. Zhou, B. S. Khakh, D. W. Tank, D. B. Chklovskii, and E. A. Pnevmatikakis. Caiman: An open source tool for scalable calcium imaging data analysis. eLife, 8:e38173, 2019.

C. G. Gross, C. d. Rocha-Miranda, and D. Bender. Visual properties of neurons in inferotemporal cortex of the macaque. Journal of neurophysiology, 35(1):96–111, 1972.

D. J. Heeger. Half-squaring in responses of cat striate cells. Visual neuroscience, 9(5):427–443, 1992.

I. Higgins, S. Racanière, and D. Rezende. Symmetry-based representations for artificial and biological general intelligence. Frontiers in Computational Neuroscience, page 28, 2022.

D. N. Hill, S. B. Mehta, and D. Kleinfeld. Quality metrics to accompany spike sorting of extracellular signals. Journal of Neuroscience, 31(24):8699–8705, 2011.

D. H. Hubel and T. N. Wiesel. Receptive fields, binocular interaction and functional architecture in the cat’s visual cortex. The Journal of physiology, 160(1):106, 1962.

J. P. Jones and L. A. Palmer. An evaluation of the two-dimensional gabor filter model of simple receptive fields in cat striate cortex. Journal of neurophysiology, 58(6):1233–1258, 1987a.

J. P. Jones and L. A. Palmer. The two-dimensional spatial structure of simple receptive fields in cat striate cortex. Journal of neurophysiology, 58(6):1187–1211, 1987b.

J. J. Jun, N. A. Steinmetz, J. H. Siegle, D. J. Denman, M. Bauza, B. Barbarits, A. K. Lee, C. A. Anastassiou, A. Andrei, Ç. Aydın, et al. Fully integrated silicon probes for high-density recording of neural activity. Nature, 551(7679):232–236, 2017.

M. K. Kapadia, M. Ito, C. D. Gilbert, and G. Westheimer. Improvement in visual sensitivity by changes in local context: parallel studies in human observers and in v1 of alert monkeys. Neuron, 15(4):843–856, 1995.

Y. Karklin and M. S. Lewicki. Emergence of complex cell properties by learning to generalize in natural scenes. Nature, 457(7225):83–86, 2009.

C. Keck and J. Lücke. Learning of lateral connections for representational invariant recognition. In Artificial Neural Networks–ICANN 2010: 20th International Conference, Thessaloniki, Greece, September 15-18, 2010, Proceedings, Part III 20, pages 21–30. Springer, 2010.

L. Kirchberger, S. Mukherjee, U. H. Schnabel, E. H. van Beest, A. Barsegyan, C. N. Levelt, J. A. Heimel, J. A. Lorteije, C. van der Togt, M. W. Self, et al. The essential role of recurrent processing for figure-ground perception in mice. Science advances, 7(27):eabe1833, 2021.

V. Klymenko and N. Weisstein. Spatial frequency differences can determine figure-ground organization. Journal of Experimental Psychology:Human Perception and Performance, 12(3): 324–330, 1986.

V. Klymenko, N. Weisstein, R. Topolski, and C.-H. Hsieh. Spatial and temporal frequency in figure-ground organization. Perception and Psychophysics, 45(4):395–403, 1989.

H. Ko, S. B. Hofer, B. Pichler, K. A. Buchanan, P. J. Sjöström, and T. D. Mrsic-Flogel. Functional specificity of local synaptic connections in neocortical networks. Nature, 473(7345):87–91, 2011.

W.-C. A. Lee, V. Bonin, M. Reed, B. J. Graham, G. Hood, K. Glattfelder, and R. C. Reid. Anatomy and function of an excitatory network in the visual cortex. Nature, 532(7599):370–374, Apr. 2016.

J. W. Lichtman and W. Denk. The big and the small: challenges of imaging the brain’s circuits. Science, 334(6056):618–623, 2011.

K.-K. Lurz, M. Bashiri, K. Willeke, A. K. Jagadish, E. Wang, E. Y. Walker, S. A. Cadena, T. Muhammad, E. Cobos, A. S. Tolias, A. S. Ecker, and F. H. Sinz. Generalization in data-driven models of primary visual cortex. In Proceedings of the International Conference for Learning Representations (ICLR), page 2020.10.05.326256, Oct. 2020.

M. Maruyama, F. Girosi, and T. Poggio. A connection between grbf and mlp. 1992.

A. Mathis, P. Mamidanna, K. M. Cury, T. Abe, V. N. Murthy, M. W. Mathis, and M. Bethge. DeepLabCut: markerless pose estimation of user-defined body parts with deep learning. Nature Neuroscience, 21(9):1281–1289, 2018. ISSN 15461726. doi: 10.1038/s41593-018-0209-y. URL http://dx.doi.org/10.1038/s41593-018-0209-y.

MICrONS Consortium, J. Alexander Bae, M. Baptiste, A. L. Bodor, D. Brittain, J. Buchanan, D. J. Bumbarger, M. A. Castro, B. Celii, E. Cobos, F. Collman, N. M. da Costa, S. Dorkenwald, L. Elabbady, P. G. Fahey, T. Fliss, E. Froudarakis, J. Gager, C. Gamlin, A. Halageri, J. Hebditch, Z. Jia, C. Jordan, D. Kapner, N. Kemnitz, S. Kinn, S. Koolman, K. Kuehner, K. Lee, K. Li, R. Lu, T. Macrina, G. Mahalingam, S. McReynolds, E. Miranda, E. Mitchell, S. S. Mondal, M. Moore, S. Mu, T. Muhammad, B. Nehoran, O. Ogedengbe, C. Papadopoulos, S. Papadopoulos, S. Patel, X. Pitkow, S. Popovych, A. Ramos, R. Clay Reid, J. Reimer, C. M. Schneider-Mizell, H. Sebastian Seung, B. Silverman, W. Silversmith, A. Sterling, F. H. Sinz, C. L. Smith, S. Suckow, M. Takeno, Z. H. Tan, A. S. Tolias, R. Torres, N. L. Turner, E. Y. Walker, T. Wang, G. Williams, S. Williams, K. Willie, R. Willie, W. Wong, J. Wu, C. Xu, R. Yang, D. Yatsenko, F. Ye, W. Yin, and S.-C. Yu. Functional connectomics spanning multiple areas of mouse visual cortex. Aug. 2021.

C. M. Niell and M. P. Stryker. Highly selective receptive fields in mouse visual cortex. Journal of Neuroscience, 28(30):7520–7536, 2008.

C. Olah, A. Mordvintsev, and L. Schubert. Feature visualization. Distill, 2017. doi: 10.23915/distill.00007. https://distill.pub/2017/feature-visualization.

C. Olah, N. Cammarata, L. Schubert, G. Goh, M. Petrov, and S. Carter. Zoom in: An introduction to circuits. Distill, 5(3):e00024–001, 2020.

M. Pachitariu, S. Sridhar, and C. Stringer. Solving the spike sorting problem with kilosort. BioRxiv, pages 2023–01, 2023.

E. A. Pnevmatikakis, D. Soudry, Y. Gao, T. A. Machado, J. Merel, D. Pfau, T. Reardon, Y. Mu, C. Lacefield, W. Yang, et al. Simultaneous denoising, deconvolution, and demixing of calcium imaging data. Neuron, 89(2):285–299, 2016.

T. Poggio and E. Bizzi. Generalization in vision and motor control. Nature, 431(7010):768–774, 2004.

T. Poggio and F. Girosi. Networks for approximation and learning. Proceedings of the IEEE, 78 (9):1481–1497, 1990.

C. R. Ponce, W. Xiao, P. F. Schade, T. S. Hartmann, G. Kreiman, and M. S. Livingstone. Evolving images for visual neurons using a deep generative network reveals coding principles and neuronal preferences. Cell, 177(4):999–1009, 2019.

R. Q. Quiroga, L. Reddy, G. Kreiman, C. Koch, and I. Fried. Invariant visual representation by single neurons in the human brain. Nature, 435(7045):1102–1107, 2005.

J. Reimer, E. Froudarakis, C. R. Cadwell, D. Yatsenko, G. H. Denfield, and A. S. Tolias. Pupil fluctuations track fast switching of cortical states during quiet wakefulness. neuron, 84(2): 355–362, 2014.

M. Riesenhuber and T. Poggio. Hierarchical models of object recognition in cortex. Nature neuroscience, 2(11):1019–1025, 1999.

L. F. Rossi, K. D. Harris, and M. Carandini. Spatial connectivity matches direction selectivity in visual cortex. Nature, 588(7839):648–652, Dec. 2020.

O. Russakovsky, J. Deng, H. Su, J. Krause, S. Satheesh, S. Ma, Z. Huang, A. Karpathy, A. Khosla, M. Bernstein, A. C. Berg, and L. Fei-Fei. Imagenet large scale visual recognition challenge. International Journal of Computer Vision, 115:211–252, 2015.

N. C. Rust and J. J. DiCarlo. Selectivity and tolerance (“invariance”) both increase as visual information propagates from cortical area v4 to it. Journal of Neuroscience, 30(39):12978–12995, 2010.

U. H. Schnabel, C. Bossens, J. A. Lorteije, M. W. Self, H. Op de Beeck, and P. R. Roelfsema. Figure-ground perception in the awake mouse and neuronal activity elicited by figure-ground stimuli in primary visual cortex. Scientific reports, 8(1):17800, 2018.

O. Schoppe, N. S. Harper, B. D. Willmore, A. J. King, and J. W. Schnupp. Measuring the performance of neural models. Frontiers in computational neuroscience, 10:10, 2016.

M. Schrimpf, J. Kubilius, H. Hong, N. J. Majaj, R. Rajalingham, E. B. Issa, K. Kar, P. Bashivan, J. Prescott-Roy, F. Geiger, K. Schmidt, D. L. K. Yamins, and J. J. DiCarlo. Brain-score: Which artificial neural network for object recognition is most brain-like? bioRxiv preprint, 2018. URL https://www.biorxiv.org/content/10.1101/407007v2.

M. Schrimpf, J. Kubilius, M. J. Lee, N. A. R. Murty, R. Ajemian, and J. J. DiCarlo. Integrative benchmarking to advance neurally mechanistic models of human intelligence. Neuron, 2020. URL https://www.cell.com/neuron/fulltext/S0896-6273(20)30605-X.

L. Schubert, C. Voss, N. Cammarata, G. Goh, and C. Olah. High-low frequency detectors. Distill, 2021. doi: 10.23915/distill.00024.005. https://distill.pub/2020/circuits/frequency-edges.

T. Serre and M. Riesenhuber. Realistic modeling of simple and complex cell tuning in the hmax-model, and implications for invariant object recognition in cortex. 2004.

T. O. Sharpee, M. Kouh, and J. H. Reynolds. Trade-off between curvature tuning and position invariance in visual area v4. Proceedings of the National Academy of Sciences, 110(28): 11618–11623, 2013.

F. Sinz, A. S. Ecker, P. Fahey, E. Walker, E. Cobos, E. Froudarakis, D. Yatsenko, Z. Pitkow, J. Reimer, and A. Tolias. Stimulus domain transfer in recurrent models for large scale cortical population prediction on video. Advances in neural information processing systems, 31, 2018.

N. J. Sofroniew, D. Flickinger, J. King, and K. Svoboda. A large field of view two-photon mesoscope with subcellular resolution for in vivo imaging. elife, 5:e14472, 2016.

S. Tafazoli, H. Safaai, G. De Franceschi, F. B. Rosselli, W. Vanzella, M. Riggi, F. Buffolo, S. Panzeri, and D. Zoccolan. Emergence of transformation-tolerant representations of visual objects in rat lateral extrastriate cortex. Elife, 6:e22794, 2017.

K. Tanaka. Inferotemporal cortex and object vision. Annual review of neuroscience, 19(1):109–139, 1996.

D. Y. Tsao, W. A. Freiwald, R. B. Tootell, and M. S. Livingstone. A cortical region consisting entirely of face-selective cells. Science, 311(5761):670–674, 2006.

R. von der Heydt and E. Peterhans. Mechanisms of contour perception in monkey visual cortex. i. lines of pattern discontinuity. Journal of Neuroscience, 9(5):1731–1748, 1989.

C. Wah, S. Branson, P. Welinder, P. Perona, and S. Belongie. The caltech-ucsd birds-200-2011 dataset. 2011.

E. Y. Walker, F. H. Sinz, E. Cobos, T. Muhammad, E. Froudarakis, P. G. Fahey, A. S. Ecker, J. Reimer, X. Pitkow, and A. S. Tolias. Inception loops discover what excites neurons most using deep predictive models. Nature neuroscience, 22(12):2060–2065, 2019.

B. Wang and C. R. Ponce. Tuning landscapes of the ventral stream. Cell Reports, 41(6):111595, 2022.

E. Y. Wang, P. G. Fahey, K. Ponder, Z. Ding, A. Change, T. Muhammad, S. Patel, Z. Ding, D. T. Tran, J. Fu, et al. Towards a foundation model of the mouse visual cortex. bioRxiv, pages 2023–03, 2023.

A. Wertz, S. Trenholm, K. Yonehara, D. Hillier, Z. Raics, M. Leinweber, G. Szalay, A. Ghanem, G. Keller, B. Rózsa, K.-K. Conzelmann, and B. Roska. PRESYNAPTIC NETWORKS. single-cell-initiated monosynaptic tracing reveals layer-specific cortical network modules. Science, 349(6243):70–74, July 2015.

K. F. Willeke, P. G. Fahey, M. Bashiri, L. Pede, M. F. Burg, C. Blessing, S. A. Cadena, Z. Ding, K.-K. Lurz, K. Ponder, et al. The sensorium competition on predicting large-scale mouse primary visual cortex activity. arXiv preprint arXiv:2206.08666, 2022.

D. L. Yamins and J. J. DiCarlo. Using goal-driven deep learning models to understand sensory cortex. Nature neuroscience, 19(3):356–365, 2016.

D. L. Yamins, H. Hong, C. F. Cadieu, E. A. Solomon, D. Seibert, and J. J. DiCarlo. Performance-optimized hierarchical models predict neural responses in higher visual cortex. Proceedings of the national academy of sciences, 111(23):8619–8624, 2014.

M. D. Zeiler and R. Fergus. Visualizing and understanding convolutional networks. In Computer Vision–ECCV 2014: 13th European Conference, Zurich, Switzerland, September 6–12, 2014, Proceedings, Part I 13, pages 818–833. Springer, 2014.

C. A. Zhan and C. L. Baker Jr. Boundary cue invariance in cortical orientation maps. Cerebral Cortex, 16(6):896–906, 2006.

