## Supplementary figures for "Bipartite invariance in mouse primary visual cortex"

### Supplementary Information

Supplementary Fig. 1 - Upper bound of the correlation coefficient and test correlation coefficient of V1 neurons.
Supplementary Fig. 2 - Example stimuli presented in closed-loop experiments.
Supplementary Fig. 3 - DEIs capture invariance observed in natural images with DEI-like activation.
Supplementary Fig. 4 - MEI and DEIs activated neurons with high specificity in all mice.
Supplementary Fig. 5 - Individual DEIs activated target neurons strongly.
Supplementary Fig. 6 - Similarity between MEI and DEI *in vivo* responses was not inflated by trial-to-trial eye movement.
Supplementary Fig. 7 - Neuronal-space DEIs evoked strong and selective *in vivo* responses in target neurons while exhibiting population-decodable differences.
Supplementary Fig. 8 - DEIs generalized across different conditions.
Supplementary Fig. 9 - Bipartite invariance quantification.
Supplementary Fig. 10 - Excluding neurons with small variable subfields did not alter the similarity between DEI and partial-texture DEI *in vivo* responses.
Supplementary Fig. 11 - Both subfields of the partial-texture DEIs are necessary and specific for evoking high *in vivo* responses.
Supplementary Fig. 12 - DEI closed-loop verification for randomly selected neurons.
Supplementary Fig. 13 - Example MEI, DEIs, and partial-texture DEIs from electrophysiological recordings for units classified as “single” (left) and “multiple” (right) based on spike sorting.
Supplementary Fig. 14 - Quantification of diversity and bipartite invariance indices from electrophysiological data.
Supplementary Fig. 15 - DEIs cannot be well explained by shift invariance in both subfields.
Supplementary Fig. 16 - Spatial division is necessary for explaining DEIs.
Supplementary Fig. 17 - Bipartite receptive field cannot be explained by the center-surround structure.
Supplementary Fig. 18 - Bipartite receptive field cannot be explained by neither trial-to-trial eye movement nor spatial readout location variation.
Supplementary Fig. 19 - DEI bipartite masks aligned with object boundaries in highly activating natural crops.
Supplementary Fig. 20 - Alignment between bipartite mask and natural object boundaries was robust across different thresholds for classifying patches containing object boundaries.
Supplementary Fig. 21 - Dynamic static model *in vivo* validation.
Supplementary Fig. 22 - MEI activated neurons with high specificity in both static and dynamic-static models.
Supplementary Fig. 23 - DEIs activated neurons with high specificity in both static and dynamic-static models.
Supplementary Fig. 24 - Partial-texture DEIs activated neurons with high specificity in both static and dynamic-static models.
Supplementary Fig. 25 - Postsynaptic neurons and ADP controls had similar diversity indices.

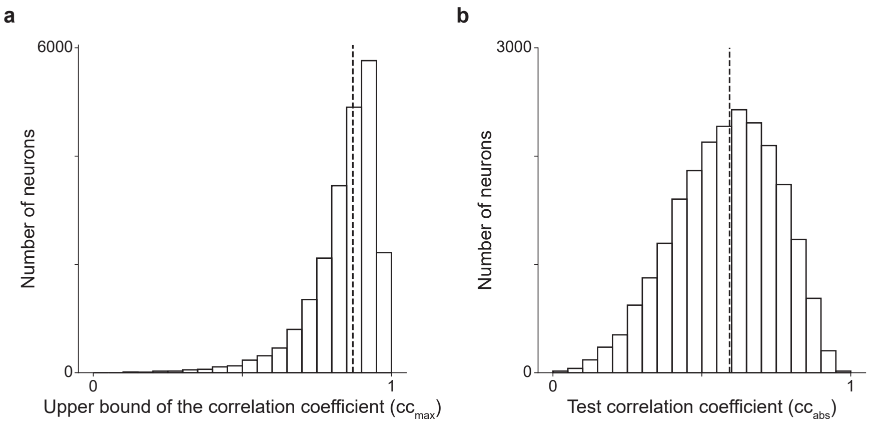


**Supplemental Fig. S1. Upper bound of the correlation coefficient and test correlation coefficient of V1 neurons. a**, Histogram of the upper bound of the correlation coefficient ($CC_{max}$) (Schoppe et al., 2016) for V1 neurons. The dashed line indicates median value at $0.87$. **b**, Histogram of the test correlation coefficient ($CC_{abs}$) (Schoppe et al., 2016) for V1 neurons. The dashed line indicates median value at $0.60$. Excessively noisy neurons ($CC_{max} < 0.1$) were excluded ($0.19$% of all neurons) and values outside of $0$ and $1$ are clipped ($0.01$% and $0.02$% for **a** and **b**, respectively) for visualization. Data were pooled over 33,714 neurons from 14 mice.

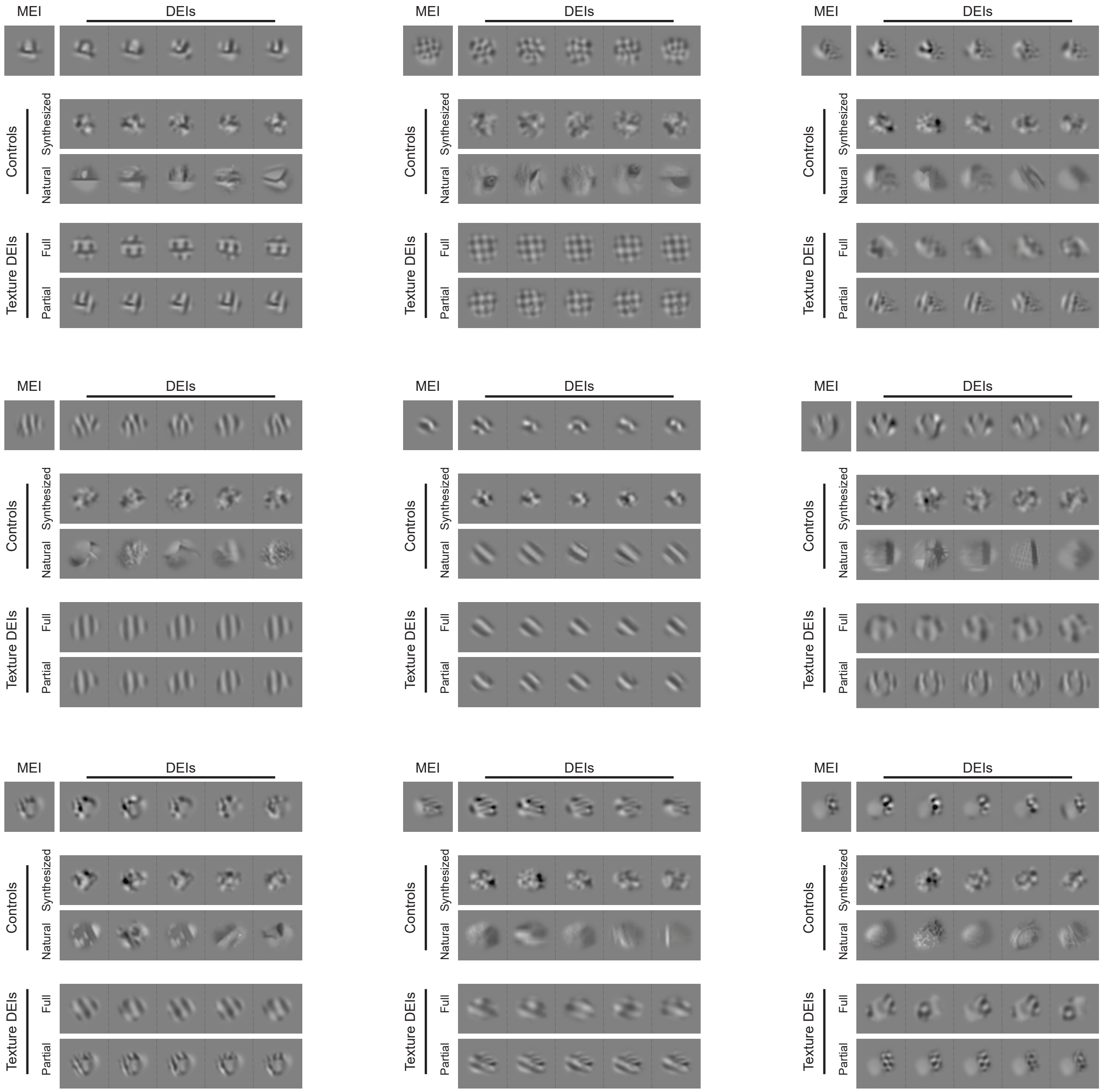


**Supplemental Fig. S2. Example stimuli presented in closed-loop experiments.** MEI, DEIs, DEIs controls including natural and synthesized controls, and parametric texture DEIs including partial-texture and full-texture DEIs that were presented back to the animals in closed-loop experiments for 9 example neurons.

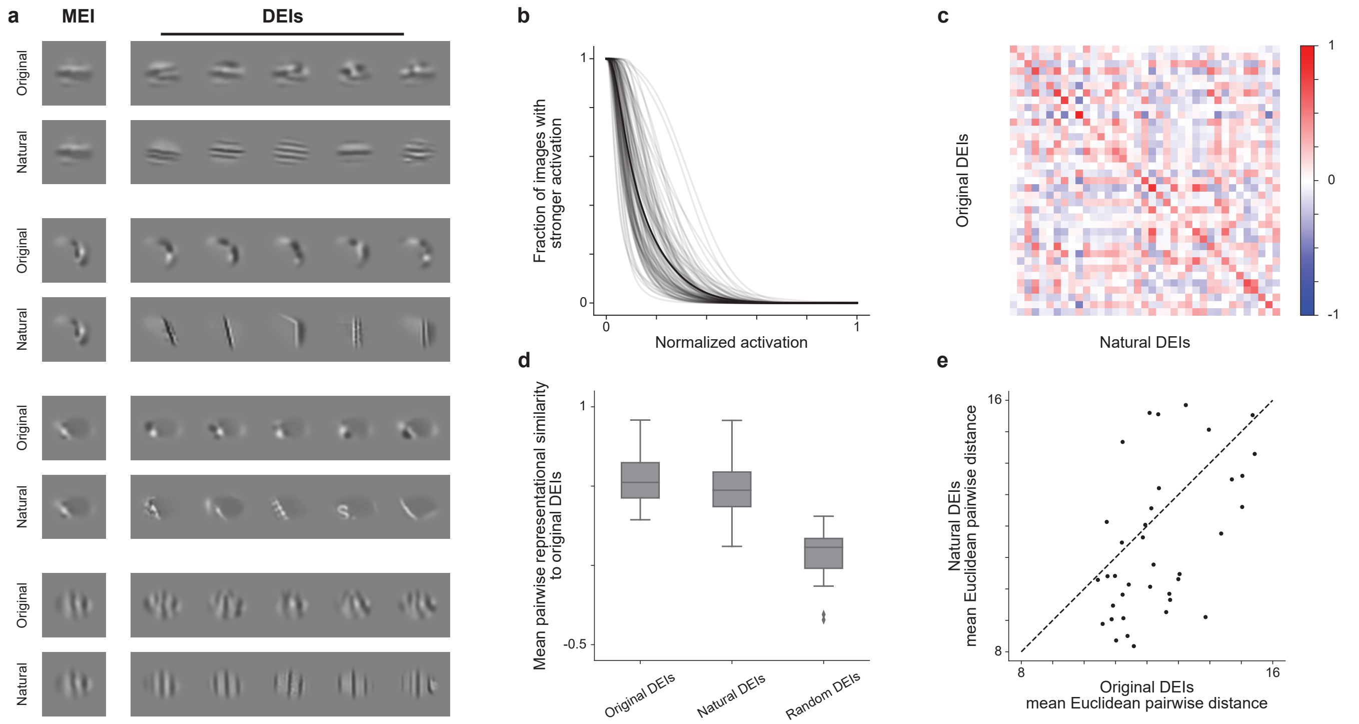


**Supplemental Fig. S3. DEIs capture invariance observed in natural images with DEI-like activation.** **a**, Examples of MEI, DEIs, and "natural DEIs" for 4 example neurons. For each neuron, we searched through 41 million ImageNet image patches *in silico* to identify natural crops that elicited activations equal to or greater than 85% of the MEI response (i.e. DEI-like activation). Among neurons with at least 20 natural DEIs (37 out of 100 neurons), we selected 20 natural DEIs—matching the number of synthesized DEIs per neuron—by greedily maximizing their minimum pairwise distance, mirroring the DEI synthesis procedure. These selected images are denoted as "natural DEIs". **b**, Neuron responses to masked natural images are sparse and smaller than those to MEIs and DEIs. The gray lines show the fraction out of 41 million masked images that elicit a given activation or higher for 100 model target neurons; black is the average. Responses from each cell are divided by the response to its MEI; on average, 1.2% of images produced activations above 50%, 0.02% above 75%, and 0.006% above 85% of the MEI activation. **b**, Natural DEIs maintained high specificity to their target neuron. Confusion matrices showed *In silico* representational similarity between original DEIs and highly activating natural crops. Each entry represents the mean pairwise cosine similarity between two sets of DEIs (see Methods for details). Representational similarity between original DEIs and highly activating natural crops for the same neurons (diagonal) was larger than cross-neuron similarity (off-diagonal) (two-sided permutation test, $P < 10^{-4}$ for all conditions after BH corrections). **c**, Natural DEIs closely resembled the original DEIs. The original DEIs were more similar to highly activating natural crops than random neurons' DEIs generated using the original method (two-sided Wilcoxon signed-rank test, $W = 0$, $P = 1.1 \times 10^{-7}$). **d**, Original DEIs have higher mean Euclidean pairwise distances than those of natural DEIs (two-sided Wilcoxon signed-rank test, $W = 179$, $P = 0.01$). Data were pooled over 100 neurons randomly sampled from 8 mice.

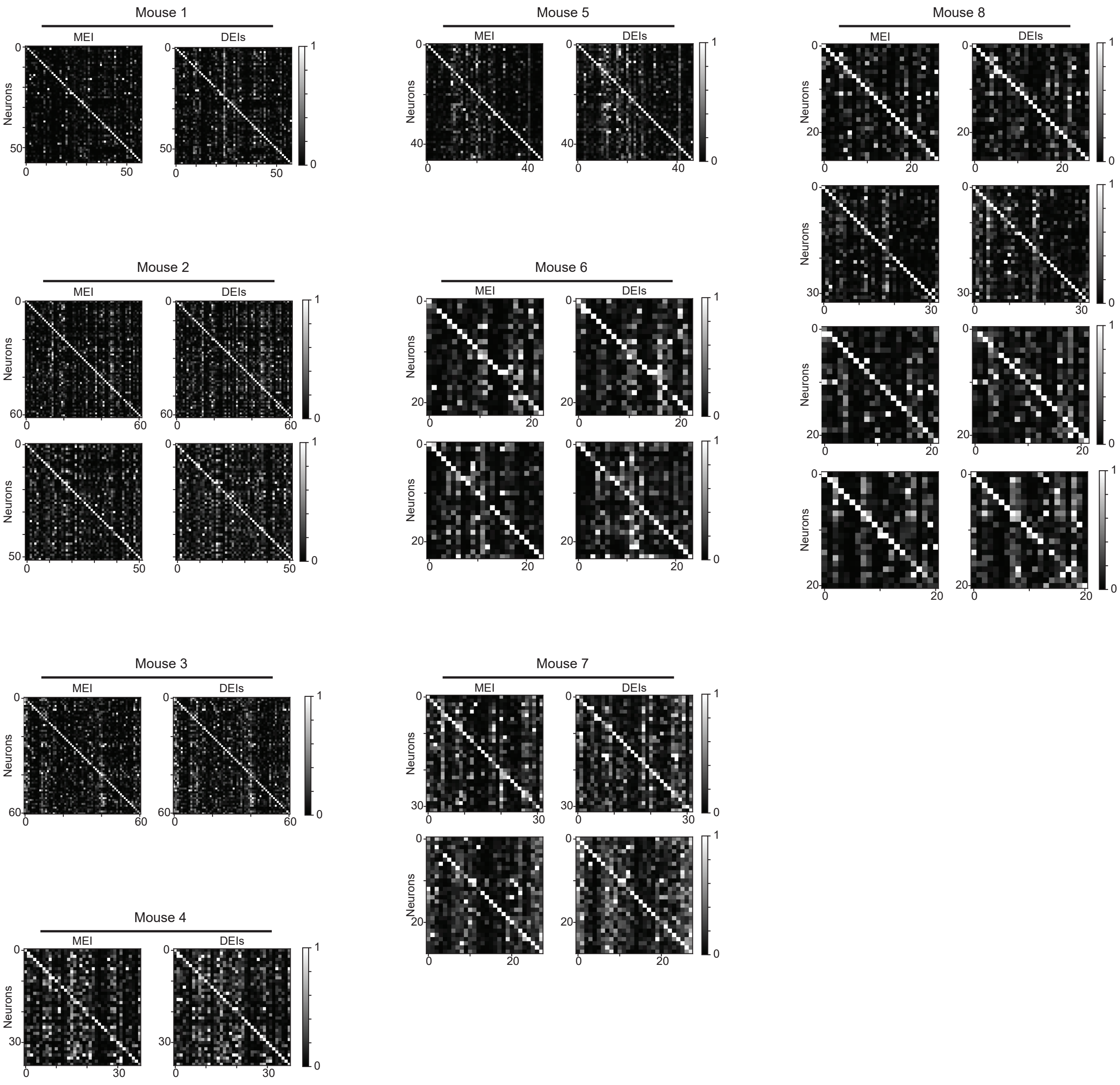


**Supplemental Fig. S4. MEI and DEIs activated neurons with high specificity in all mice**. The confusion matrices showed the responses of each neuron to the MEI (left) and DEIs (right) of all target neurons in individual scans where we presented the stimuli back to the mouse in closed-loop experiments. MEI responses were averaged across 20 repeats of the same image while DEIs responses were averaged across 20 different images with single repeat. The responses of each neuron were normalized, and each row was scaled so the maximum response across all images equals 1. Responses of neurons to their own MEI and DEIs (along the diagonal) were larger than to other MEIs and DEIs, respectively (two-sided permutation test, $P < 10^{-9}$ for both cases across all mice after BH correction).

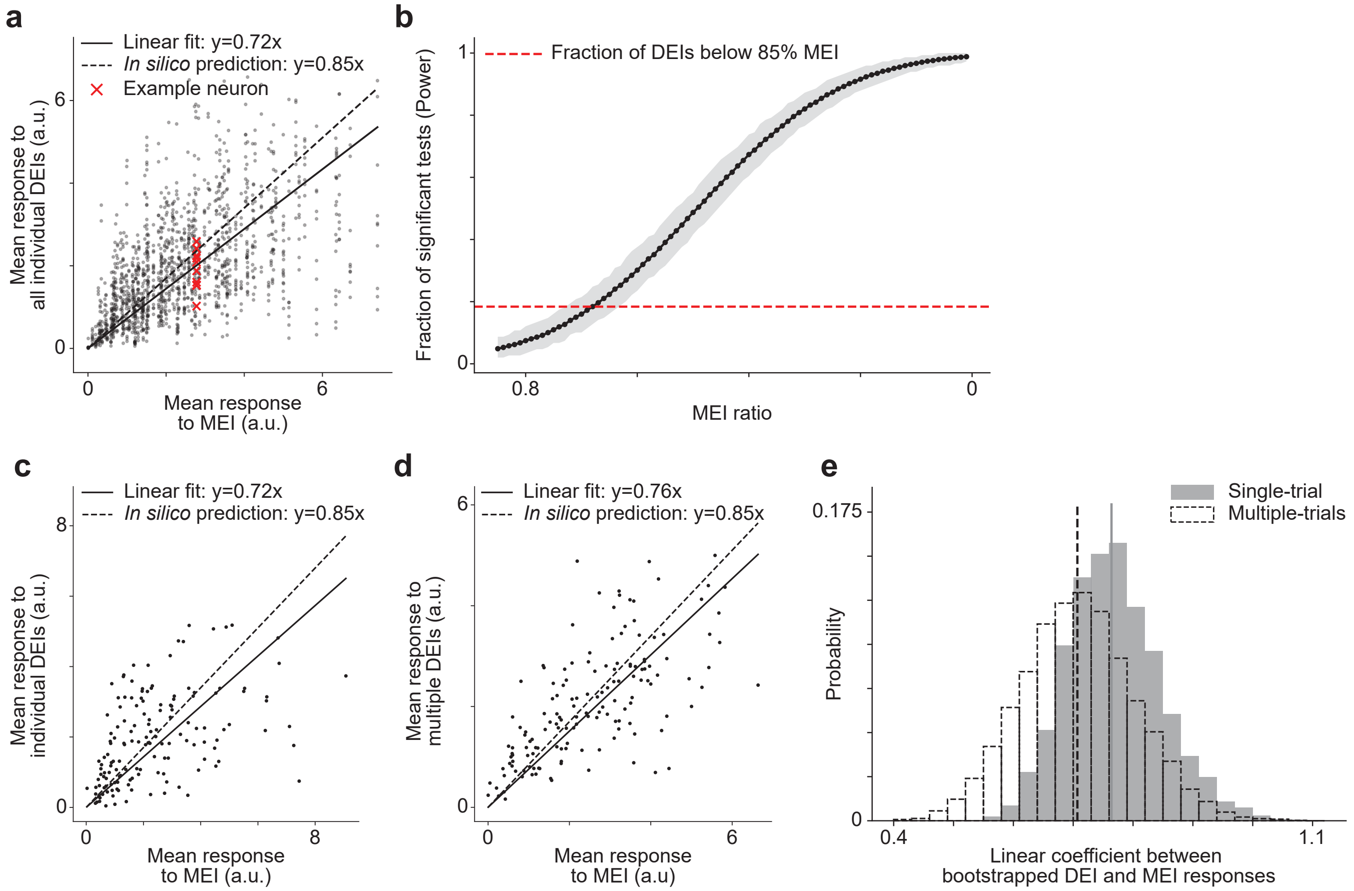


**Supplemental Fig. S5. Individual DEI activated their target neuron strongly.** **a**, For each neuron, we randomly selected 10 DEIs (red) from the set of 20 DEIs. Individual DEIs stimulated *in vivo* closely to the level predicted *in silico* with respect to MEI ($72 \pm 4\%$ versus $85\%$) (two-sided Wilcoxon signed-rank test, $W = 519434,\ P = 0.03$), with only 274 out of all 1490 individual DEIs ($18.4\%$) evoked responses lower than $85\%$ of the corresponding MEI ($4.8\%$ after BH correction; $P < 0.05$, one-sided Welch's $t$-test with $32.5$ average d.f.). **b**, Percentage of correctly rejected null hypotheses of statistical tests as a function of reduced response from 85% threshold. For each neuron, we randomly sampled two sets of 20 MEI trials with replacement: one set was scaled to 85% of the original MEI response (reference), and the other set was scaled to a smaller percentage to simulate different level of activation reduction. We then applied a one-tailed Welch's t-test ($sigma = 0.05$) to compare the two sets. The power is defined as the fraction of neurons where we detect a significant difference. We repeat this process 10,000 times and report the 95% CI in shaded area. **c,d**, To quantify the *in vivo* response relationship between MEI and DEIs, we bootstrapped by either averaging across 20 randomly selected trials from a single DEI (**b**) or 20 randomly selected trials from 10 different DEIs (**c**) (see Methods for details). **e**, The linear coefficients estimated using individual DEI (**c**, median=$0.71$) were similar to those estimated using multiple DEIs (**d**, median=$0.76$) ($P = 0.59$, two-sided bootstrapped mean difference against $0$). Data were pooled over 149 neurons from 2 mice.

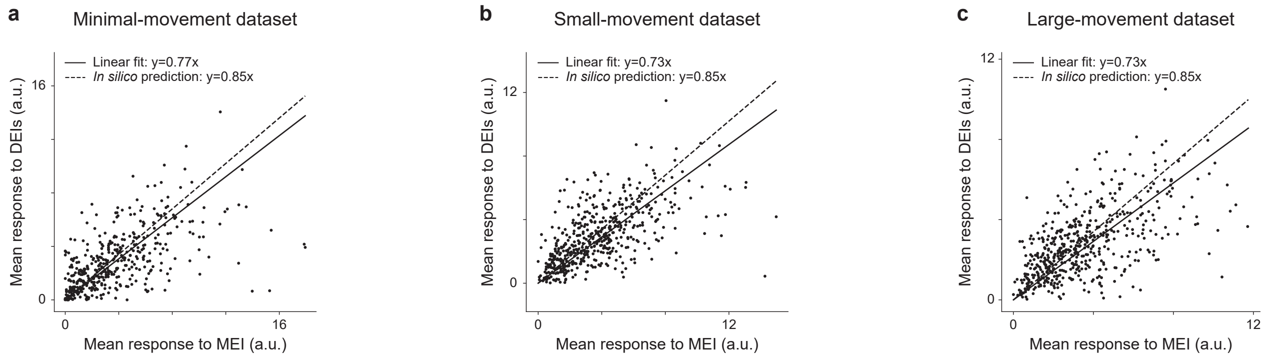


**Supplemental Fig. S6. Similarity between MEI and DEI *in vivo* responses was not inflated by trial-to-trial eye movement. a–c**, We created three sub-datasets from MEI and DEI trials using inclusion criteria based on different thresholds of eye movement size: a minimal-movement dataset including only trials with minor deviation from the average pupil position (approximately $19.3 \pm 12.3\%$ of all trials), a small-movement dataset including trials with pupil movements in the bottom $50^{\text{th}}$ percentile, and a large-movement dataset including trials with pupil movements in the top $50^{\text{th}}$ percentile (see Methods for details). DEIs strongly activated their target neurons comparable to the MEIs irrespective of the sub-datasets used for model fitting and image synthesis ($0.77 \pm 0.05$ for the minimal-movement dataset, $0.73 \pm 0.04$ for the small-movement dataset, $0.73 \pm 0.03$ for the large-movement dataset). Data were pooled over 500 neurons from 8 mice.

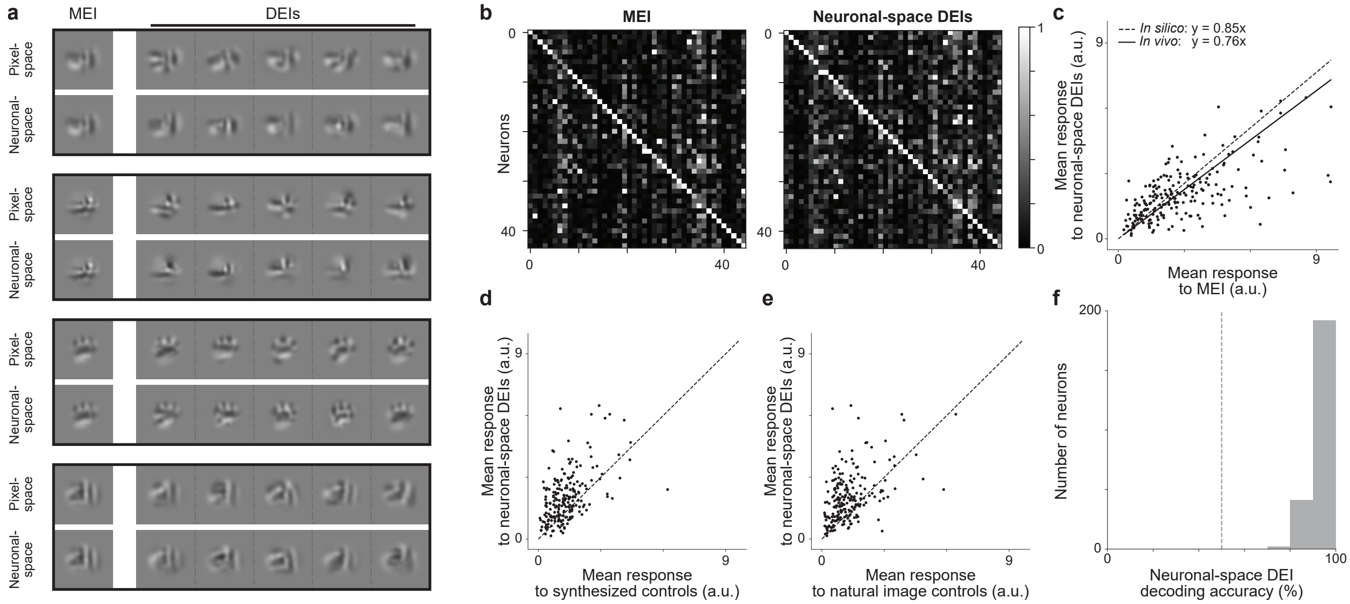


**Supplemental Fig. S7. Neuronal-space DEIs evoked strong and selective *in vivo* responses in target neurons while exhibiting population-decodable differences. a**, Examples of MEI and DEIs synthesized with image diversity evaluated by Euclidean distance in pixel space ("pixel-space") and cosine distance in *in silico* population neuronal response space ("neuronal-space") for 4 example neurons. **b**, Neuronal-space DEIs activated neurons with high specificity. The confusion matrices showed the responses of each neuron to MEI (left) and neuronal-space DEIs (right) of 44 neurons. MEI responses were averaged across 20 repeats of the same image while neuronal-space DEIs responses were averaged across 20 different images with single repeat. The responses of each neuron were normalized, and each row was scaled so the maximum response across all images equals $1$. Neurons responded more strongly to their own MEI and neuronal-space DEIs (along the diagonal) compared to other MEIs and neuronal-space DEIs respectively (two-sided permutation test, $P < 10^{-9}$ for both cases). **c**, Neuronal-space DEIs stimulated *in vivo* closely to the level predicted *in silico* with respect to MEI ($76 \pm 3\%$ versus $85\%$) (two-sided Wilcoxon signed-rank test, $W = 8464$, $P = 7.7 \times 10^{-3}$), with only $11.6\%$ of all neurons showing different responses between neuronal-space DEIs and $85\%$ of MEI ($0.48\%$ after BH correction) ($P < 0.05$, two-sided Welch's $t$-test with $32.61$ average d.f.). Data were pooled over 207 neurons from 5 mice. **d,e**, Neuronal-space DEIs activated their target neurons more strongly than synthesized and natural image controls (two-sided Wilcoxon signed-rank test, $W = 3441$, $P < 10^{-9}$ and $W = 3466$, $P < 10^{-9}$ respectively) with $13.5\%$ and $19.8\%$ of all neurons showing higher responses to neuronal-space DEIs ($0.0\%$ and $2.5\%$ after BH correction) ($P < 0.05$, two-sided Welch's $t$-test with $31.0$ and $30.6$ average d.f., respectively). **f**, *In vivo* population responses in mouse V1 Layer 2/3 discriminated between the most dissimilar pair of neuronal-space DEIs (based on cosine distance in *in silico* population neuronal response space) for each neuron. DEI identity in individual trials was decoded using a logistic regression classifier, with decoding accuracies across neurons (median = $93\%$) exceeded chance level ($50\%$, dashed line; one-sample $t$-test, $t = 138.5$, $P < 10^{-9}$). Data were pooled over 235 neurons from 3 mice.

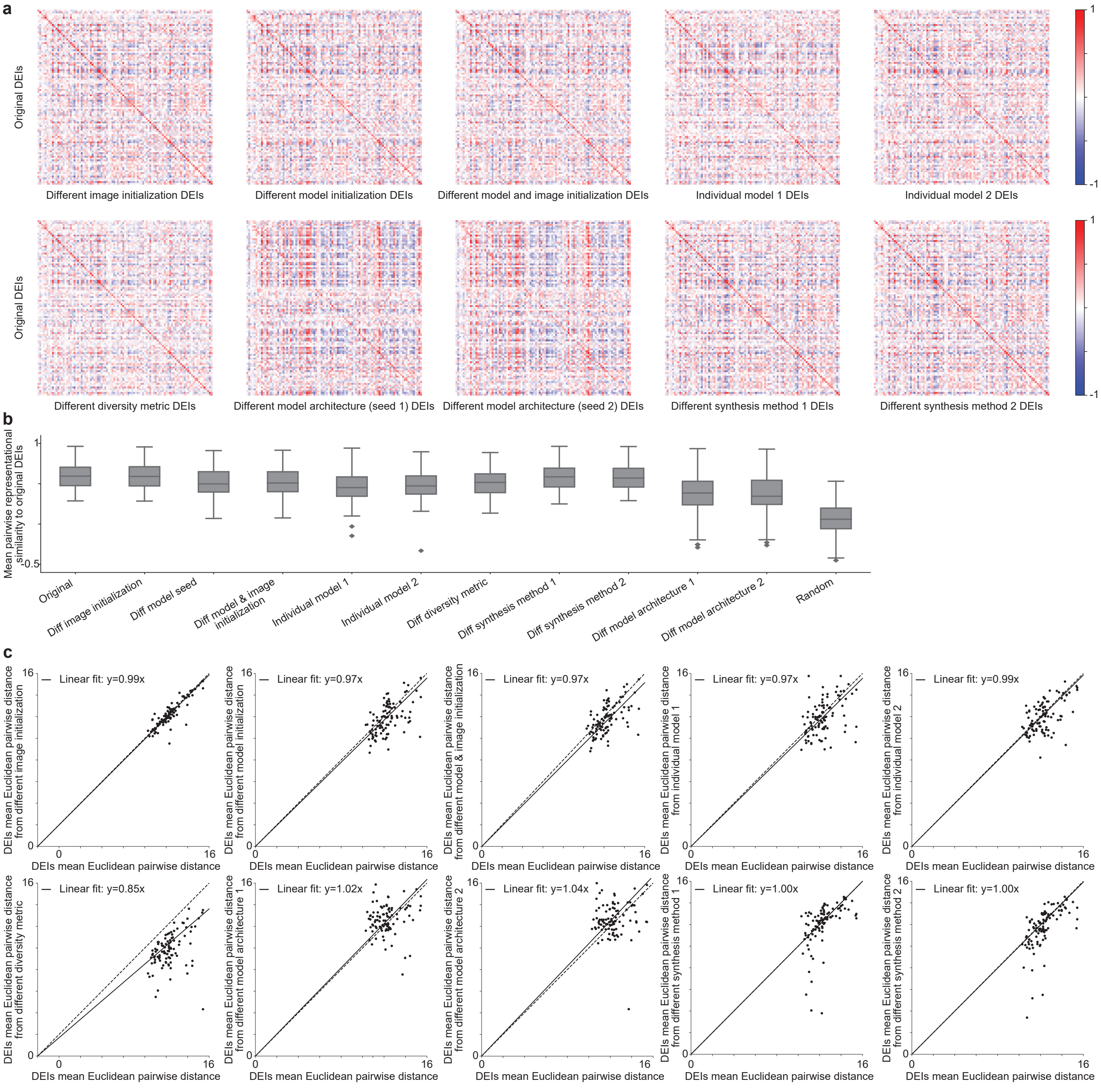


**Supplemental Fig. S8. DEIs generalized across different conditions. a**, DEIs synthesized under different conditions remained highly specific to their target neurons. The confusion matrices showed the representational similarity between the original DEIs and DEIs from (1) different image initialization for DEIs synthesis, (2) different model initialization, (3) different image and model initialization, (4,5) using single model from the ensemble for DEIs synthesis, (6) different diversity metric i.e. *in silico* population responses, (7,8) different model architecture (Willeke et al., 2022) with 2 different model seeds, (9,10) different synthesis method using implicit 1D and 2D periodic latent space neural representation model and contrastive regularization (Baroni et al., 2023). Each entry represents the mean pairwise cosine similarity between two sets of DEIs (see Methods for details). Representational similarity between original DEIs and DEIs synthesized from different conditions for the same neurons (diagonal) was larger than cross-neuron similarity (off-diagonal) (two-sided permutation test, $P < 10^{-9}$ for all conditions after BH correction). **b**, DEIs synthesized from different conditions were more similar to original DEIs than random neurons' DEIs (two-sided Wilcoxon signed-rank test, $W = 0, 0, 1, 1, 34, 0, 0, 0, 0, 0, 426$, and $357$, respectively, with $P < 10^{-9}$ for all conditions after BH correction). **c**, The mean Euclidean pairwise distances between DEIs generated using different methods were strongly correlated with those of the original DEIs (Pearson $r = 0.89, 0.56, 0.60, 0.43, 0.54, 0.45, 0.18, 0.08, 0.62$ and $0.66$, respectively with $P < 0.05$ for all conditions, two-sided $t$-test). Data were pooled over 97 neurons randomly sampled from 8 mice.

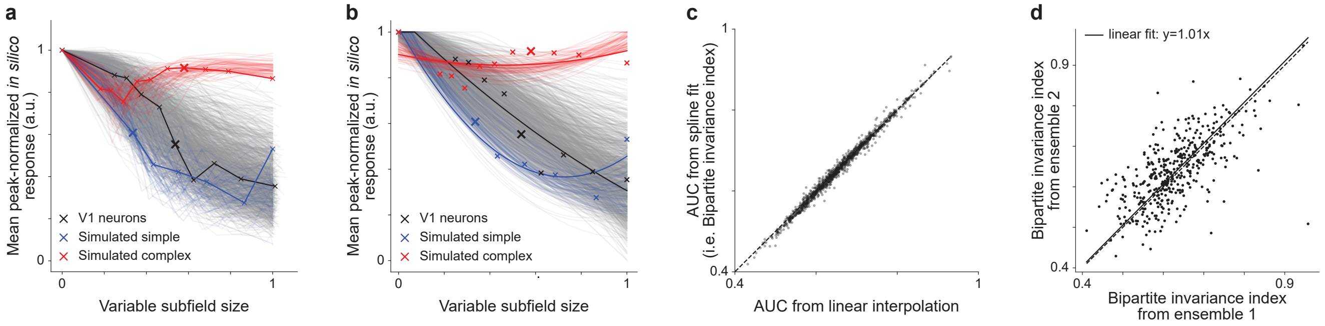


**Supplemental Fig. S9. Bipartite invariance quantification. a,b**, Mean peak-normalized *in silico* response as a function of the variable subfield size with linear interpolation (**a**) and quadratic-smoothing spline (**b**) for 1200 random V1 neurons from 6 mice, 60 simulated simple cells (blue), and 60 simulated complex cells (red). Responses were averaged across 20 random crops from the corresponding optimized texture and normalized to the MEI *in silico* response while the variable subfield sizes were normalized by the target neuron's MEI mask size. **c**, Bipartite invariance index was defined as the "Area Under the Curve" (AUC) from the quadratic-smoothing spline in **b**. AUC values were highly consistent regardless of whether linear interpolation or quadratic-smoothing spline was used (Pearson $r = 1.0, P < 0.05$, two-sided $t$-test). **d**, Bipartite invariance indices were consistent across different model initialization seeds (Pearson $r = 0.66$, $P < 0.05$, two-sided $t$-test). **a–d**, Data were pooled over 1200 neurons from 6 mice.

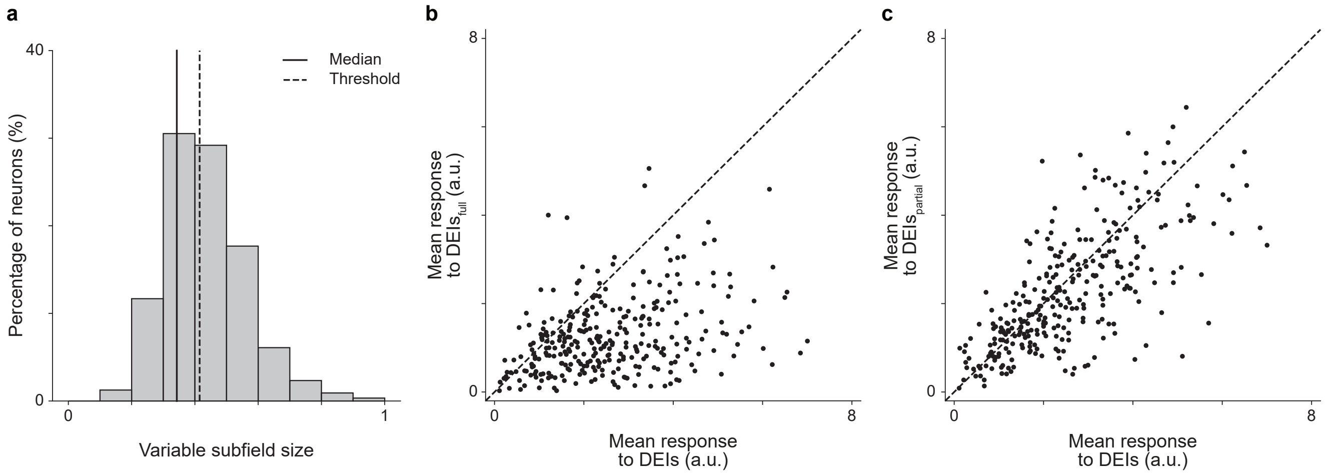


**Supplemental Fig. S10. Excluding neurons with small variable subfields did not alter the similarity between DEI and partial-texture DEI *in vivo* responses. a**, Histogram of variable subfield size for closed-loop neurons from Fig. 4e,f (median=$0.40$). **b, c**, Neurons with variable subfield sizes smaller than or equal to the 25th percentile ($0.34$) in **a** were excluded. **b**, $\text{DEIs}_{\text{full}}$ elicited weaker responses in their target neurons compared to DEIs (two-sided Wilcoxon signed-rank test, $W = 2853$, $P < 10^{-9}$) with $40.3\%$ of all neurons showing different responses ($25.7\%$ after BH correction) ($P < 0.05$, two-sided Welch's $t$-test with $30.1$ average d.f.). **c**, $\text{DEIs}_{\text{partial}}$ activated their target neurons similarly to DEIs (two-sided Wilcoxon signed-rank test, $W = 15920$, $P = 9.6 \times 10^{-6}$) with only $7.7\%$ of all neurons showing different responses ($0.0\%$ after BH correction) ($P < 0.05$, two-sided Welch's $t$-test with $33.6$ average d.f.). Data were pooled over 401 neurons from 8 mice.

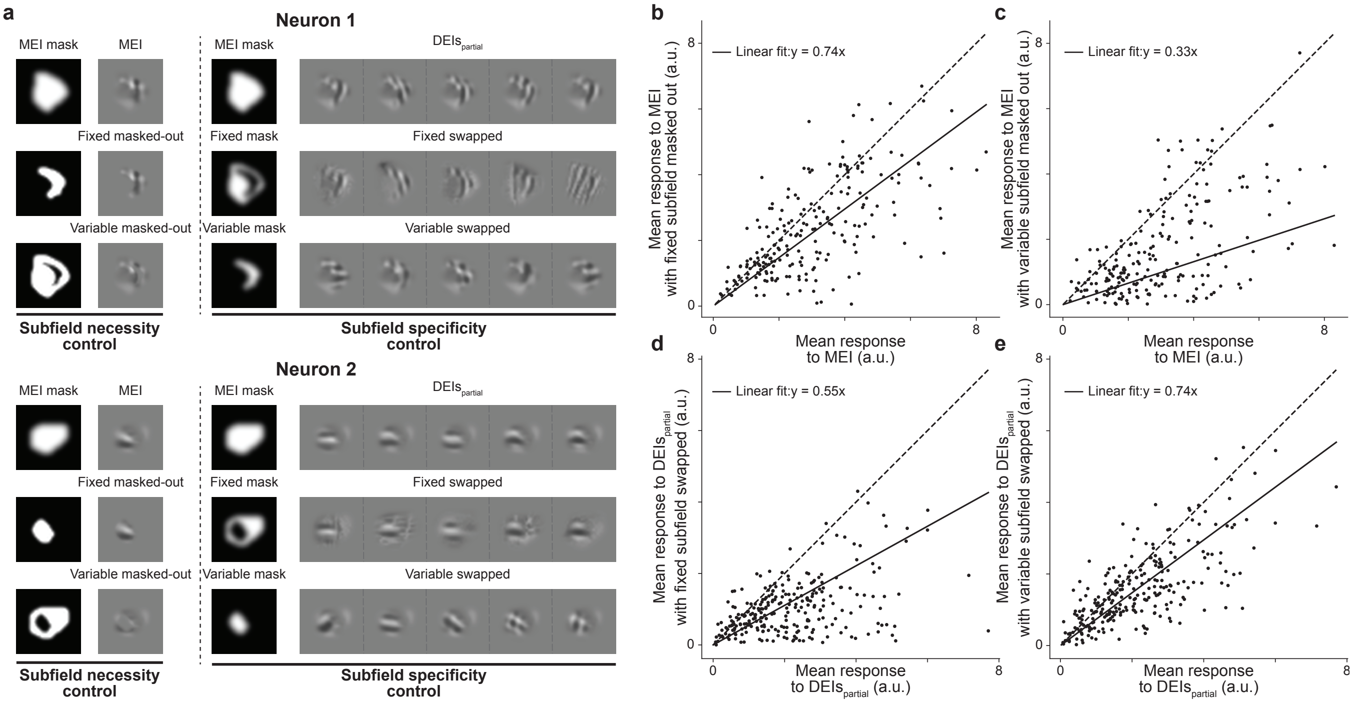


**Supplemental Fig. S11. Both subfields of the partial-texture DEIs are necessary and specific for evoking high *in vivo* responses.** **a**, Examples of the MEI, MEI with either fixed or variable subfield masked out, partial-texture DEIs ($\text{DEIs}_{\text{partial}}$), and $\text{DEIs}_{\text{partial}}$ with either fixed or variable subfield swapped with random content for two example neurons. The fixed subfield content was replaced with 20 different random natural patches while the variable subfield was replaced with patches cropped from 20 different random non-self neurons' preferred textures. When isolating a subfield, to avoid interruption of spatial pattern within it, we employed a soft-edged mask to mask out the complementary subfield. Consequently, part of the complementary subfield was preserved in the visual stimuli, making the subfield response contribution an underestimate of the actual effect. **b–e**, Each point corresponded to the response of a single neuron averaged over 20 repeats of its MEI or averaged over 20 different stimuli of the same type with single repeat. **b,c**, MEI with fixed or variable subfield masked out evoked weaker *in vivo* responses than MEI (two-sided Wilcoxon signed-rank test, $W = 5347$, $P < 10^{-9}$ and $W = 1570$, $P < 10^{-9}$, respectively) with $15.4\%$ and $38.1\%$ of all neurons showing weaker responses than their MEIs ($1.9\%$ and $27.9\%$ after BH correction) ($P < 0.05$, two-sided Welch's $t$-test with $33.2$ and $29.9$ average d.f., respectively). Data were pooled over 4 mice, displaying a total of 215 neurons. **d,e**, $\text{DEIs}_{\text{partial}}$ with either fixed or variable subfield swapped evoked weaker *in vivo* responses than $\text{DEIs}_{\text{partial}}$ (two-sided Wilcoxon signed-rank test, $W = 1960$, $P < 10^{-9}$, and $W = 5820$, $P < 10^{-9}$, respectively) with $30.1\%$ and $12.7\%$ of all neurons showing weaker responses than their partial-texture DEIs ($18.9\%$ and $0.0\%$ after BH correction) ($P < 0.05$, two-sided Welch's $t$-test with $30.0$ and $32.5$ average d.f., respectively). Data were pooled over 259 neurons from 5 mice.

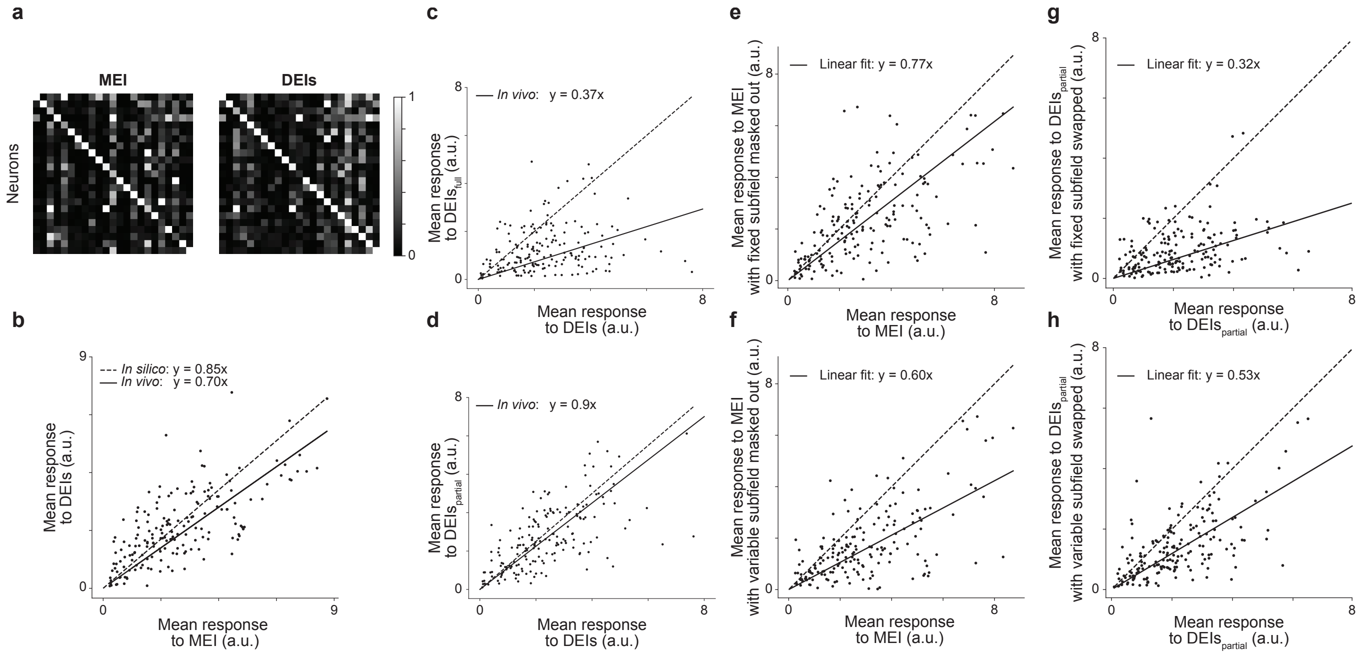


**Supplemental Fig. S12. DEI closed-loop verification for randomly selected neurons.** **a**, Confusion matrices for MEI and DEIs showed the responses of each neuron to the MEI (left) and DEIs (right) across all target neurons. The responses of each neuron were normalized, and each row was scaled so the maximum response across all images equaled $1$. Responses of neurons to their own MEI and DEIs (along the diagonal) were larger than to other MEIs and DEIs (two-sided permutation test, $P < 10^{-9}$ for both cases). **b–h**, Each point represented the average response of a single neuron over 20 repeats of its MEI or 20 different stimuli of the same type with single repeat. **b**, DEIs stimulated neurons *in vivo* closely to the level predicted *in silico* with respect to MEI ($70 \pm 6\%$ versus $85\%$) (two-sided Wilcoxon signed-rank test, $W = 7257$, $P = 0.25$), with only $9.5\%$ neurons showing different responses between DEIs and $85\%$ of MEI ($0.0\%$ after BH correction) ($P < 0.05$, two-sided Welch's $t$-test with $32.5$ average d.f.). **c**, Full-texture DEIs ($\text{DEIs}_{\text{full}}$) evoked weaker responses in their target neurons than DEIs (two-sided Wilcoxon signed-rank test, $W = 1988$, $P < 10^{-9}$) with $35.2\%$ of all neurons showing different responses than DEIs ($15.1\%$ after BH correction) ($P < 0.05$, two-sided Welch's $t$-test with $29.4$ average d.f.) **d**, Partial-texture DEIs ($\text{DEIs}_{\text{partial}}$) activated their target neurons similarly to DEIs (two-sided Wilcoxon signed-rank test, $W = 6693$, $P = 0.05$) with only $8.9\%$ of neurons showing different responses from corresponding DEIs ($0.0\%$ after BH correction) ($P < 0.05$, two-sided Welch's $t$-test with $33.1$ average d.f.). **e, f**, MEIs with either fixed or variable subfields masked out evoked weaker responses in their target neurons than MEIs (two-sided Wilcoxon signed-rank test, $W = 4484$, $P < 10^{-9}$, and $W = 1401$, $P = 2.7 \times 10^{-7}$, respectively) with $20.1\%$ and $31.3\%$ of neurons showing weaker responses than MEIs ($3.9\%$ and $16.2\%$ after BH correction) ($P < 0.05$, two-sided Welch's $t$-test with $32.6$ and $31.1$ average d.f., respectively). **g, h**, $\text{DEIs}_{\text{partial}}$ with either fixed or variable subfield swapped evoked weaker responses in their target neurons than $\text{DEIs}_{\text{partial}}$ (two-sided Wilcoxon signed-rank test, $W = 1005$, $P < 10^{-9}$, and $W = 2026$, $P < 10^{-9}$, respectively) with $43.0\%$ and $17.3\%$ neurons showing weaker responses than $\text{DEIs}_{\text{partial}}$ ($22.9\%$ and $0.6\%$ after BH correction) ($P < 0.05$, two-sided Welch's $t$-test with $28.0$ and $31.2$ average d.f., respectively). Data were pooled over 179 neurons from 3 mice.

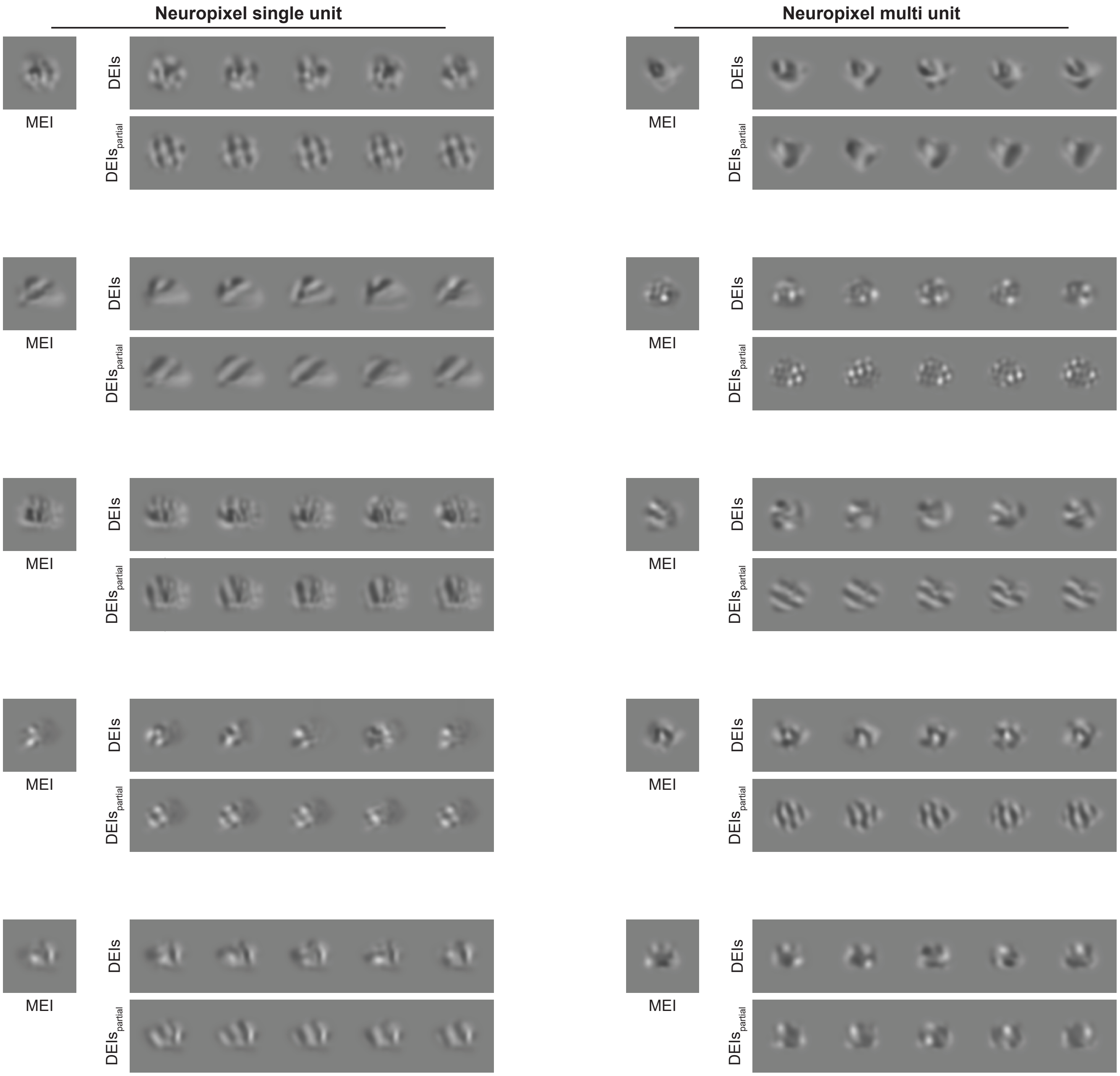


**Supplemental Fig. S13. Example MEI, DEIs, and partial-texture DEIs from electrophysiological recordings for units classified as “single” (left) and “multiple” (right) based on spike sorting.**

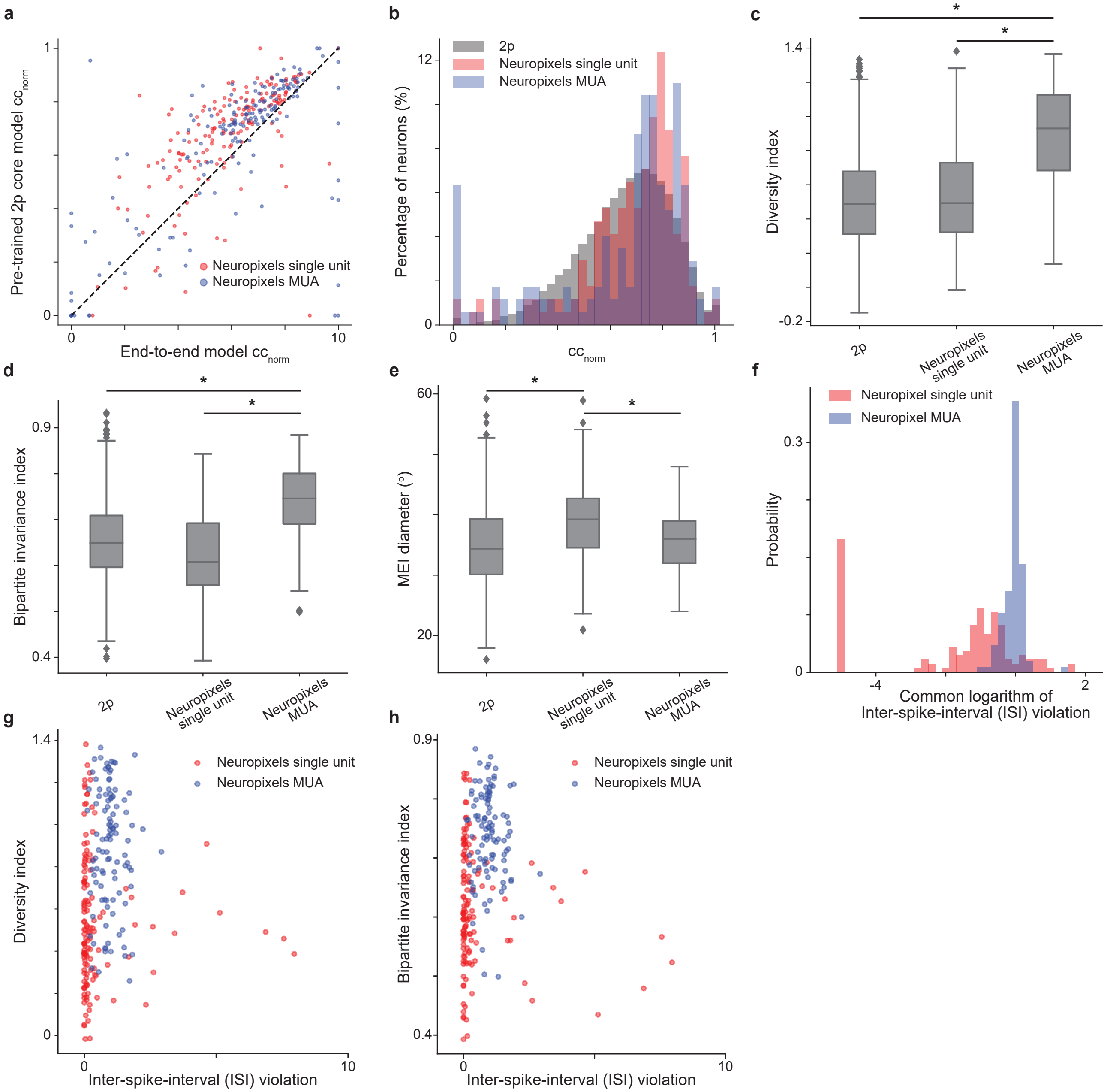


**Supplemental Fig. S14. Quantification of diversity and bipartite invariance indices from electrophysiological data. a**, Normalized correlation coefficients ($CC_{norm}$) of the model fine-tuned from a pre-trained core on two-photon recordings (denoted as "pre-trained 2p core") were higher than those from the model trained from scratch (denoted as "end-to-end") (two-sided Wilcoxon signed-rank test, $W = 10762$, $P < 10^{-9}$) with color representing units classified as either "single units" or "multi-unit activity" (denoted as "Neuropixels single" and "Neuropixels MUA", respectively). **b**, Histogram of the normalized correlation coefficient ($CC_{norm}$) for neurons modeled from two-photon recording (denoted as "2p", data were pooled over 33,714 neurons from 14 mice), and Neuropixels units modeled using electrophysiological data with pre-trained 2p core (data were pooled over 364 spikes sorted units from 6 mice). Excessively noisy neurons ($CC_{max} < 0.1$) were excluded ($0.2\%$, $7.6\%$, and $4.6\%$ of 2p, Neuropixels single, Neuropixels MUA, respectively) and neurons with values outside of $0$ and $1$ were clipped ($1.2\%$, $4.5\%$, and $6.1\%$ for 2p, Neuropixels single, and Neuropixels MUA, respectively) for visualization. **c**, Neuropixels MUA exhibited larger diversity indices than Neuropixels single units and 2p neurons ($P < 0.05$, two-sided Welch's $t$-test with $230.3$ and $122.6$ d.f., respectively) while Neuropixels single units had similar diversity indices to 2p neurons ($P = 0.21$, two-sided Welch's $t$-test with $158.4$ d.f.). **d**, Neuropixels MUA showed larger bipartite invariance indices than Neuropixels single units and 2p neurons ($P < 0.05$, two-sided Welch's $t$-test with $237.8$ and $129.5$ d.f., respectively) while Neuropixels single units had marginally lower bipartite invariance indices than 2p neurons ($P < 0.05$, two-sided Welch's $t$-test with $155.7$ d.f.). **e**, Neuropixels single units had larger RF sizes than both Neuropixels MUA and 2p neurons ($P < 0.05$, two-sided Welch's $t$-test with $235.5$ and $161.3$ d.f., respectively) while MUA had similar RF sizes to those of 2p neurons ($P = 0.22$, two-sided Welch's $t$-test with $140.9$ d.f.). **f**, Histogram of the common logarithm of the inter-spike-interval (ISI) violations for Neuropixels single and Neuropixels MUA. Neuropixels MUA exhibited high ISI violations (median=$0.94$), whereas Neuropixels single units displayed a smaller mean but broader distribution of ISI violations (median=$0.09$). **g**, Diversity indices were not correlated with the ISI violations for Neuropixels single units (Pearson's $r = -0.03$ with $p = 0.76$). **h**, Bipartite invariance indices were not correlated with the ISI violations for Neuropixels single units (Pearson's $r = -0.05$ with $p = 0.55$). **g, h**, Units with excessively large ISI violation ($> 10$) were excluded for visualization (2 single units and 1 MUA). **c–h**, Neurons were selected to have oracle score larger than $0.22$ and model test correlation larger than $0.42$. Two-photon data were pooled over 1,154 neurons from 6 mice; electrophysiological data pooled over 240 spike sorted units from 6 mice.

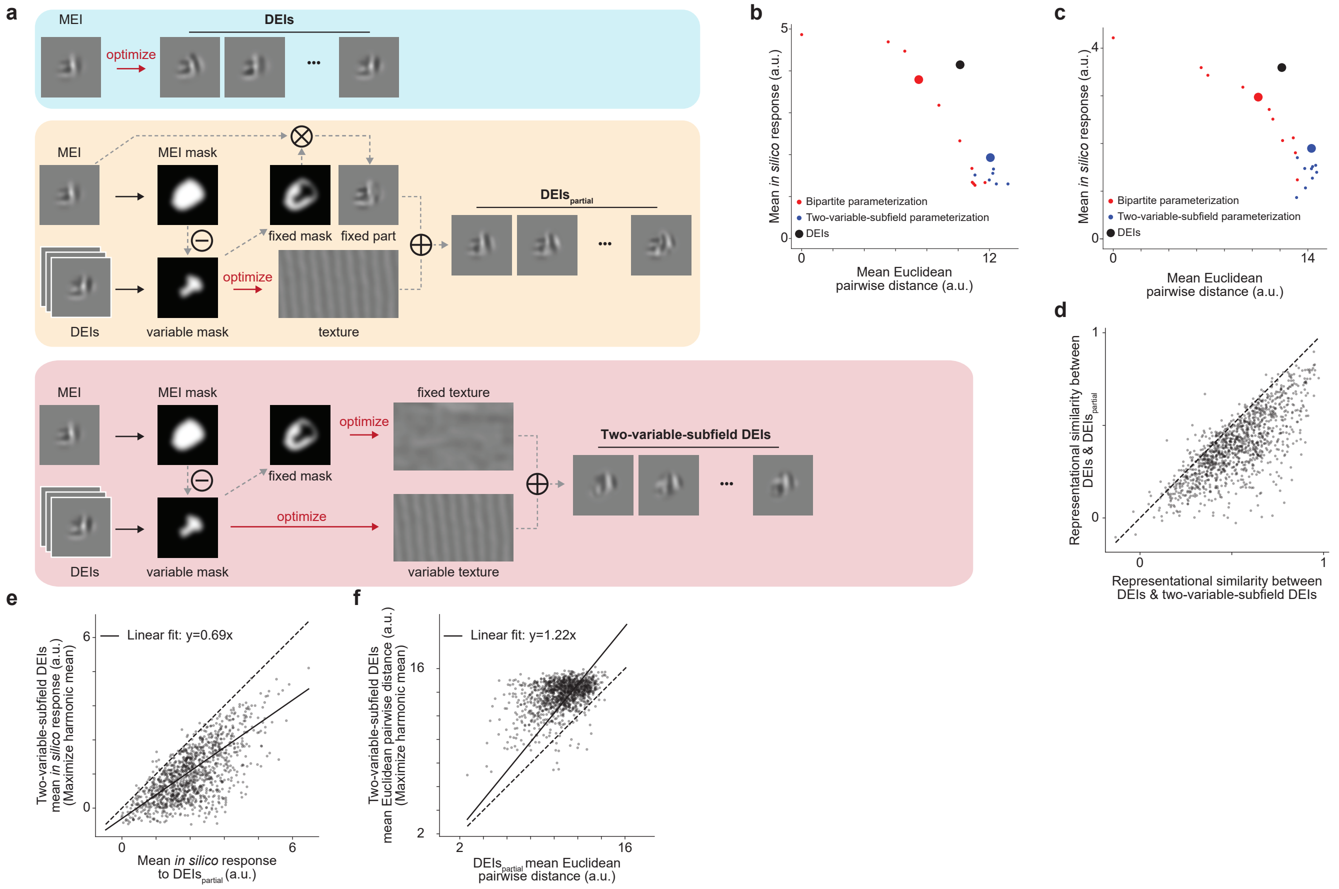


**Supplemental Fig. S15. DEIs cannot be well explained by shift invariance in both subfields. a**, Schematic illustrating three DEIs generation methods for an example V1 bipartite cell: non-parametric optimization (DEIs), bipartite parameterization (DEIs$_{\text{partial}}$), and "two-variable-subfield" parameterization. To synthesize two-variable-subfield DEIs, we followed the same procedure for DEIs$_{\text{partial}}$, but treated both subfields as shift-invariant and optimized two distinct texture images jointly for each subfield. **b, c**, Example of *in silico* response and mean Euclidean pairwise distance for DEIs optimized using bipartite and two-variable-subfield parameterization. Similar to DEIs$_{\text{partial}}$, we selected the sets of DEIs from two-variable-subfield parameterization to maximize the harmonic mean between *in silico* response and image diversity (denoted as "two-variable-subfield DEIs" and indicated by larger dots). **d**, DEIs$_{\text{partial}}$ were more similar to non-parametric DEIs than two-variable-subfield DEIs as measured by representational similarity (two-sided Wilcoxon signed-rank test, $W = 23640$, $P < 10^{-9}$). **e**, DEIs$_{\text{partial}}$ evoked higher responses in their target neurons *in silico* than two-variable-subfield DEIs (two-sided Wilcoxon signed-rank test, $W = 1977$, $P < 10^{-9}$). **f**, DEIs$_{\text{partial}}$ exhibited lower diversity than two-variable-subfield DEIs (two-sided Wilcoxon signed-rank test, $W 4842$, $P < 10^{-9}$). Data were pooled over 1200 randomly selected neurons from 6 mice.

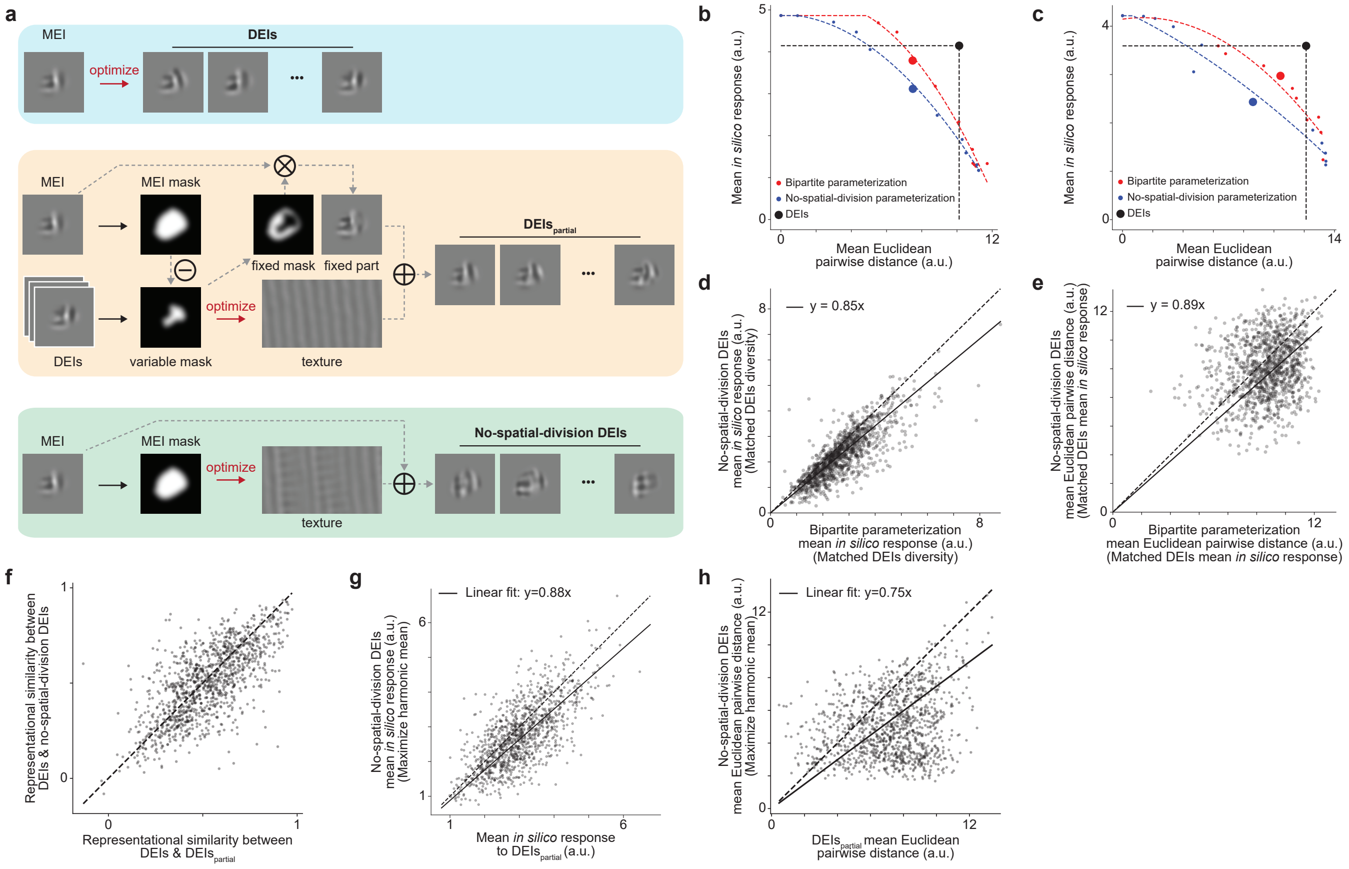


**Supplemental Fig. S16. Spatial division is necessary for explaining DEIs. a**, Schematic illustrating three DEIs generation methods for an example V1 bipartite cell: non-parametric optimization (DEIs), bipartite parameterization ($\text{DEIs}_{\text{partial}}$), and "no-spatial-division" parameterization. No-spatial-division model parameterize DEIs as the summation of two fully superimposed subfields, unlike the non-overlapping subfields in the bipartite parameterization. **b, c**, Example of *in silico* response and mean Euclidean pairwise distance for DEIs optimized using bipartite and no-spatial-division parameterization. Similar to $\text{DEIs}_{\text{partial}}$, we selected the sets of DEIs from no-spatial-division parameterization to maximize the harmonic mean between *in silico* response and image diversity (denoted as "no-spatial-division DEIs" and indicated by larger dots). For each parameterization, a quadratic-smoothing spline fit was used to estimate: 1) the *in silico* response at the diversity level matching non-parametric DEIs (denoted as "matched DEIs diversity") and 2) the diversity level at the *in silico* response matching non-parametric DEIs' response ("matched DEIs mean *in silico* response") **d**, Bipartite parameterization evoked higher *in silico* responses than no-spatial-division parameterization when matched DEIs diversity (two-sided Wilcoxon signed-rank test, $W = 82385$, $P < 10^{-9}$). **e**, Bipartite parameterization showed greater diversity than no-spatial-division parameterization when matched DEIs mean *in silico* response (two-sided Wilcoxon signed-rank test, $W = 89053$, $P < 10^{-9}$). **f**, $\text{DEIs}_{\text{partial}}$ were more similar to the original non-parametric DEIs than no-spatial-division DEIs as measured by representational similarity (two-sided Wilcoxon signed-rank test, $W = 293622$, $P = 2.8 \times 10^{-8}$). **g**, $\text{DEIs}_{\text{partial}}$ evoked stronger *in silico* responses than no-spatial-division DEIs (two-sided Wilcoxon signed-rank test, $W = 199252$, $P < 10^{-9}$). **h**, $\text{DEIs}_{\text{partial}}$ exhibited higher diversity compared to no-spatial-division DEIs (two-sided Wilcoxon signed-rank test, $W = 241714$, $P < 10^{-9}$). Data were pooled over 1200 randomly selected neurons from 6 mice.

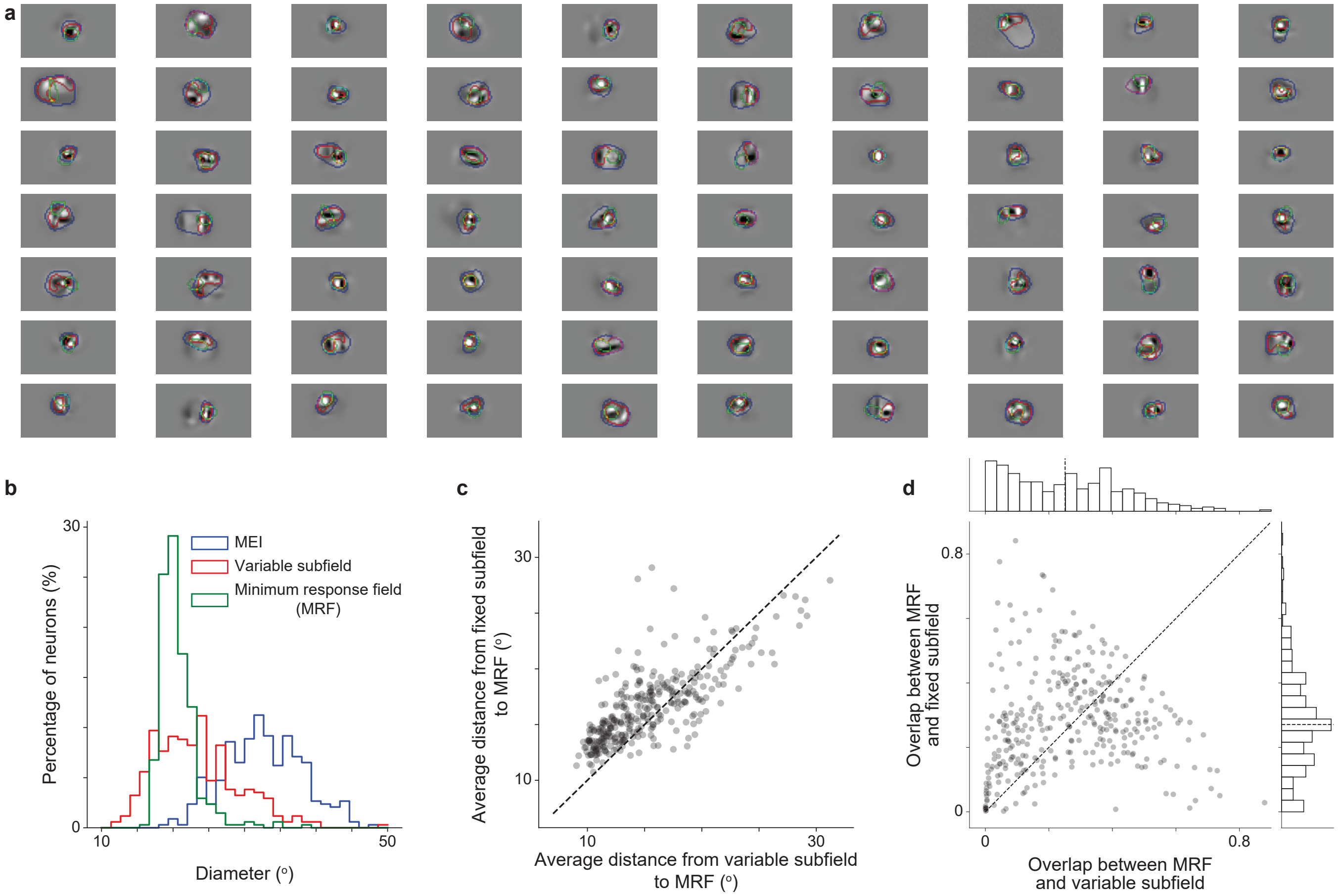


**Supplemental Fig. S17. Bipartite receptive field cannot be explained by the center-surround structure. a**, Examples MEIs overlaid with their boundary (blue), variable subfield boundaries (red), and "Minimum Response Fields" (MRF) boundaries (green) for visualization. **b**, Histogram of MEI, MRF, and variable subfield diameters. The mean diameters of MEI, MRF, and variable subfield across all neurons were (mean $\pm$ s.e.m.): $32.9 \pm 0.02$, $20.6 \pm 0.01$, $23.3 \pm 0.02$ degrees. **c**, Fixed subfield is further from MRF than variable subfield quantified by average pairwise distance between the MRF and each of the two subfields across all pixels (two-sided Wilcoxon signed-rank test, $W = 14278,\ P < 10^{-9}$). **d**, However, fixed subfield overlapped more with MRF than variable subfield (two-sided Wilcoxon signed-rank test, $W = 24373,\ P = 0.01$; median = $27.1\%$ and $25.1\%$, respectively). Data were pooled over 340 neurons from 2 mice.

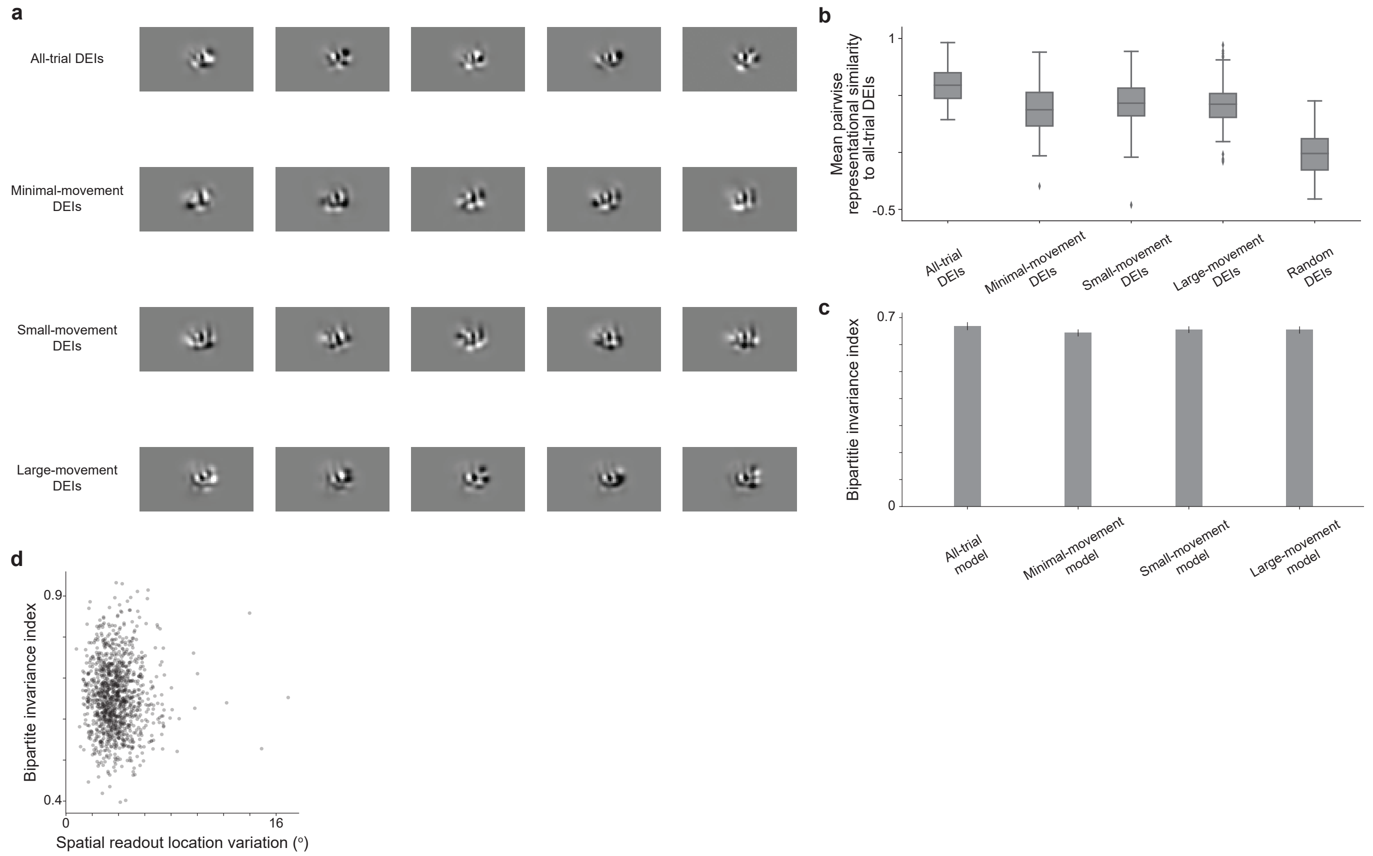


**Supplemental Fig. S18. Bipartite receptive field cannot be explained by neither trial-to-trial eye movement nor spatial readout location variation.** **a**, We created three sub-datasets from full-field natural image trials using inclusion criteria based on different thresholds of eye movement size: 1) minimal-movement trials only, 2) small-movement trials only, and 3) large-movement trials only (see Methods for details). For each sub-dataset, an additional model was trained for the same neuron, in addition to the original model trained on all trials. Examples of DEIs from 4 models for an example neuron were shown for visualization. **b**, DEIs synthesized from all three additional models were more similar to original DEIs than to random neuron DEIs as measured by the representational similarity (two-sided Wilcoxon signed-rank test, $W = 213, 121$, and $101$, respectively, with $P < 10^{-9}$ for all conditions after BH correction). **c**, Bipartite invariance indices were similar across all four models (one-way ANOVA $F = 0.97, P = 0.38$). **d**, Bipartite invariance indices were not correlated with the spatial readout location variation (Pearson $r = 0.037$, $P = 0.99$). Spatial readout location variation was defined as the mean Euclidean pairwise distance between individual model readout locations in visual degree. Data were pooled over 1200 randomly selected neurons from 6 mice.

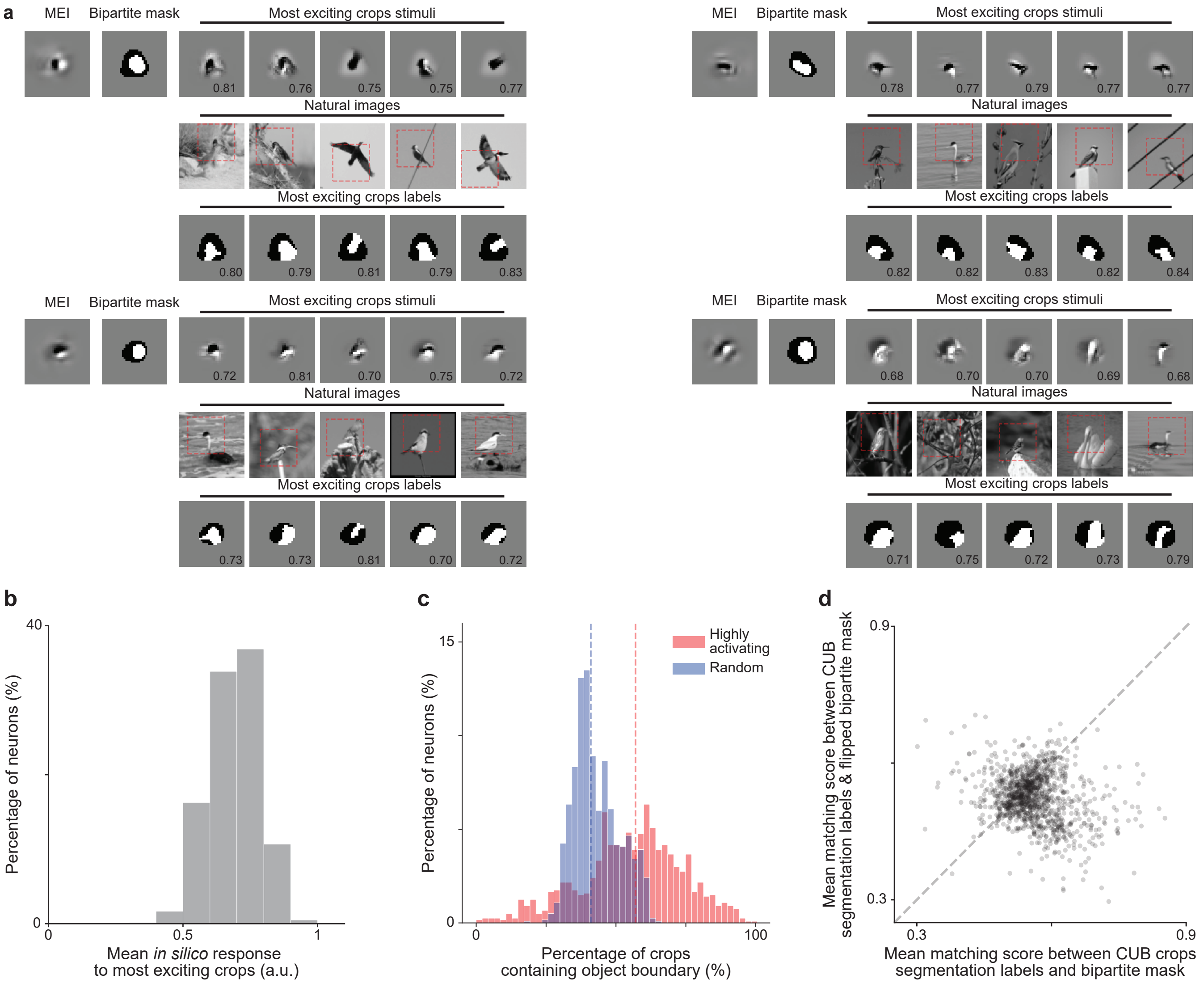


**Supplemental Fig. S19. DEI bipartite masks aligned with object boundaries in highly activating natural crops.** **a**, For each of the 4 example neurons, we showed the MEI, the bipartite mask (black denotes the fixed subfield and white denotes the variable subfield), along with 5 randomly selected crops from the top 100 most activating natural crops, their corresponding full-field natural images, and the segmentation labels. Red dashed boxes on the full-field natural images indicated the regions where the crops were taken from. We showed the *in silico* response normalized by MEI *in silico* response at the corner of each crop and the corresponding matching score at the bottom of the segmentation label. **b**, Highly activating natural crops were more likely to contain object boundaries compared to random crops (two-sided Wilcoxon signed-rank test, $W = 573606,\ P < 10^{-9}$). **c**, Histogram of mean *in silico* response for the top 100 most activating natural crop, normalized by the corresponding MEI *in silico* response (median=$0.70$). **d**, Flipping the bipartite mask with respect to the center of the receptive field (RF) decreased the matching score for the highly activating natural crops (two-sided Wilcoxon signed-rank test $W = 495850,\ P < 10^{-9}$) with $41.9\%$ of all neurons showing weaker matching scores than original bipartite mask ($37.6\%$ after BH correction) ($P < 0.05$, two-sided Welch's $t$-test with $99.0$ average d.f.). One neuron was excluded from the analysis as it strictly preferred patches without object boundary. Data were pooled over 1200 randomly selected neurons from 6 mice.

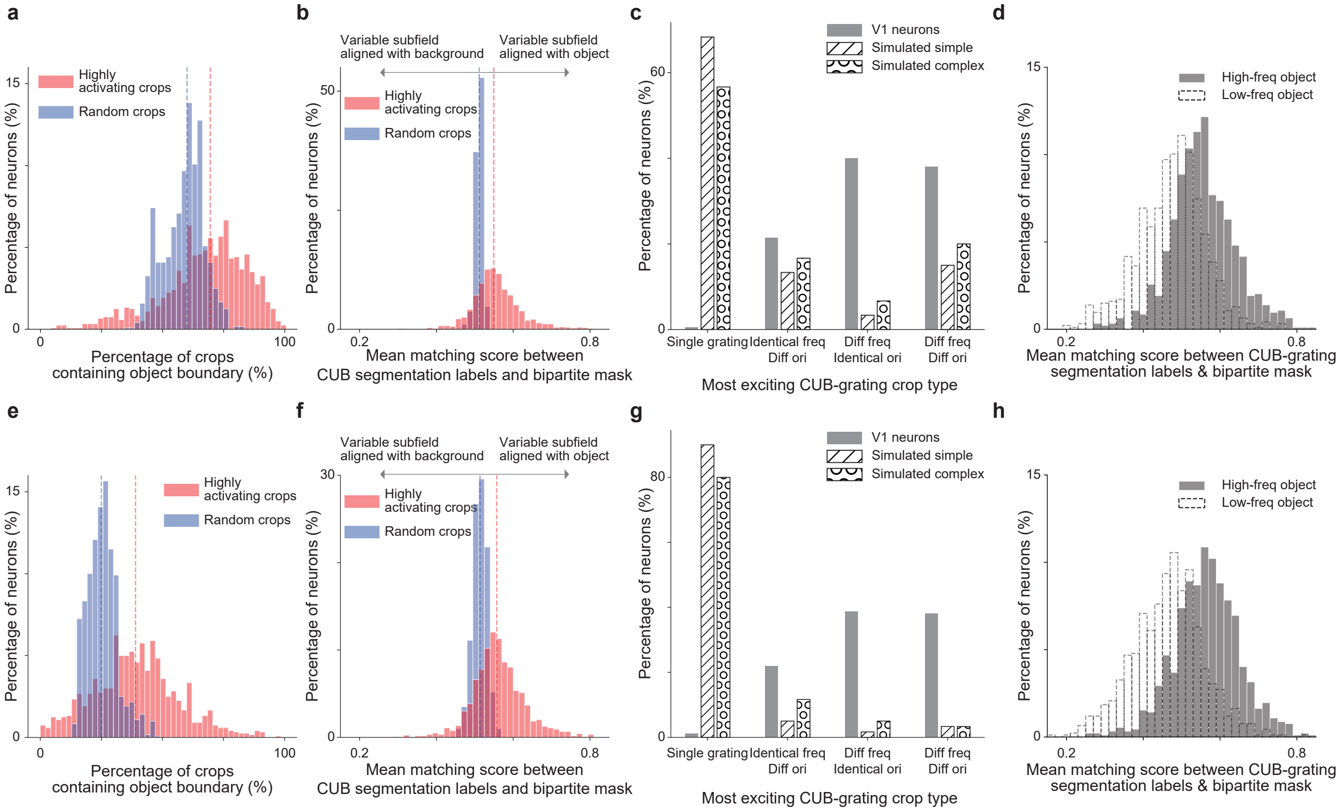


**Supplemental Fig. S20. Alignment between bipartite mask and natural object boundaries was robust across different thresholds for classifying patches containing object boundaries.** We systematically varied the minimum fraction of both object and background required within the receptive field (RF) to classify a patch as containing an object boundary. **a–d, e–h**, Patches were classified as containing an object boundary if they contained more than 10% (Condition 1) or 30% (Condition 2) of both object and background. **a, e**, Highly activating natural crops were more likely to contain object boundaries compared to randomly crops (two-sided Wilcoxon signed-rank test, $W = 545046$, $P < 10^{-9}$ and $W = 603265$, $P < 10^{-9}$ respectively). **b, f**, Highly activating CUB crops with object boundary yielded higher matching scores than those for random natural crops with object boundary (two-sided Wilcoxon signed-rank test, $W = 85378$, $P < 10^{-9}$ and $W = 92707$, $P < 10^{-9}$) with $54.7\%$ and $44.7\%$ of all neurons showing greater matching scores to highly activating crops than to random crops ($50.8\%$ and $38.2\%$ after BH correction) while only $5.3\%$ and $4.4\%$ of all neurons showing smaller matching scores to highly activating crops ($4.6\%$ and $3.3\%$ after BH correction) ($P < 0.05$, two-sided Welch's $t$-test with $107.2$ and $42.3$ average d.f. respectively). Six cells were excluded from condition 2 for this analysis since they preferred patches without object boundaries (fewer than 2 of the top 100 most activating crops containing object boundaries). **c, g**, Simulated simple and complex cells predominantly prefer "single grating" image type ($68.3\%$, $56.7\%$ for condition 1 and $90.0\%$, $80.0\%$ for condition 2, respectively). In contrast, V1 neurons exhibited a distinct pattern with only $0.5\%$ and $1.2\%$ of V1 neurons prefer "single grating" for two conditions, respectively (one-way chi-squared test, $\chi^2 = 6152$, $P < 10^{-9}$, $\chi^2 = 2889$, $P < 10^{-9}$ and $\chi^2 = 15994$, $P < 10^{-9}$, $\chi^2 = 8123$, $P < 10^{-9}$ for comparison against simulated simple and complex cells in two conditions respectively). Instead, most neurons preferred "different frequency but identical orientation" ($40.1\%$ and $38.8\%$), followed by "different frequency and orientation" ($38\%$ and $38.1\%$), and "identical frequency but different orientation" ($21.4\%$ and $22.0\%$) for two conditions, respectively. **d, h**, Among neurons preferring different frequencies within the RF, matching score for highly activating crops were higher in "high-frequency object" dataset than "low-frequency object" dataset (two-sided Wilcoxon signed-rank test, $W = 336493$, $P < 10^{-9}$ and $W = 338329$, $P < 10^{-9}$ respectively) with $64.6\%$ and $67.2\%$ of all neurons showing greater matching scores ($64.6\%$ and $67.0\%$ after BH correction) while only $24.7\%$ and $19.5\%$ of all neurons showing smaller matching scores ($24.2\%$ and $19.0\%$ after BH correction) ($P < 0.05$, two-sided Welch's $t$-test with $182.8$ and $145.5$ average d.f., respectively). Simulated simple and complex cells composed of 60 neurons for each population. Data were pooled over 1200 randomly selected neurons from 6 mice.

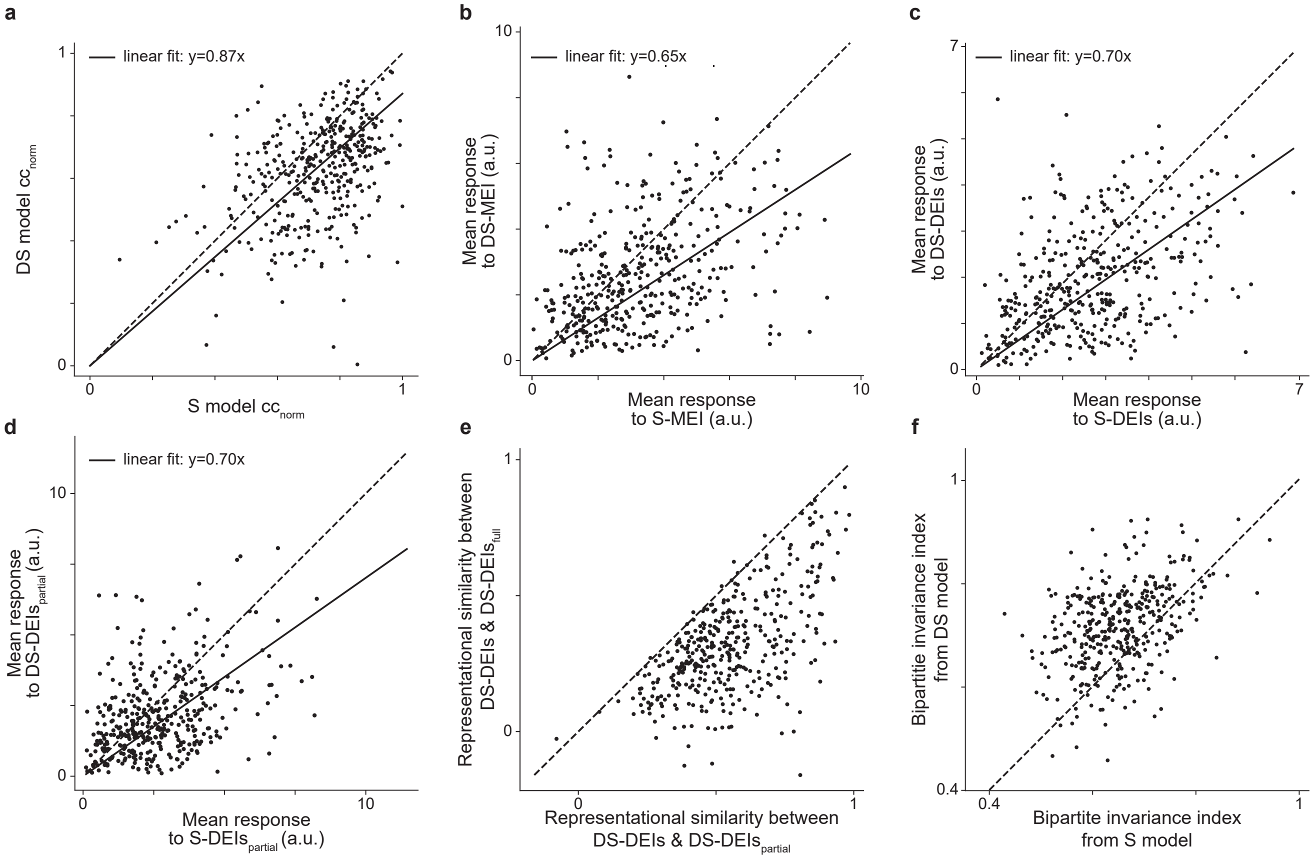


**Supplemental Fig. S21. Dynamic static model *in vivo* validation a**, Both static and dynamic-static model yield high normalized correlation coefficient ($CC_{norm}$) for all well-matched neurons (median=$0.75, 0.65$, respectively). **b**, Both static MEI and dynamic-static MEI evoked high *in vivo* responses but static MEI evoked larger responses in target neurons than dynamic-static MEI (two-sided Wilcoxon signed-rank test, $W = 24493$, $P < 10^{-9}$) with $22.3\%$ of all neurons showing higher responses to static MEI while $6.3\%$ showing lower responses ($10.8\%$ and $2.3\%$ after BH correction) ($P < 0.05$, two-sided Welch's $t$-test with $32.0$ average d.f.). **c**, Similarly, static DEIs evoked larger *in vivo* responses in target neurons more than dynamic-static DEIs (two-sided Wilcoxon signed-rank test, $W = 21414$, $P < 10^{-9}$) with $14.8\%$ of all neurons showing higher responses to static DEIs while $2.5\%$ showing lower responses ($2.5\%$ and $0.3\%$ after BH correction) ($P < 0.05$, two-sided Welch's $t$-test with $32.1$ average d.f.). **d**, Static DEIs$_{partial}$ drove target neurons more than dynamic-static DEIs$_{partial}$ (two-sided Wilcoxon signed-rank test, $W = 25738?$, $P < 10^{-9}$) with $16.3\%$ of all neurons showing higher responses to static DEIs$_{partial}$ while $5.5\%$ showing lower responses ($1.8\%$ and $0.3\%$ after BH correction) ($P < 0.05$, two-sided Welch's $t$-test with $31.5$ average d.f.). **e**, Dynamic-static DEIs$_{partial}$ were more similar to dynamic-static non-parametric DEIs than dynamic-static DEIs$_{full}$ (two-sided Wilcoxon signed-rank test, $W = 180$, $P < 10^{-9}$). **f**, Bipartite invariance indices from the dynamic-static were highly correlated with those from the static model (Pearson $r = 0.66, P < 0.05$, two-sided $t$-test). Data was collected from 3 mice, displaying a total of 399 neurons.

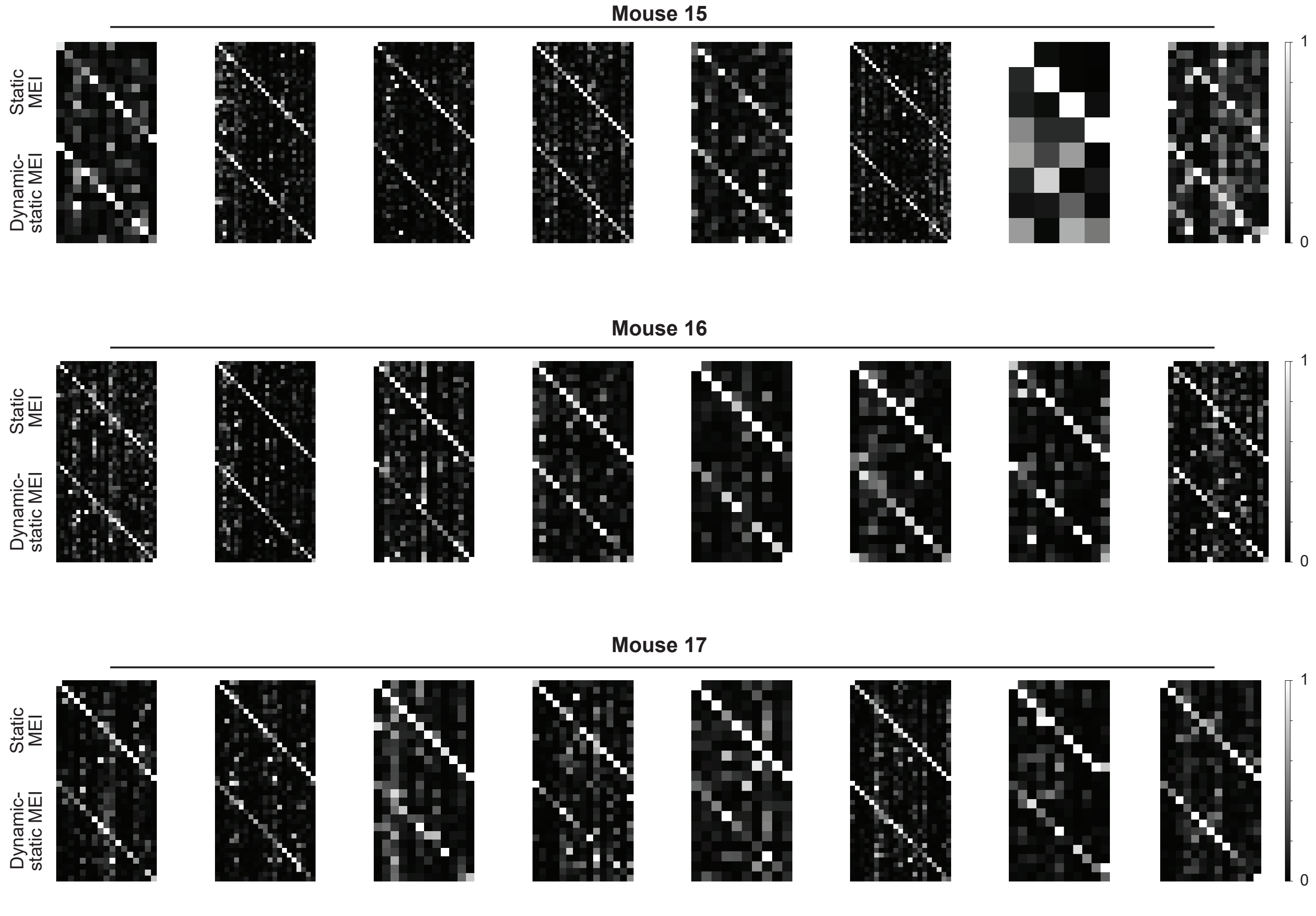


**Supplemental Fig. S22. MEI activated neurons with high specificity in both static and dynamic-static models.** The confusion matrices showed the responses of each neuron to static MEI (top) and dynamic-static MEI (bottom) of all target neurons in individual scans where we presented both models' stimuli back to the mouse in closed-loop experiments. MEI responses were averaged across 20 repeats of the same image. The responses of each neuron were normalized, and each row was scaled so the maximum response across all images equals $1$. Responses of neurons to their own MEI (along the diagonal) were larger than to other MEIs (two-sided permutation test, $P < 10^{-4}$ across all mice after BH correction).

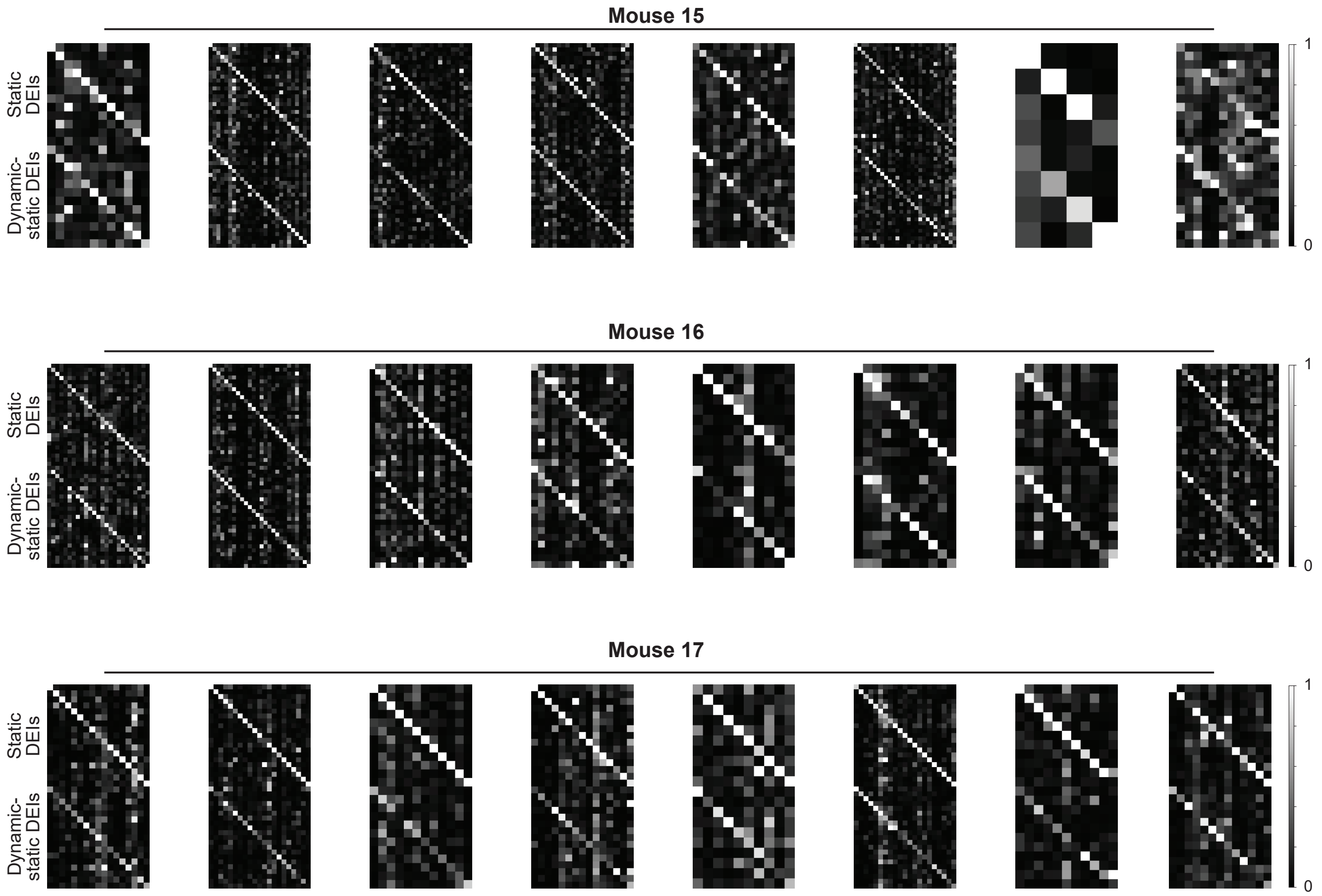


**Supplemental Fig. S23. DEIs activated neurons with high specificity in both static and dynamic-static models.** The confusion matrices showed the responses of each neuron to static DEIs (top) and dynamic-static DEIs (bottom) of all target neurons in individual scans where we presented both models' stimuli back to the mouse in closed-loop experiments. DEI responses were averaged across 20 different images with single repeat. The responses of each neuron were normalized, and each row was scaled so the maximum response across all images equals $1$. Responses of neurons to their own DEIs (along the diagonal) were larger than to other DEIs (two-sided permutation test, $P < 10^{-4}$ across all mice after BH correction).

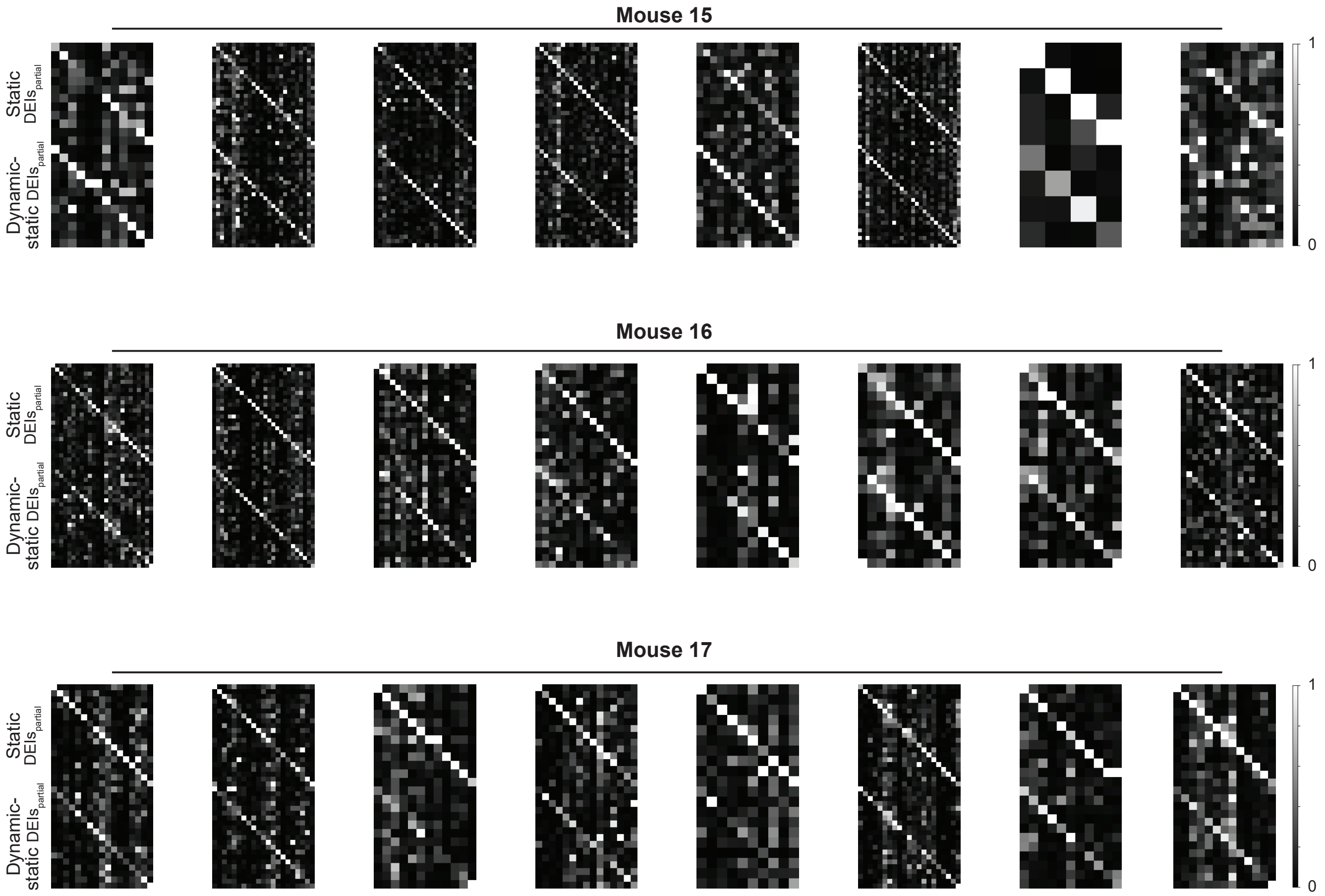


**Supplemental Fig. S24. Partial-texture DEIs activated neurons with high specificity in both static and dynamic-static models.** The confusion matrices showed the responses of each neuron to static DEIs$_{partial}$ (top) and dynamic-static DEIs$_{partial}$ (bottom) of all target neurons in individual scans where we presented both models' stimuli back to the mouse in closed-loop experiments. DEI$_{partial}$ responses were averaged across 20 different images with single repeat. The responses of each neuron were normalized, and each row was scaled so the maximum response across all images equals $1$. Responses of neurons to their own DEIs$_{partial}$ (along the diagonal) were larger than to other DEIs$_{partial}$ (two-sided permutation test, $P < 10^{-4}$ across all mice after BH correction).

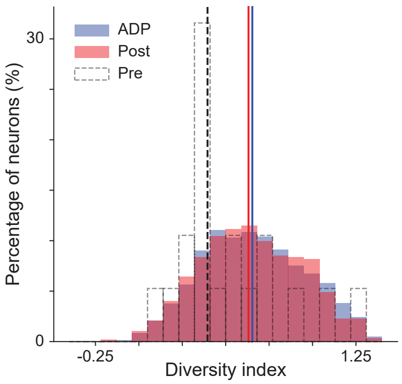


**Supplemental Fig. S25. Postsynaptic neurons and ADP controls had similar diversity indices.** Histogram of diversity index of presynaptic, postsynaptic, and ADP controls. Diversity indices were similar between postsynaptic and ADP neurons ($P = 0.4$, two-sided Welch's $t$-test with 869.0 d.f.). The postsynaptic and ADP control groups were formed by pooling data across all presynaptic neurons. Data were shown from 19 presynaptic, 570 postsynaptic, and 2,486 ADP neurons, resulting in 706 connected pairs and 18,162 ADP controls.
